# Spatially resolved dendritic integration: Towards a functional classification of neurons

**DOI:** 10.1101/657403

**Authors:** Christoph Kirch, Leonardo L Gollo

## Abstract

The vast tree-like dendritic structure of neurons allows them to receive and integrate input from many neurons. A wide variety of neuronal morphologies exist, however, their role in dendritic integration, and how it shapes the response of the neuron, is not yet fully understood. Here, we study the evolution and interactions of dendritic spikes in excitable neurons with complex real branch structures. We focus on dozens of digitally reconstructed illustrative neurons from the online repository NeuroMorpho.org, which contains over 100,000 neurons. Yet, our methods can be promptly extended to any other neuron. This approach allows us to estimate and map specific and heterogeneous patterns of activity observed across extensive dendritic trees with thousands of compartments. We propose a classification of neurons based on the location of the soma (centrality) and the number of branches connected to the soma. These are key topological factors in determining the neuron’s energy consumption, firing rate, and the dynamic range, which quantifies the range in synaptic input rate that can be reliably encoded by the neuron’s firing rate. Moreover, we find that bifurcations, the structural building blocks of complex dendrites, play a major role in increasing the dynamic range of neurons. Our results provide a better understanding of the effects of neuronal morphology in the diversity of neuronal dynamics and function.

## Introduction

Neurons are specialized excitable cells that are characterized by distinctive and often complex structures. Although the dendritic complexity is evident in various neuron types, it is often disregarded in computational models, and its role in dendritic integration in the presence of naturalistic stimuli is largely unknown. Moreover, the topology of dendritic trees may reflect fundamental elements for dendritic computation.

Each neuron in the brain is unique, and they can be classified into a myriad of neuron types and subtypes [1]. With the ever-growing NeuroMorpho.org [2–4] public online repository, there are nearly 100,000 digital reconstructions of neuronal morphology available. These data can be used for anatomically realistic models [5–7] and morphometric analyses [8]. They can be invaluable for neuronal classification, which is typically based on morphology, electrophysiology, molecular, or functional properties [9–11]. By obeying fundamental consistence rules such as enforcing neurons to have a tree topology (characterized by the absence of loops), digital reconstructions provide an unprecedented wealth of data with exquisite spatial resolution.

In stark contrast to what some simple and influential punctual models suggest, such as the leaky integrate-and-fire model [12], neuronal dendrites are not passive media that filters electric input by leaking part of the current and propagate the rest. Instead, dendrites are capable of producing supralinear amplification called dendritic spikes that occur owing to the voltage-gated dynamics of ion channels [13–15]. These nonlinear properties and the interactions between neighboring compartments regulate the transmission of information along dendrites. Hence, neuronal integration of input from various synaptic sources depends on these non-linear (non-additive) dynamics taking place at dendrites with complex topology. Much work has been done to investigate how topology determines the capability of single neurons to detect intensity of stimulus [16], to reliably detect dendritic spikes [17], to discriminate input patterns [18], and to perform other forms of dendritic computation [19–21]. There has been some attempts to study this problem analytically [22, 23]. However, given the complexity of the task, they are usually limited to regular or oversimplified dendritic structure [22].

Simple models play a major role at revealing fundamental dynamic mechanisms in neuroscience [24]. Here we describe the spatial structure of neurons and focus on main dynamics taking place at dendrites [25]. Conventional multi-compartment models often have less than 100 compartments (e.g. [26–28]) and overlook the large number of synapses (about 10,000 in a human neuron) that lead to complex nonlinear interactions. Some approaches feature a detailed description of a specific neuron. However, a major limitation of this realistic approach is the large number of parameters in the model [21, 29–34]. Many of these parameters represent unknown variables, which is typical from such high-dimensional problems that include the description of dynamics of a variety of (often spatially dependent) ion channels along the dendritic tree.

Utilizing simplified neuronal dynamics, we mapped the heterogeneous response of dendritic compartments, independently subjected to stochastic excitatory input, in digitally reconstructed neurons. The structure of these neurons are trees that can be considered as complex networks, and consist of up to 10,000 compartments. We focused on the input-output response curve of these neurons as the intensity of incoming stimuli varies over several orders of magnitude. These response curves can be used to quantify features of the neuronal dynamics [23, 35]. Fundamental principles of neuronal functions may be determined by these dynamic features. Knowing the neuronal functions may be helpful as a way to classify and compare neurons and neuron types [36] within and across species [37].

By applying this theoretical framework, our main aim is to investigate the consequences of complex dendritic structure on dendritic integration and neuronal activity. We characterize the effects of topological properties of the neurons on the dynamic range of the response functions, which quantifies the ability of neurons to discriminate the intensity of incoming input, and identify how bifurcations contribute to heterogeneous activity. Finally, we show how these findings can provide a functional classification of neurons.

## Methods

Given the large number of compartments and bifurcations that make up the dendritic arbor, any attempts of analytically modelling the propagation and interaction of potentially hundreds of spikes simultaneously are rendered nearly impractical. Further, we expect these interactions to be highly non-linear owing to the heterogeneity of the neuronal topology. We overcome this complexity in spike dynamics by adapting a discrete computational model from previous studies [16, 23, 38]. Our model preserves the main features of excitable systems, and by implementing real dendritic structures we focus on the resultant spatial properties of neurons with active dendrites.

### Digital reconstructions

NeuroMorpho is a free online database of tens of thousands of three-dimensional neuron reconstructions. Each neuron has up to thousands of individual compartments, and the dataset is available in a standardized format, allowing the development of frameworks that can implement any neuron. In contrast to previous studies [16, 23, 38], we take the entire spatial information provided by NeuroMorpho and treat the compartments as fundamental units of the neuron that are governed by identical dynamical rules (see next section). The list of neurons used in this paper is given in Table 1 (see below for a definition of the centrality). The NeuroMorpho version was 7.6.

**Table 1:**
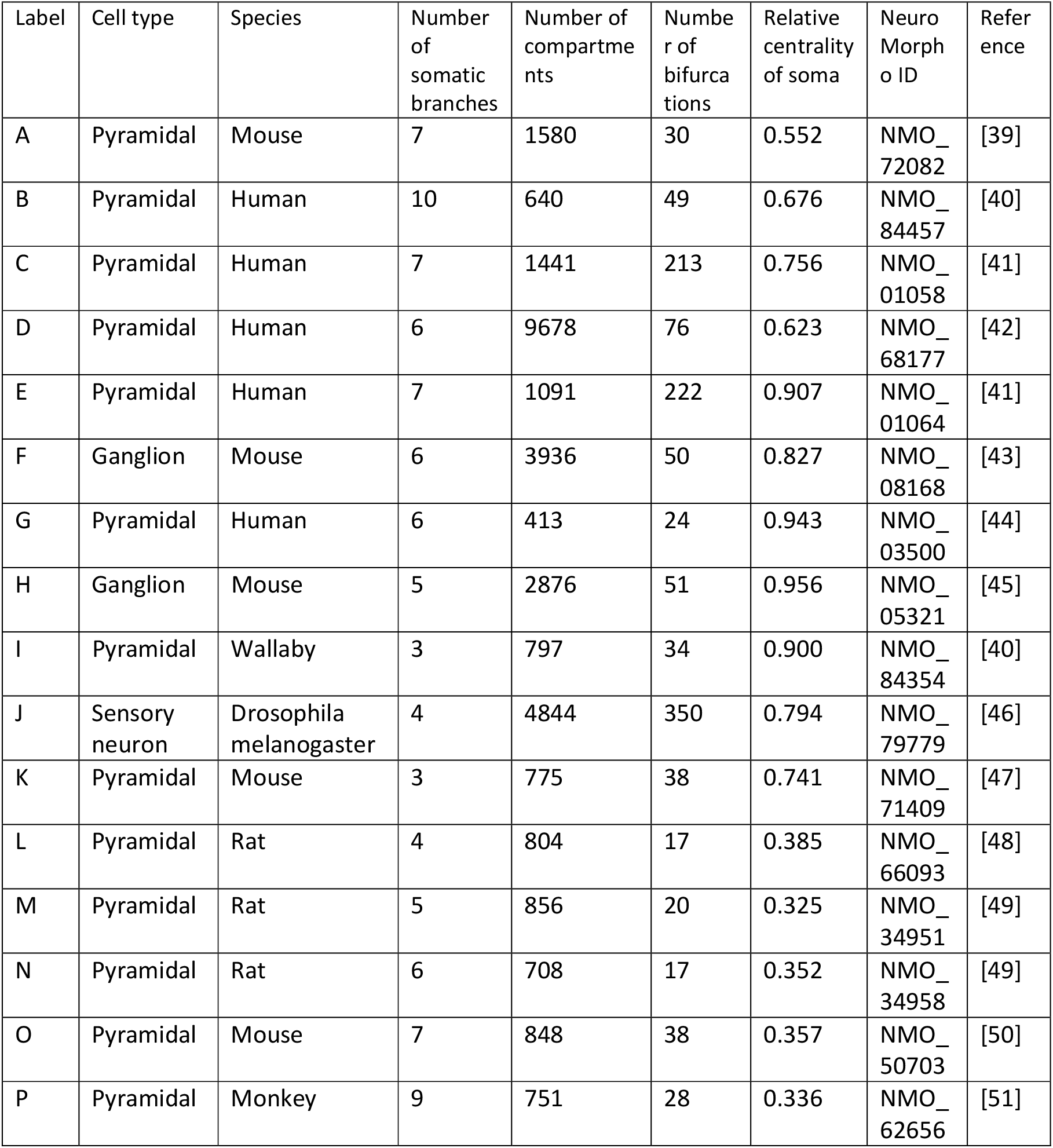
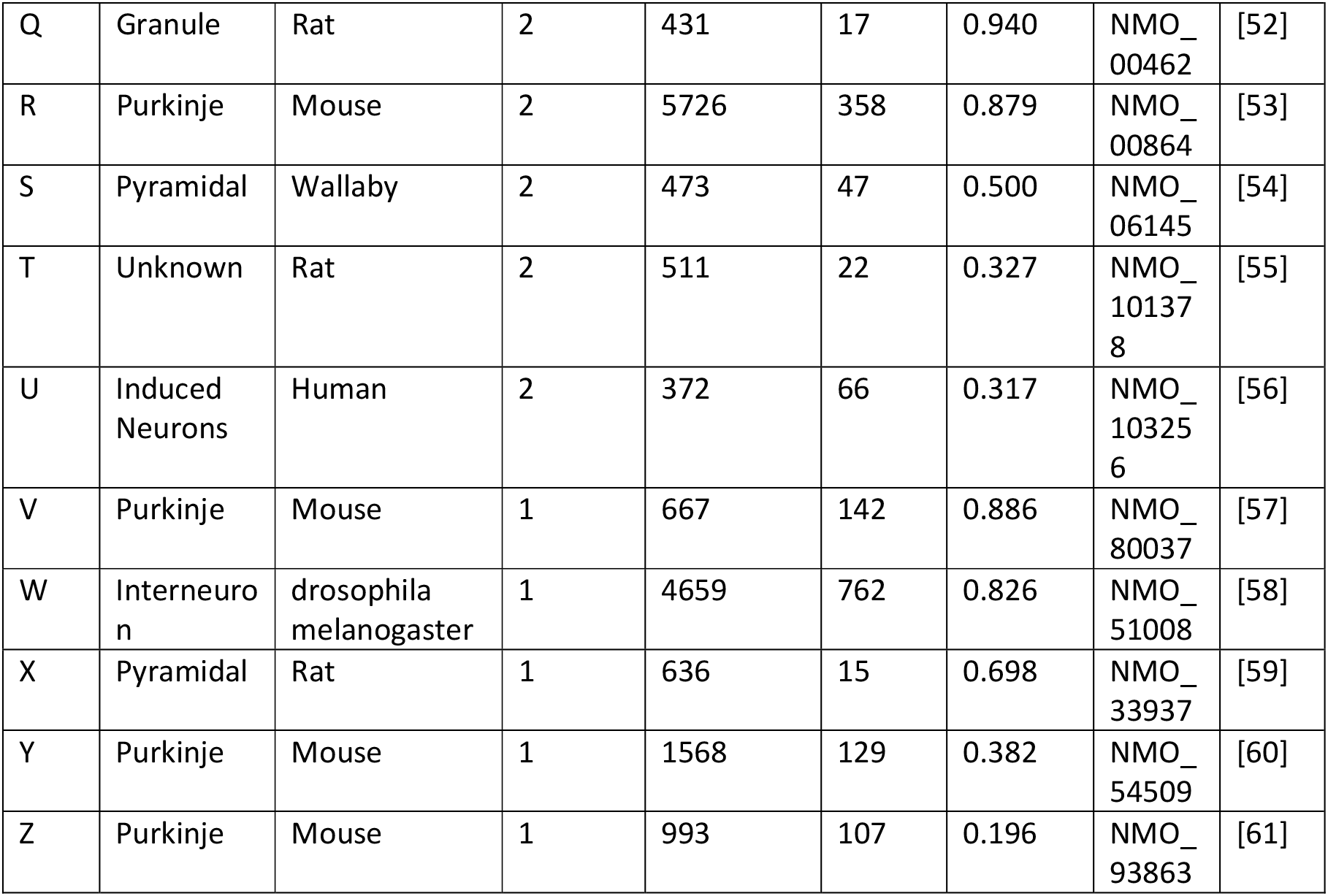
List of neuron reconstructions used in this study, taken from NeuroMorpho.Org (version 7.6).

| Label | Cell type | Species | Number of somatic branches | Number of compartments | Number of bifurcations | Relative centrality of soma | Neuro Morpho ID | Reference |
| --- | --- | --- | --- | --- | --- | --- | --- | --- |
| A | Pyramidal | Mouse | 7 | 1580 | 30 | 0.552 | NMO_72082 | [39] |
| B | Pyramidal | Human | 10 | 640 | 49 | 0.676 | NMO_84457 | [40] |
| C | Pyramidal | Human | 7 | 1441 | 213 | 0.756 | NMO_01058 | [41] |
| D | Pyramidal | Human | 6 | 9678 | 76 | 0.623 | NMO_68177 | [42] |
| E | Pyramidal | Human | 7 | 1091 | 222 | 0.907 | NMO_01064 | [41] |
| F | Ganglion | Mouse | 6 | 3936 | 50 | 0.827 | NMO_08168 | [43] |
| G | Pyramidal | Human | 6 | 413 | 24 | 0.943 | NMO_03500 | [44] |
| H | Ganglion | Mouse | 5 | 2876 | 51 | 0.956 | NMO_05321 | [45] |
| I | Pyramidal | Wallaby | 3 | 797 | 34 | 0.900 | NMO_84354 | [40] |
| J | Sensory neuron | Drosophila melanogaster | 4 | 4844 | 350 | 0.794 | NMO_79779 | [46] |
| K | Pyramidal | Mouse | 3 | 775 | 38 | 0.741 | NMO_71409 | [47] |
| L | Pyramidal | Rat | 4 | 804 | 17 | 0.385 | NMO_66093 | [48] |
| M | Pyramidal | Rat | 5 | 856 | 20 | 0.325 | NMO_34951 | [49] |
| N | Pyramidal | Rat | 6 | 708 | 17 | 0.352 | NMO_34958 | [49] |
| O | Pyramidal | Mouse | 7 | 848 | 38 | 0.357 | NMO_50703 | [50] |
| P | Pyramidal | Monkey | 9 | 751 | 28 | 0.336 | NMO_62656 | [51] |
| Q | Granule | Rat | 2 | 431 | 17 | 0.940 | NMO_00462 | [52] |
| R | Purkinje | Mouse | 2 | 5726 | 358 | 0.879 | NMO_00864 | [53] |
| S | Pyramidal | Wallaby | 2 | 473 | 47 | 0.500 | NMO_06145 | [54] |
| T | Unknown | Rat | 2 | 511 | 22 | 0.327 | NMO_101378 | [55] |
| U | Induced Neurons | Human | 2 | 372 | 66 | 0.317 | NMO_103256 | [56] |
| V | Purkinje | Mouse | 1 | 667 | 142 | 0.886 | NMO_80037 | [57] |
| W | Interneuron | drosophila melanogaster | 1 | 4659 | 762 | 0.826 | NMO_51008 | [58] |
| X | Pyramidal | Rat | 1 | 636 | 15 | 0.698 | NMO_33937 | [59] |
| Y | Purkinje | Mouse | 1 | 1568 | 129 | 0.382 | NMO_54509 | [60] |
| Z | Purkinje | Mouse | 1 | 993 | 107 | 0.196 | NMO_93863 | [61] |

### Compartment dynamics

We adapt a synchronous susceptible – infected (active) – refractory – susceptive model (SIRS) used in previous studies [16]. A compartment switches states as a result of interactions from its neighboring compartments, or stochastic processes. A compartment that is in the susceptible state (state *0*) will remain there until activated either externally via synapses (see below), or by a propagation of activity from an active neighbor. A signal propagates to a susceptible neighboring compartment with a constant probability *P*. Once active (state *1*), a compartment will switch to the refractory period (state *2*) for a specific time, after which it will return to state *0*. Here, we fix the refractory period to 8 time steps. Because of these dynamic rules, activity may spread to all susceptible neighboring compartment and travel in various directions (see Video S1). Moreover, two opposing signals will not add but annihilate each other [62]. The model also correctly recreates backpropagation, in which an action potential will travel back up the dendritic arbor once the soma has been activated [63]. The soma itself also follows the same rules, however, it remains distinctive because it may be connected to many branches (Table 1).

In reality, many external factors are responsible for determining whether and when a compartment should fire. For example, a compartment could have thousands of synaptic connections, some inhibitory, some excitatory. In the end, however, the result will be either On (activate the compartment) or Off (remain in the susceptible state). We model this result using a probabilistic approach, where the probability of an excitatory synaptic signal is *r* = 1 - exp(−*h*∙*δt*), where *h* is the excitation rate, and *δt* is the time step of 1ms. Here we will focus on a range of *h* that spans several orders of magnitude (from 10^−4^ to 10^4^ Hz), and *P* that varies from 0.5 to 1 (as values of *P* smaller than 0.5 exhibit a very strong attenuation, which is not plausible and give rise to little spatial contribution). We simulate each neuron for 10^6^ time steps (1000 seconds) for combinations of *h* and *P*, running every simulation five times. Over each simulation, we count the total number of times the soma fires (*F*_*S*_) and the total number of times a dendritic compartment fires (*F*_*D*_).

### Firing rate

The characteristic sigmoidal response function of a neuron is recovered by plotting the firing rate at some compartment against the excitation rate *h* for some value of *P*. It shows how the neuron responds to various levels of input activity. An important feature of the response curve is the dynamic range, which represents the range of input rates that the neuron can effectively discern. By convention [35, 64], it is defined as:

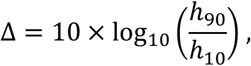

where *h*_10_ and *h*_90_ correspond to the excitation rates that produce a firing rate that is 10% and 90% of the maximum firing rate.

### Relative energy consumption

Another metric of determining performance is the relative energy consumption, which we define here as

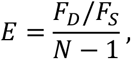

where *N* is the total number of compartments, and *F*_*D*_ and *F*_*S*_ are the number of dendritic and somatic spikes, respectively. The energy indicates how active the whole dendritic tree is compared to the soma, that is, how many times, on average, dendritic compartments activate for each somatic spike.

### Dynamics of benchmark neurites

To better understand the relationship between neuronal morphology and dynamics, we constructed and simulated a set of artificial neurites, which consist only of two branches. The primary branch is taken to be of fixed length *N*, whereas the secondary branch changes both in length *L* and the position *Q* at which it bifurcates from the primary branch, where *Q* is the index of the parent compartment of the primary branch. These toy neurons can be used to isolate the behavior we see in the full neurons.

### Centrality

A way of quantifying the soma’s position in the neuron is by estimating how far away it is from the furthest endpoint. Let *T* be the number of dendritic endpoints (branch terminals) of the neuron, and *D*_*ij*_ be the distance (number of compartments) along the neuron from the *i*-th compartment to the *j*-th terminal compartment. Since no loops exist, this distance is always unique. Then we define the centrality of the *i*-th compartment as

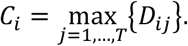

Once the set of all compartmental centralities *C* = { *C*_1_, …, *C*_*N*_} has been calculated (excluding axonal compartments), the relative centrality of the soma is given by

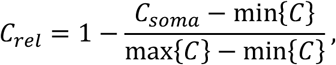

where *C*_*rel*_ = 0 implies that the soma is the least central compartment of the neuron, while *C*_*rel*_ = 1 implies that the soma is the most central compartment.

## Results

We investigate how morphology affects the dynamics of neurons. To focus on the topological properties of neurons with many compartments (100-10000), we utilized a simple dynamic model: a canonical cyclic cellular automata model [16, 35, 65]. We first introduce, illustrate, and characterize the relationship between neuronal structure and dynamics in a pyramidal mouse neuron [39]. Then, we compare the resulting dynamical properties of multiple neurons from different species with a variety of neuronal cell types and structures (see Methods). Each compartment is considered to have synapses that can receive external input, and become active at a rate *h*, which is varied over several orders of magnitude. The activity propagates to quiescent (susceptible) neighbors with a probability *P*. By keeping track of the somatic and compartmental firing events, we quantify some fundamental dynamical properties of the neuron, such as the firing rate, relative energy consumption, and dynamic range. The emerging dynamics of the model is depicted in Fig. 1 (see also Video S1).

**Figure 1.**
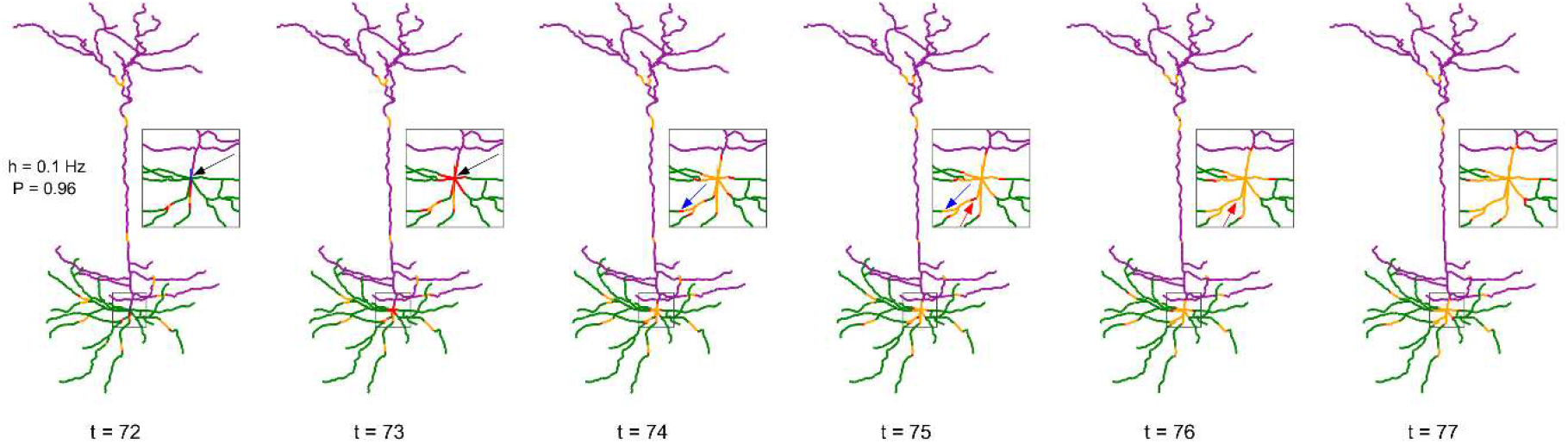
Series of snapshots showing how signals propagate along the neuron. Visualization of digital reconstruction of neuron A (see Methods), where compartments are colored according to the following scheme: green for basal dendrites, purple for apical dendrites, blue for the soma (too small to be seen here), red for active (spiking) compartments, orange for refractory compartments. In each time step, spikes may propagate from active compartments to susceptible neighboring compartments with transmission probability *P* (here, *P* = 0.96). A susceptible compartment may also spike due to the synaptic input, which we model stochastically with a Poisson rate of *h* (here, *h* = 0.1 Hz). Once active, a compartment transitions to the refractory period and is unable to spike for 8 time steps. The inset plot provides a closer look at the soma and surrounding compartments. At *t* = 73, the neuron fires as a result of the dendritic integration of synaptic input (somatic spike, see black arrow). Between *t* = 74 and *t* = 75, a signal fails to transmit (see blue arrow). Between *t* = 75 and *t* = 76, two spikes can be seen annihilating each other (see red arrow). The snapshots were taken from Video S1.

### Spatial maps of the activation rate

One strength of our computational model is the ability to simulate the complex dynamics of very large neurons with many compartments and including their specific digitally reconstructed morphology. To highlight the heterogeneity across dendritic branches, these results can be visualized spatially in the form of heat maps. The average rate of activation across time (*T* = 10^6^ ms) of each compartment (*N* = 1580) is illustrated in Fig. 2 for different values of *P*, the probability of propagation of activity (see Fig. S1 for other neurons). It is clear that the topology of the neuron affects the firing rate. Crucially, the soma (indicated by the arrow, left panel) becomes active at a higher rate than other compartments. Please note the different colormaps for the different panels. Because the model assumes homogeneity across compartments (*P* and *h*), this amplification of firing rate at the soma occurs solely due to the topology. The soma has seven branches connected to it. These branches increase the likelihood of activations to reach the soma in comparison to other compartments because they can come from any active neighbor. Moreover, the lower *h* and *P* are, the greater is the relative amplification of the firing rate at the soma (see Fig. S2 for other neurons and values of *h*). These results also lead to the prediction that the activation rate can vary substantially across dendritic sites. For neuron A (Fig. 2), the firing rate increases near the soma, but the firing rate is usually larger in sites close to a large number of bifurcations (see Figs. S1 and S2).

**Figure 2.**
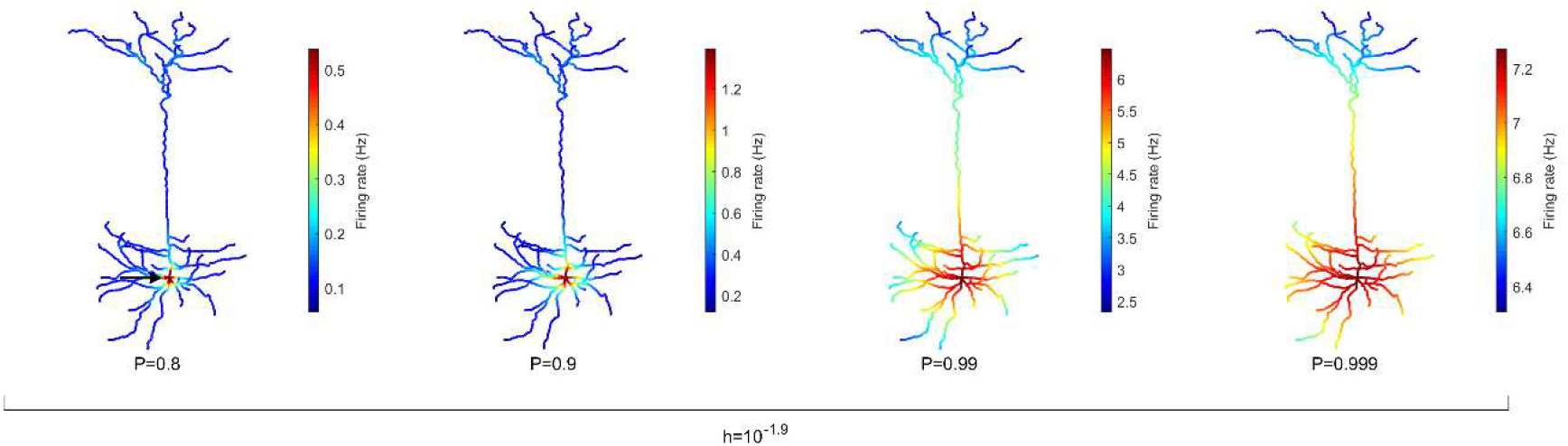
Spatial map of the compartmental firing rate. The soma (marked by the arrow) lies among the most active region of the neuron, regardless of the signal transmission probability *P*. The amplification of the activity at the soma is stronger for lower values of *P*. Refer to Figures S1 and S2 for heat maps of other neurons. Note different color scales.

### Energy consumption for neuronal spikes

Active dendrites are capable of generating dendritic spikes, which allows for enhancement of activity and non-linear interactions. However, these dynamical benefits require additional energy. Here we estimated the relative energy consumption of the soma with respect to the number of activations the whole neuron experiences (see Methods for details). In our model, two processes are responsible for increasing the relative energy consumption of the neuron: (i) the external driving *h*, and (ii) the propagation and bifurcation of a dendritic spike as it passes a dendritic junction, initiating an additional spike that consumes energy. On the other hand, there are three mechanisms that reduce the relative energy consumption of the neuron: (i) energy dissipates stochastically due to the attenuation rate (*1-P*), (ii) signals can propagate in nonlinear waves that can be annihilated, and (iii) a signal travelling away from the soma will necessarily die when reaching an endpoint of a branch.

It is important for a neuron to balance its energy usage while performing its functions. In Fig. 3, we plot the relative energy consumption over the parameter space. The blue regions highlight areas of most efficient operation, as somatic spikes require fewer dendritic spikes to take place. In this case, it corresponds to a weak external driving rate (*h* < 0.01 Hz) and a rate of transmission that allows for failure of dendritic spike propagation (*P* < 0.95).

**Figure 3.**
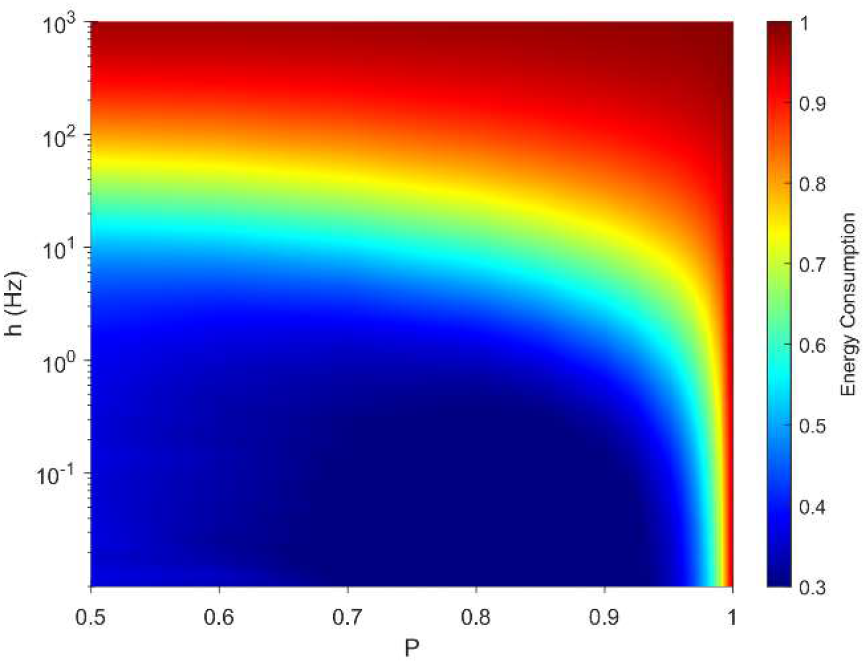
Variations in relative energy consumption of the neuron over the parameter space. The relative energy consumption of a neuron (demonstrated here using neuron A, see Methods) is a measure of how often, on average, dendritic compartments spike per somatic spike. Here the energy consumption grows for high external driving and towards deterministic propagation (*P* = 1).

The behavior of our measure of energy consumption in the limits of the parameter space, as seen in Fig. 3, can be explained. Firstly, as *P* approaches 1, every signal will be able to visit all compartments of the neuron once, or interact with another signal which will be able to visit the remaining compartments. In that case, the average firing rate of the compartments is identical and does not depend on topology. Hence, every dendritic compartment has to fire once for the soma to fire once (deterministic behavior). Secondly, in line with other studies [66], the energy consumption of spikes increases with *h*. Moreover, as *h* approaches ∞, every compartment will fire independently at the highest possible rate,

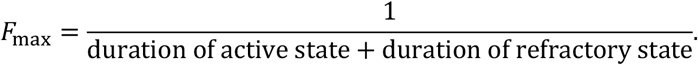

Again, the average firing rate of the compartments is identical, hence we expect a relative energy consumption of 1. Only at these simple limiting cases is the dynamical behavior independent of morphology.

### Spatially resolved response function and dynamic range

One informative and influential way to quantify how dendritic trees process incoming signals is given by input-output response functions. It is defined by the mean output activation rate (across a long time interval, here *T* = 10^6^ ms) as a function of the rate of activations induced by external driving *h* (neuronal input). This means that the firing rate can be computed for each compartment. Response functions have their minimal in the absence of external input and their maximal for very strong external driving. As illustrated for different recording sites, response functions exhibit a sigmoidal shape (Fig. 4). These results show for this neuron a larger firing rate at the soma compared to other regions, especially at low external driving, which is consistent with the amplification of the firing rate observed at the soma (Fig. 2). At high values of *h*, the saturation of the response curves occurs in a similar manner regardless of the recording site. Moreover, for *P* ≈ 1, this spatial heterogeneity vanishes.

**Figure 4.**
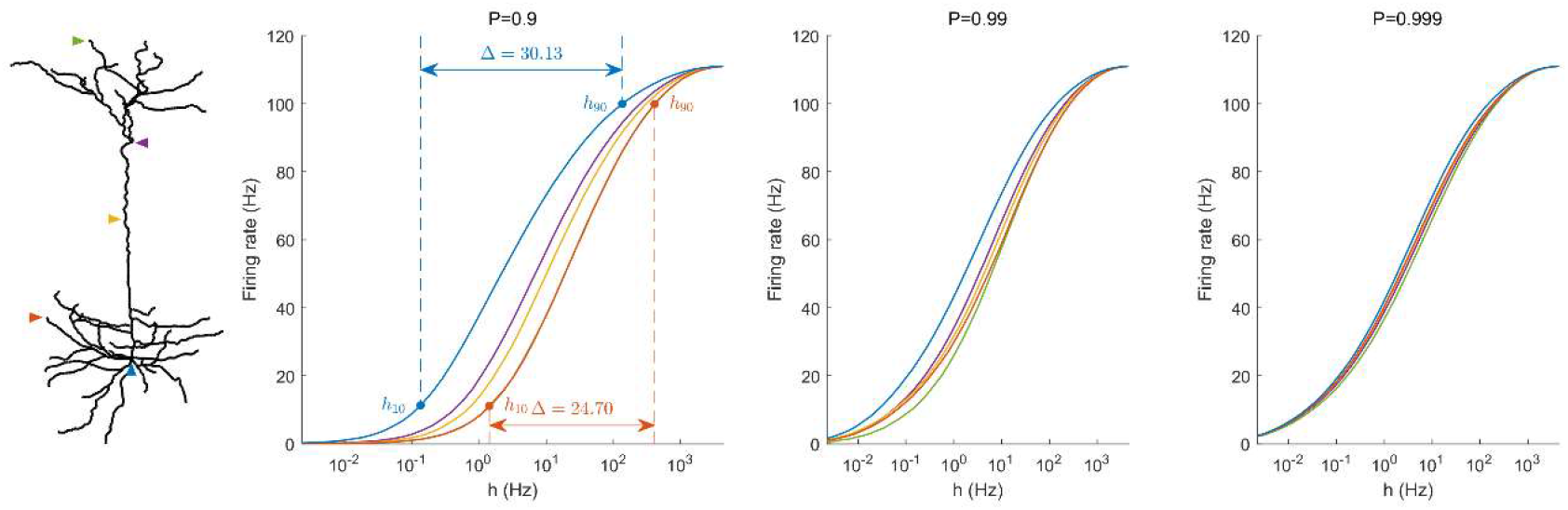
Response functions at representative compartments. For each of the marked compartments, we plot the firing rate against the synaptic input rate *h*. The dynamic range Δ is a function of *h*_*10*_ and *h*_*90*_ (see Methods). Lower values of *P* yield stronger spatial dependence in the response functions of compartments.

An important feature of response functions that can be quantified corresponds to the dynamic range Δ [16, 35]. It depends on the values of external driving at which the neuron responds at 10% of its maximal firing rate (*h*_10_), and at 90% (*h*_90_). The dynamic range quantifies the range between *h*_10_ and *h*_90_ (see Methods for details). It assumes that, based on the firing rate, the neuron is unable to reliably distinguish activation rates too close to saturation, *h* < *h*_10_ and *h* > *h*_90_. Figure 4 also illustrates the definition of the dynamic range for the response function measured at the soma and at a basal dendrite.

The dynamic range is a measure of the sensitivity to changes in the input rate. A large dynamic range indicates that a neuron can discern signals produced by a large range of input rates. For example, ganglion cells from the retina require a large dynamic range to be able to reliably respond to changes in lighting conditions that vary over several orders of magnitude [67].

Our model allows us to identify the spatial gradients observed in the dynamic range (Fig. 5). Despite changes in the transmission probability *P*, the soma tends to exhibit high values of dynamic range. However, if *P* is close enough to 1, the amplification of signals become detrimental, and the dynamic range at the soma can be lower than other regions. In this latter case, it is also relevant to notice that the differences in dynamic range across the neuron are overall very small (< 1dB). This happens because *h*_10_ remains essentially unchanged whilst the saturation of the response function (*h*_90_) occurs slightly earlier at the soma (see right panel of Fig 4).

**Figure 5.**
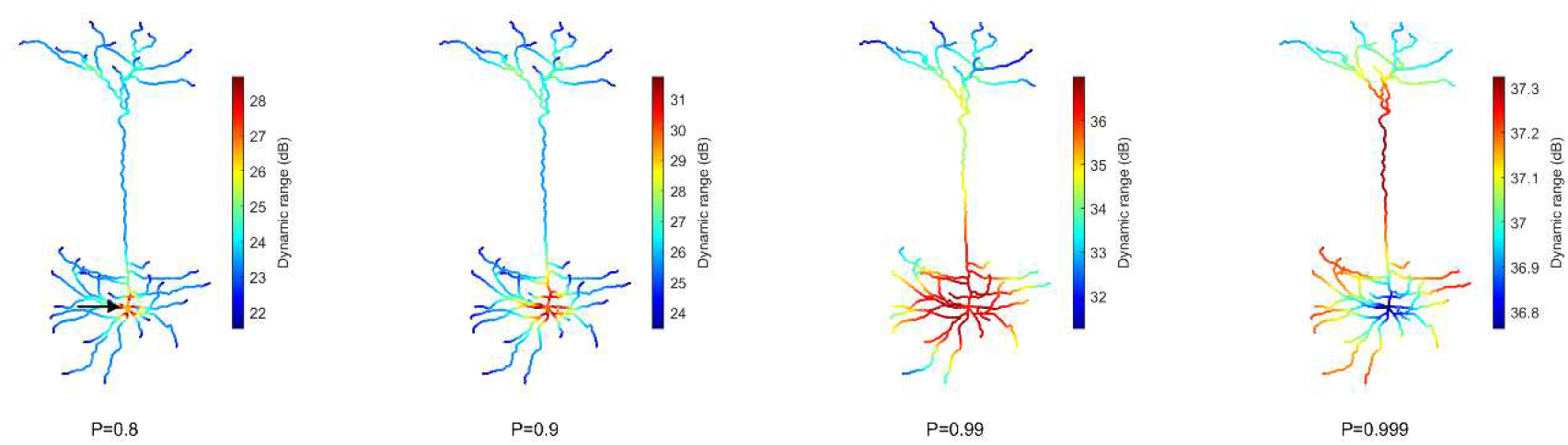
Heat maps of the spatial distribution of the dynamic range. The soma (marked by the arrow) usually exhibits the opposite performance of the extremities. The dynamic range is larger at the soma for low transmission probability P, and lower for extremely high *P*. Please note the different color bars. See Fig. S3 for other neurons.

### Teasing apart the effects of a single branch on the dynamic range

To characterize the effects of neuronal topology, we explored the dynamic range of each compartment in the neuron as a function of its distance from the soma (Fig. 6). This new perspective reveals a general relationship that is mostly governed by *P*. For *P* < 0.99, the dynamic range mostly decreases with the distance from the soma, and bifurcations generate a local boost in dynamic range, while a drop occurs at branch endpoints. For larger values of *P*, the dynamic range peaks at the main branch to the distal dendrites (yellow).

**Figure 6.**
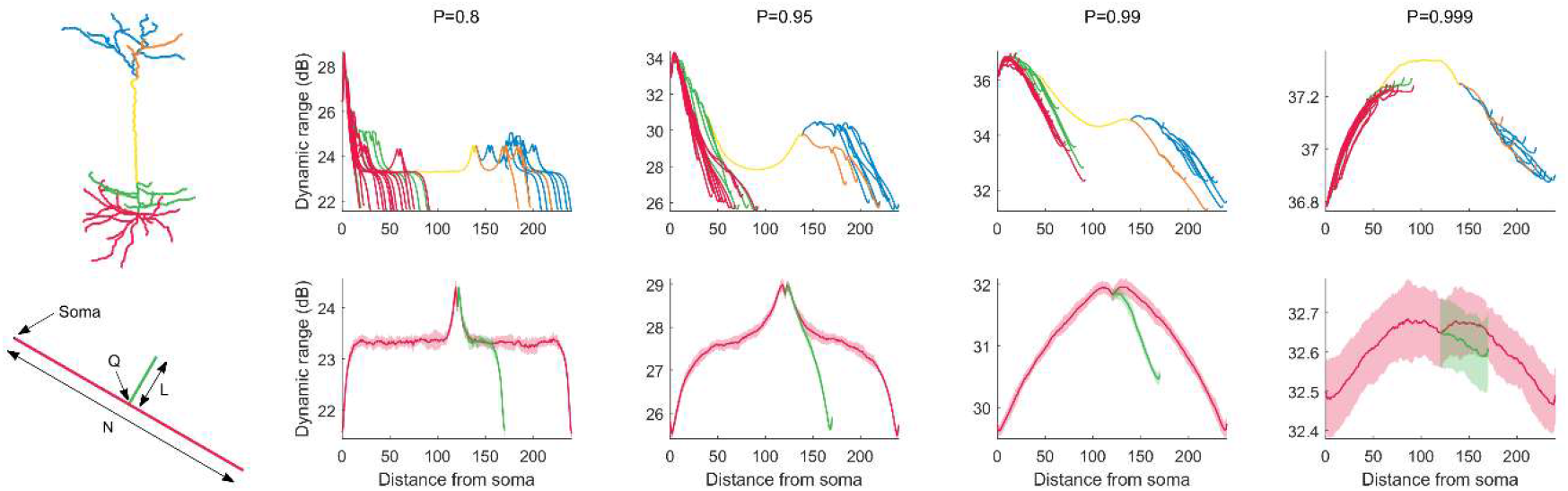
Dynamic range as a function of the distance to the soma. Top row: The functional profile depends largely on the signal transmission probability *P*. Distinct structural regions of the neuron are color coded. For other neurons (see below), refer to Fig. S4. Bottom row: Minimalistic structural model of a neuron with a main branch (red) of length *N* = 240, and a single bifurcation branch (green) of length *L* = 50. The secondary branch is connected to the primary branch at compartment index *Q* = 120. Many features in the variation of dynamic range can be exhibited by the simple toy neuron. The shaded regions are ±1 standard deviation of the mean, over 10 trials. The simple neurons were simulated for 10^5^ time steps. For additional results where we vary *N*, *L* and *Q*, refer to Fig. S5.

**Figure 7.**
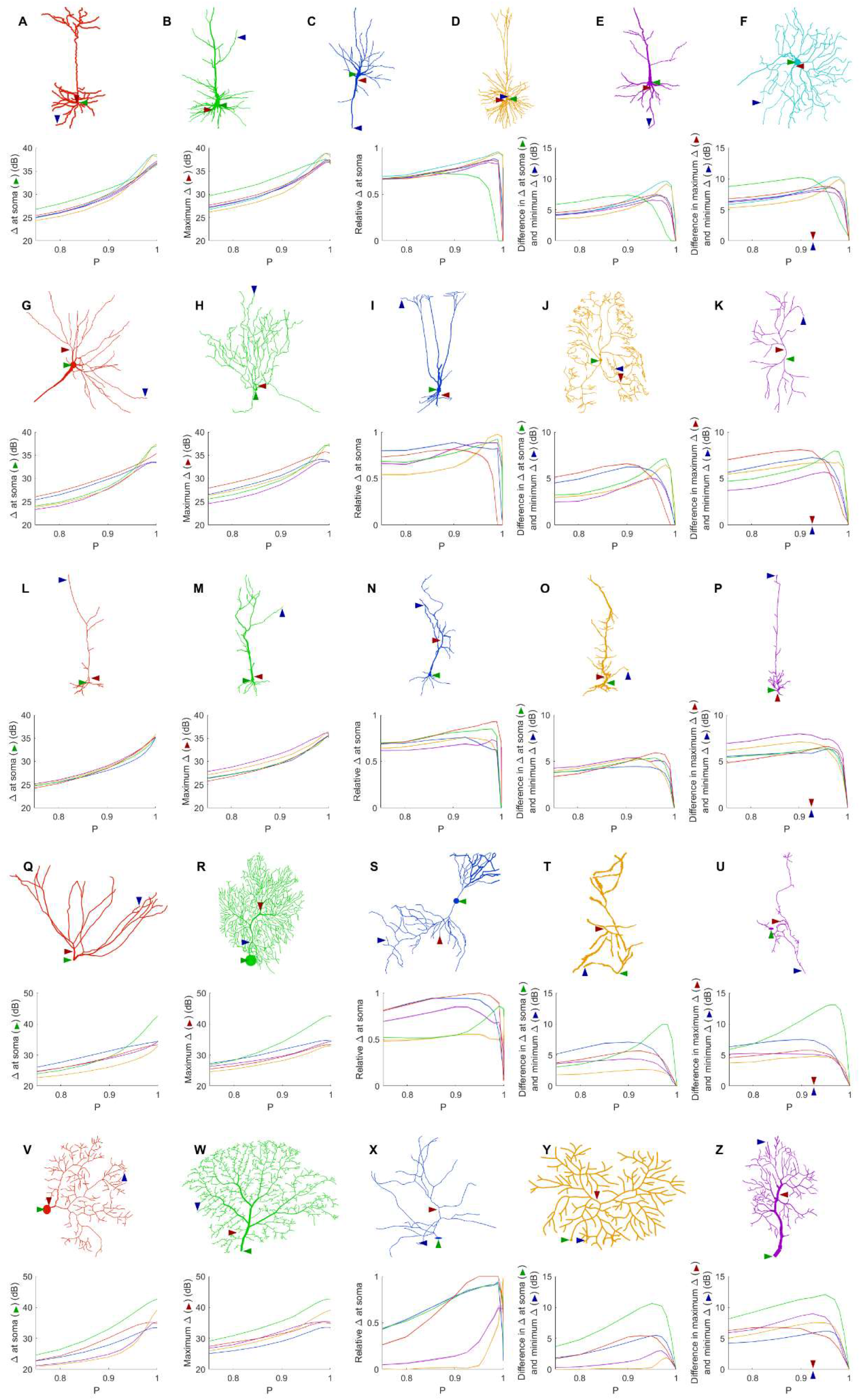
Comparison of dynamic ranges across neurons. Each row of graphs corresponds to the row of neurons above (see Methods for a description of each neuron). Colors do not have physical meaning, and are used only to differentiate the neurons of each row. The soma is marked by the green arrow, while the red and blue arrows indicate the location of the compartment at which the dynamic range is highest and lowest when *P* = 0.92. The relative dynamic range was calculated using (Δ_soma_ – Δ_min_) / (Δ_max_ – Δ_min_). See Table 1 for original references and description of neurons.

To better understand the relationship between dendritic topology and neuronal dynamics, we systematically studied how a single branch modifies the dynamic range. To pinpoint the effects of a single bifurcation on the dynamic range, we created a set of very simple neurons containing a single bifurcation with a small branch (Fig. 6). Starting with a primary branch (black) of constant length, we append a secondary branch (red) of length *L* to the primary branch at position *Q*, then run the simulations. Despite its simplified spatial structure, the minimal toy neuron faithfully reproduces many features in its dynamic range profile that we see from the full neuron reconstruction. For example, the effect of a single bifurcation or branch endpoint on the local dynamic range is consistent. A more complete set of toy neurites and their dynamic range is provided in Fig. S5.

### How does the dynamic range change across neurons?

Given the wide variety of neuronal morphologies, one might expect very different neurons to exhibit very different dynamics. In general, the dynamic range at the soma, the maximum dynamic range and the minimum dynamic range of the neuron increase with *P*. At its highest, the dynamic range at the soma attains values of more than 35 dB for all neurons, and up to 43 dB. In addition, as shown before (Figs. 4 and 5), the measures of the dynamic range at the different sites become more homogeneous for very large values of *P*. The measure of the relative Δ at the soma shows that the soma is most often close to the sites of maximum dynamic range. However, some neurons (R, T, Y and Z) exhibit the dynamic range at the soma somewhat smaller than the maximum dynamic range of the neuron. Moreover, the maximum heterogeneity of dynamic range across neurons varied substantially (from 7 to 13 dB). This spatial heterogeneity, even when the dynamics of the compartments is assumed to be identical, demonstrates that the topology of neurons plays a major role in shaping neuronal dynamics.

### Trends in energy consumption

The dynamic range tells us about the capability of neurons to encode stimuli that vary over orders of magnitude. However, this process has a cost, and the dynamic range does not reveal the neuronal efficiency in terms of energy consumption. Previously, we have introduced our measure of relative energy consumption, defined as the average number of times a dendritic compartment spikes for an action potential to be generated (Fig. 3). If we compare the energy consumption across all neurons, three distinct types of behaviors emerge (Figs. 8-10), along with a transitioning behavior (Fig. 11). For the full set of energy consumption plots, refer to Fig. S6.

**Figure 8.**
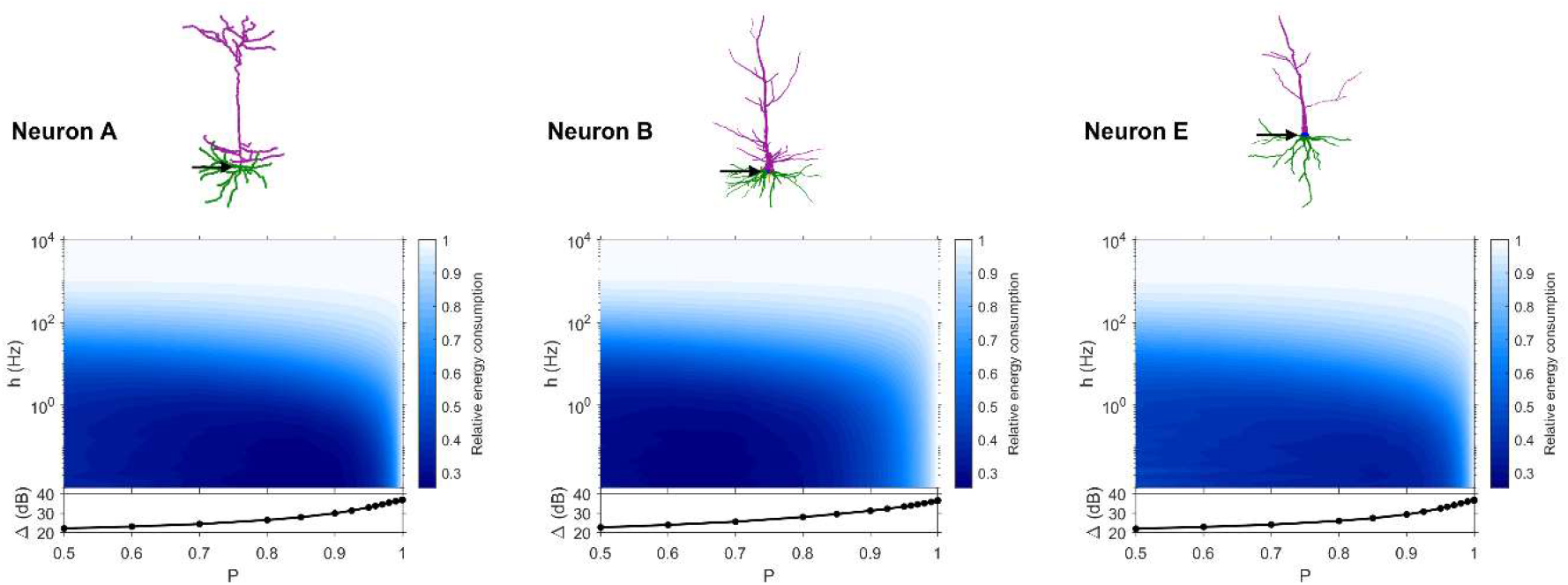
Relative energy consumption for Type 1 neurons. The location of the soma is indicated by the arrow, and its dynamic range is plotted in the bottom graph. Note the different color bars.

**Figure 9.**
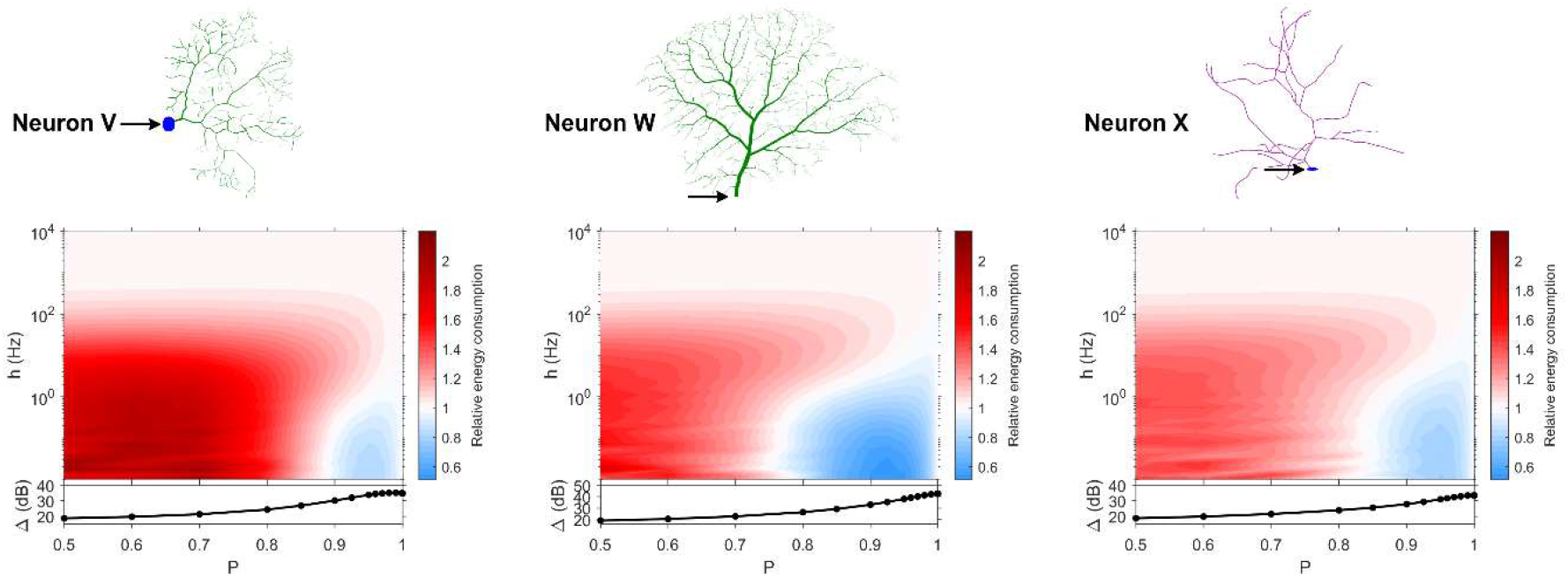
Relative energy consumption for Type 2 neurons. The location of the soma is indicated by the arrow, and its dynamic range is plotted in the bottom graph. Note the different color bars.

**Figure 10.**
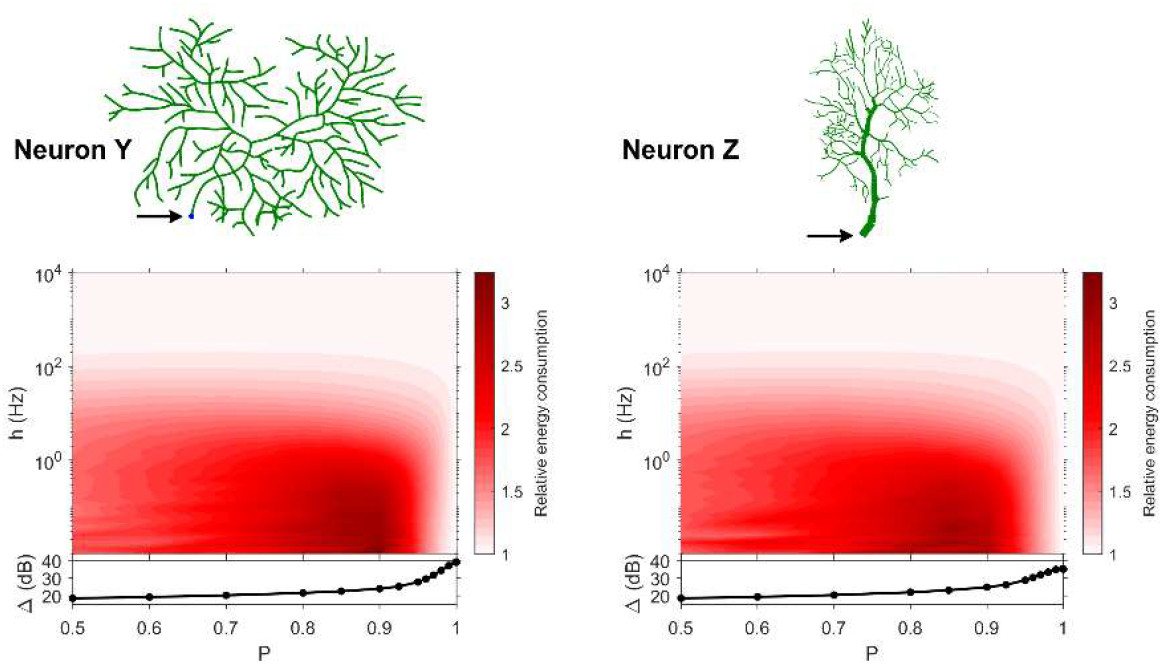
Relative energy consumption for Type 3 neurons. The location of the soma is indicated by the arrow, and its dynamic range is plotted in the bottom graph. Note the different color bars.

**Figure 11.**
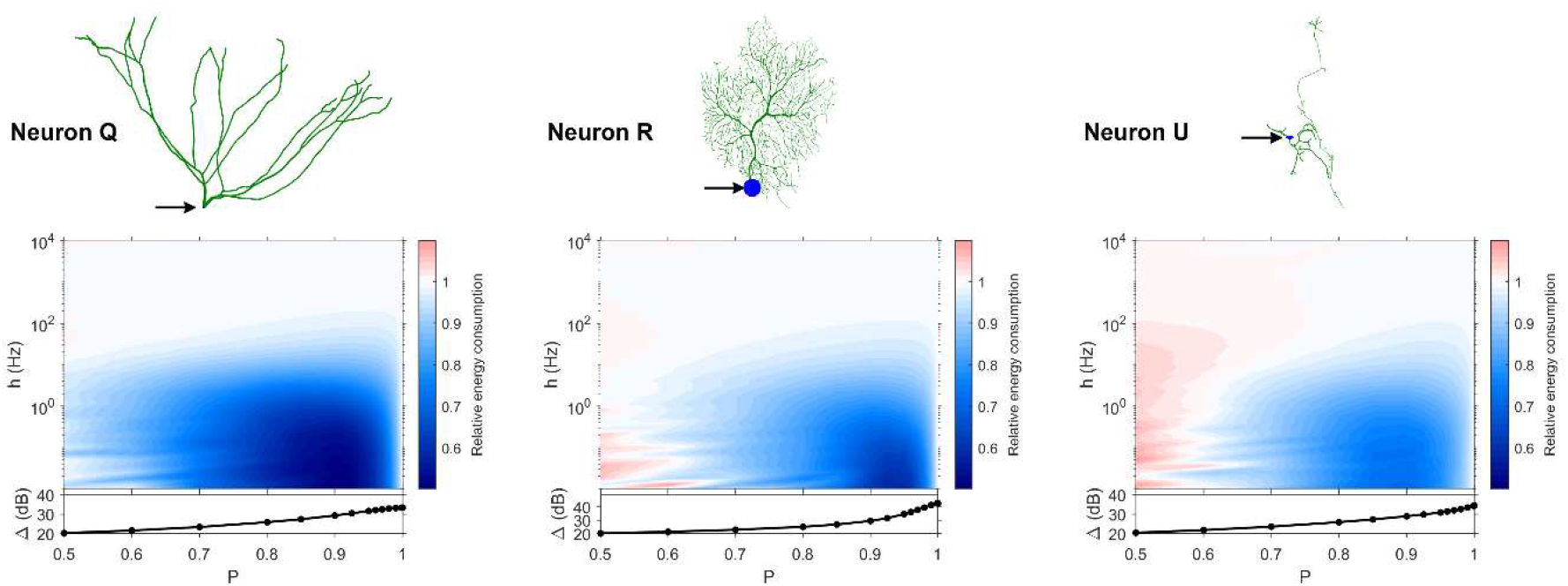
Relative energy consumption for Type T neurons (transition type). The location of the soma is indicated by the arrow, and its dynamic range is plotted in the bottom graph. Note the different color bars.

Type 1 (Fig. 8) occurs for the majority of neurons, despite the stark differences in morphologies. For these neurons, the energy is minimized for approximately *h* < 10 Hz and *P* < 0.95. Although the maximum dynamic range at the soma of every neuron occurs near *P* = 1, it corresponds to a very high relative energy consumption. However, a slight decrease in *P* can almost minimize the energy consumption in these neurons, while keeping the dynamic range near its maximum. An optimally performing neuron would therefore slightly subject signal propagation to failure, saving energy without considerable loss in dynamic range.

Type 2 (Fig. 9) corresponds to a narrower region of minimal energy consumption (*h* < 1 Hz and 0.85 < *P* < 0.95). Unlike Type 1, decreasing the transmission probability below 0.8 is detrimental to the efficient operation of these neurons.

Type 3 (Fig. 10) display high relative energy consumption, with a maximum in the region *h* < 1 Hz and 0.8 < *P* < 0.9. Moreover, the maximum and minimum energy consumption is much higher than for neurons of other types, and it never reaches a value below 1. As such, Type 3 seems to be intrinsically energy inefficient.

The transition (Fig. 11) Type T exhibits a behavior that is between Types 1 and 2. Given the different behaviors, it is clear that the dendritic morphology affects the energy consumption of neurons. A crucial element in the computation of the energy consumption corresponds to the location of the soma, and the number of branches it has. Neurons of Type 1 have the soma located in a centralized position.

### Centrality

To quantitatively describe the soma’s relative position within the overall extent of the dendritic arbor, we devise a general measure of centrality (see Methods). When *C* = 1, the soma is considered the most central compartment in the neuron. When *C* = 0, it is considered the least central. If a compartment is not central, it does not necessarily imply that it lies near the border of the neuron; for example, as for neuron A, compartments in the two separated regions of high bifurcation densities would experience a low centrality despite being surrounded by many compartments. Only the branch connecting these two areas would be central. Heat maps of the compartmental centrality for all neurons are provided in Fig. S7. The spatial mapping reveals that centrality is related to how symmetrically the rest of the neuron is distributed around a compartment.

### Categorization

Based on our estimation of energy consumption, the behavior of Type 2 can be distinguished from the behavior of Type 3 by the centrality (Fig. 12). Moreover, the behavior of Type 1 and the transition Type T can be explained by the number of branches connecting to the soma. This is an important property of neurons, as more branches naturally allow the soma to capture more information from the dendritic arbor. Then, if the soma is also located centrally, information can reach the soma more easily from any part of the neuron, and vice versa.

**Figure 12.**
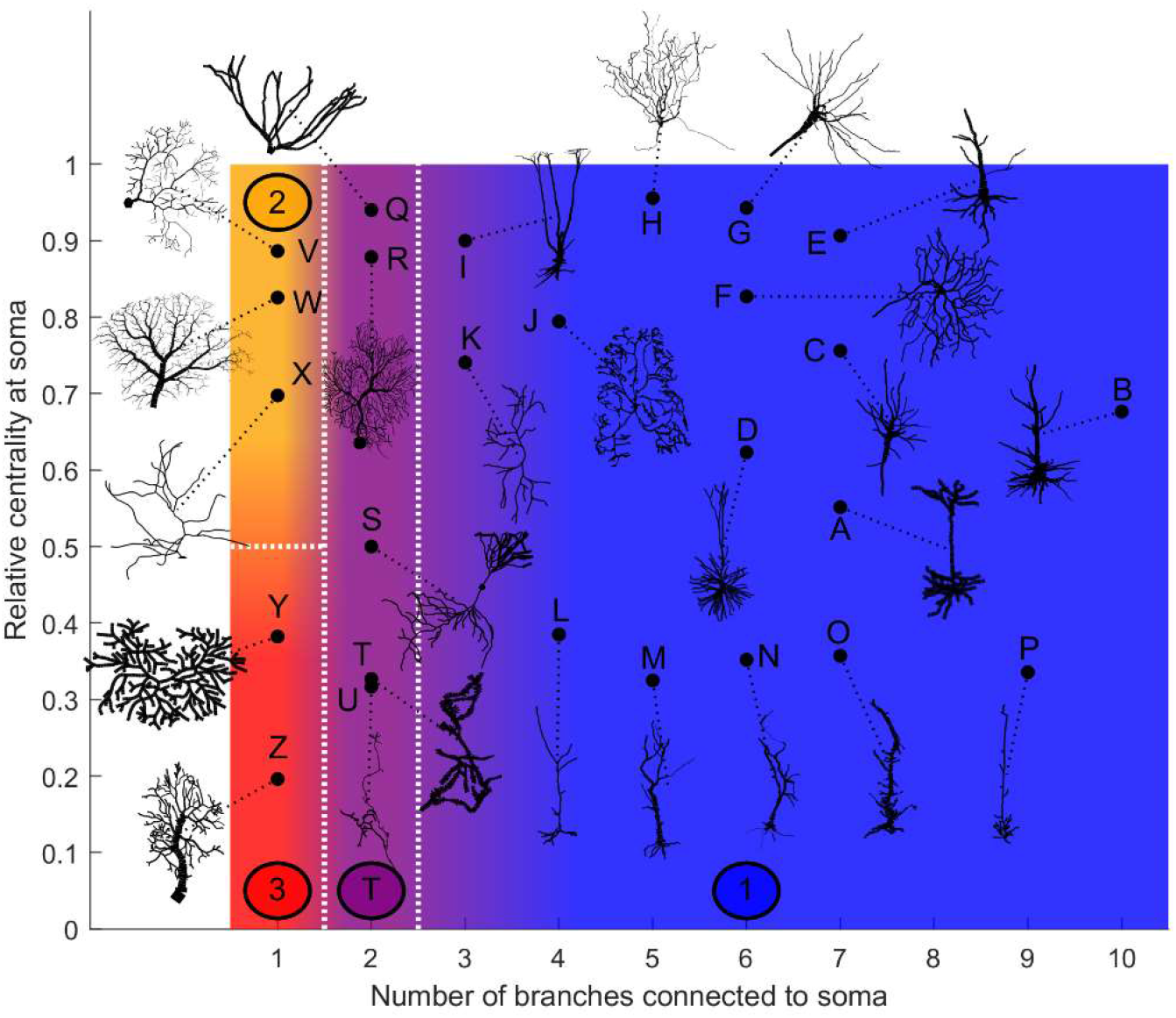
Categorization of neurons according to their relative energy consumption profile. Type 1 (blue) corresponds to Fig. 8, Type 2 (yellow) to Fig. 9, Type 3 (red) to Fig. 10, and Type T (transition, purple) to Fig. 11. The white dotted lines represent the approximate boundaries between different neuron types, and the colors indicate the different types.

Taking into account how these main structural features affect neuronal dynamics, we propose a categorization of neurons based on the relative centrality of the soma, and number of somatic branches (Fig. 12). Neurons in the same category exhibit qualitatively similar energy consumption profiles, and thus determine how efficiently the neuron operates over the parameter space: Type 1 neurons are intrinsically energy efficient, while Type 3 is intrinsically inefficient. The transition between classes is smooth. For example, neurons with 3 or 4 somatic branches have a higher minimum energy consumption than those with more somatic branches (see Fig. S6), however, they follow the same general behavior.

The general neuronal firing behavior of neurons can also be ascribed to their classification. For *P* = 1, the average response functions of neurons within types are very similar, as they do not depend much on the centrality and number of somatic branches. However, as *P* decreases, each class tends to behave independently (Fig. 13 a-c). For a given input rate, neurons in category 1 fire more often, followed by the neurons in the transition regime, neurons of category 2 and neurons of category 3. Furthermore, we find that a low probability of signal propagation has a detrimental effect on the somatic dynamic range of neurons with a low number of somatic branches (Fig. 13 d). Additionally, neurons with a large number of compartments or bifurcations can achieve a higher maximum dynamic range independently of other morphological features (Fig. 13. e, f). How close this site with maximum dynamic range is to the soma, however, does depend on the neuronal morphology.

**Figure 13.**
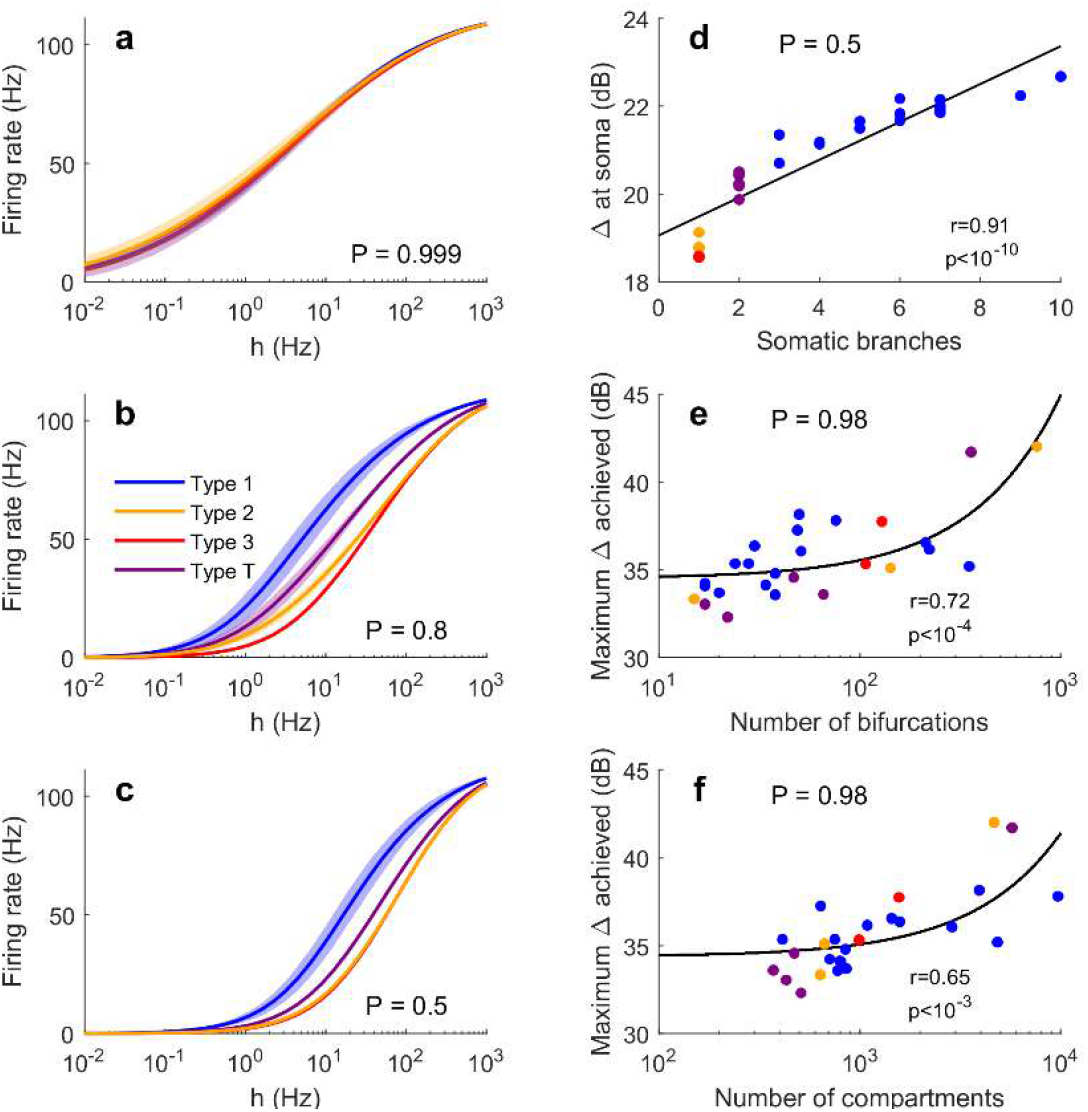
Comparison of firing behavior. (a-c) Average somatic response functions of the neurons in each category (see Fig. 12) for different values of *P*. The shading represents the standard deviation of firing rate within each group for a given stimulus intensity. For large *P*, all groups converge to the same behavior. For lower *P*, there is a distinct behavior between the average response functions of each group. (d) Dynamic range at the soma of each neuron against the number of somatic branches for *P* = 0.5. The colors represent the groups. The black trend line and Pearson correlation (*r* = 0.91) confirm that the dynamic range is strongly correlated to the number of somatic branches for lower values of *P*. (e, f) At high values of P, the number of bifurcations and number of compartments are good indicators of the maximum dynamic range that a neuron can achieve (here P = 0.98). Please note the logarithmic scale. The black trend lines and Pearson correlation show that the maximum Δ achieved is approximately linearly correlated with the neuron’s size and complexity.

## Discussion

Dendritic computation occurs as a result of multiple non-linear interactions taking place at dendrites [13, 68]. To provide insights into this phenomenon, we proposed a modelling approach that considers real neurons under naturalistic conditions, receiving independent synaptic-like input at thousands of compartments. These detailed neurons are spatially-extended excitable trees from NeuroMorpho, a database containing over 100,000 digitally reconstructed neurons [2–4, 69–73]. Here we focus our results on a set 26 neurons. Nonetheless, using the code provided, it is straightforward to extend our approach to any neuron of the database.

We demonstrated the presence of substantial spatial dependence in neuronal dynamics that can be attributed to morphological features. Specifically, we mapped the excitability (firing rate) and dynamic range of dendritic branches and the soma. We identified bifurcations as a major structural source that can be very effective in raising the dynamic range. Furthermore, we showed how the number of branches connected to the soma and its centrality influence the energy consumption of neurons, and can be explored to classify neuron types. Hence, we classified neurons into three different families based on centrality and number of branches connecting the soma, and a family that is within a transition zone. We found that a soma with only one branch is special, and a general behavior is expected when the soma has many branches. It is also possible to observe a transition that happens when the soma has two or three branches.

### Neuronal diversity

Diversity is a hallmark of neurons, and this is clearly demonstrated by the large variety of digitally reconstructed neurons found in the NeuroMorpho database, which currently has >100,000 neurons, from >640 cell types, and 60 species. Each of these neurons is unique. They have a tree topology, and their morphological features are crucial for classification. However, additional attempts have also been made to classify neurons based not only on their morphology but taking into account features of their electrophysiology, and their dynamics [1, 9, 74, 75]. These proposals attempt to improve neuronal classification with information about dynamics and function of neurons. Along this line, here we propose to incorporate a few key structural features that inform about neuronal dynamics and function. Utilizing a minimal dynamic model, we were able to simulate the dynamics of many neurons with thousands of compartments. This simple method is suitable to identify and highlight the most important structural features of neurons. Here we focused on a variety of neurons, representing several neuron types, from different species, and acquired at multiple laboratories. Given this diversity, we did not focus on harmonizing the length of compartments. However, NeuroMorpho is a very rich dataset that allows a parcellation that forces compartments to have the same length in order to improve comparisons among neurons.

Action potentials and dendritic spikes consume energy because of the active flux of ions that is required to charge the membrane capacitance. To allow active signaling, these ions have to be pumped and this process uses energy provided by ATP. The energy cost of action potentials can vary considerably across neurons [76]. Here we propose a different approach that focusses not on the cost of an action potential, but on the cost of a neuronal spike relative to the cost of the routing of electric activity through dendrites. We found that our estimation of relative energy consumption of neurons appears in stereotypical forms that can be linked to specific topological features. Hence, we propose that these features are relevant to characterize neuronal dynamics. According to our approach, it is clear that specific morphological fingerprints such as bitufted cells can be classified as belonging to a specific category (T). Furthermore, neurons with multiple branches are more effective at generating somatic spikes. These branches increase the convergence of input to the soma and reduce the overall density of dendritic spikes typically required to trigger a somatic spike. In contrast, neurons with a non-central soma connected to a single branch show the largest relative energy consumption, and they require more than one dendritic spike per compartment for each somatic spike. These distinct behaviors suggest specific computational function for neurons belonging to different families. Moreover, this simple approach reveals a clear role of dendritic topology, which might not be so evident in more complex neuron models that require a large number of parameters.

### Neuronal topology and dynamics

Other aspects of neuronal dynamics affected by dendritic topology include response function and dynamic range. It has been previously shown that the size of dendritic trees play a crucial role in determining the maximum dynamic range an active dendritic tree can attain [38]. However, this proposal was based only on a single topology (a regular and binary Cayley tree) with variable sizes (number of layers). This regular and artificial topology is relevant but the approach does not distinguish clearly the number of compartments from the number of bifurcations. Here, by utilizing real neurons from NeuroMorpho, we can assess the role of these two factors. We found that both the number of bifurcations (Pearson correlation, *r* = 0.72, *P* < 10^−4^) as well as the number of compartments (Pearson correlation, *r* = 0.65, *P* < 10^−4^) can be good predictors for the maximum dynamic range of neurons. However, it must be taken into account that the number of compartments is also correlated with the number of bifurcations. In this dataset, it is important to consider that the number of compartments is mostly determined by the resolution of the digital reconstruction. In contrast, the number of bifurcations reflects more fundamental properties of dendritic topology.

Typically, the larger is the number of bifurcations, the larger is the dynamic range. This trend is also consistent with our minimal modelling approach containing a single bifurcation. We found that a single bifurcation tends to increase the dynamic range by a few decibels. A real dendritic topology can have many bifurcations that contribute to increase the dynamic range of the neuron. Although their contribution is not additive, it is reasonable to find that the number of bifurcations is perhaps the most important element to increase the dynamic range. As large dendrites have many bifurcations, the resolution used to digitalize such a complex structure needs to be fine. As a result, the number of compartments often correlates with the number of bifurcations. Together, our results suggest that the number of bifurcations is likely among the most essential features of dendrites to shape the dynamic range.

Centrality is also fundamental for the dynamics because central branches exhibit larger firing rates. This feature tends to increase the dynamic range as it is very effective at amplifying weak inputs. However, this amplification can also be beneficial to the dynamic range when *P* approaches 1. In this case, when *P* is nearly deterministic, we found that the dynamic range of central compartments is usually higher than at non-central ones (see Fig. S8).

### Neuronal models

Our model is suitable to explore the topological effects of tens of thousands of digitally reconstructed neurons with thousands of compartments in neuronal dynamics under complex input conditions. In order to focus on these crucial spatial aspects of neurons, here we considered explicitly simplified neuronal dynamics. We disregarded heterogeneity of ion channels along neurons and assumed homogeneity for simplicity. This allows us to highlight topological features of neurons and ascribe the heterogeneous types of dynamics exclusively to dendritic structure. We argue that our simple modelling approach retains the essential features to simulate the dynamics of excitable systems without the burden of an excessive number of details and parameters. However, this is clearly a simplification. Future work should address the role of more sophisticated biophysical models with additional free parameters that describe the membrane potential of dendritic branches as continuous variables (differential equations, instead of a map with discrete states [77]). These more detailed models might also incorporate heterogeneity of dynamics, taking into account impedance gradients, the dendritic diameter, type of dendrite, distance from soma, and so on [78]. Moreover, our simplifying assumptions of homogeneous and constant input can also be extended in future works. As a first step, a more detailed description of the dynamics of the soma can be considered in which the model has two types of dynamics, one for the dendrites and one for the soma, and the role of two types of integration can be independently assessed. This distinction might be relevant because it is known that signal integration at the soma can be crucial for coincidence detection and the resulting network response[79].

Changes in neuronal structure are reported in many neuropsychiatric disorders [80–84]. The minimal modelling approach proposed here can be used to characterize the changes in neuronal dynamics caused by these structural alterations [85]. Our simplified approach might also be relevant for simulating motifs and circuits of neurons with detailed dendritic structure under complex and realistic input conditions. Future work can focus on more complex and specific extensions of the model. For example, spatial- and time-dependent input may reveal other main features of neuronal dynamics with dendritic computation taking place in parallel at functional subunits in cortical circuits.

### Conclusions

Our model provides insights into the role the dendritic structure plays in the behavior of a neuron, both local and on the larger scale. We used digital reconstructions of real neurons to address the effects of intricate and nonhomogeneous spatial features of neurons on their dynamics. Our results indicate that two main morphological features – the centrality of the soma, and the number of branches connected to the soma – can determine the type of behavior a neuron exhibits. Neurons whose soma lies on the border and in a non-central location are intrinsically energy inefficient, whereas neurons with many branches connected to the soma are intrinsically energy efficient. Furthermore, we have shown that bifurcations in the dendritic tree can enhance the dynamic range, and that the maximum dynamic range of neurons increase with the number of bifurcations and compartments. Our approach can be extended to more than 100,000 neurons available at the NeuroMorpho database.

## Code availability

MATLAB code to simulate neuronal activity is available at http://www.sng.org.au/Downloads.

## Acknowledgements

This work was supported by the Australian Research Council and the Australian National Health and Medical Research Council (APP1110975). The authors are grateful for additional financial support from the Dentons Australia Honours Scholarship.

## Supporting information captions

**Video S1. Demonstration of the neurodynamics**. Here we use the input rate *h* = 0.1 Hz and transmission probability *P* = 0.96, and neuron reconstruction A (see Methods).

**Figure S1. Firing rate heat maps of all neurons varied over P**. The activation rate is held constant, *h* = 0.1 Hz, and the firing rates were averaged over five trials. The soma is marked by the arrow (left panel). Note the different color bars.

**Figure S2. Firing rate heat maps of all neurons varied over h**. The transmission probability is held constant, *P* = 0.9, and the firing rates were averaged over five trials. The soma is marked by the arrow. Note the different color bars.

**Figure S3. Dynamic range heat maps of all neurons varied over P**. Values were averaged over five trials. The soma is marked by the arrow. Note the different color bars.

**Figure S4. Compartmental dynamic range as a function of distance from soma for all neurons**. See methods for description of the neurons.

**Figure S5. Additional dynamics of benchmark neurites**. The main branch consists of *N* = 1000 compartments. The value of *P* is constant for each collection of sub-plots, and plots are arranged according to the position (*Q*) and length (*L*) of the secondary branch. In each small sub-plot, the compartmental dynamic range (y axis, in dB) is plotted against the distance from the soma (x axis), with shadings indicating ± 1 standard deviation from the mean over 10 trials. Each trial was run for 105 time steps (ms). As *P* increases, the amplitude and extent of the spike in dynamic range at the bifurcation point increases. However, for large *P*, short secondary branches do not have a significant effect on the dynamic range.

**Figure S6. Energy consumption plots of all neurons**. The energy consumption is a measure of how often each dendritic spike fires per somatic spike (see Methods). There are several distinct behaviors, which we attribute to the number of branches connected to the soma, and the relative centrality of the soma. Note the different color bars.

**Figure S7. Relative centrality heat maps for all neurons**. Red corresponds to the most central compartments (*C* = 1), while blue corresponds to the least central ones (*C* = 0). See Methods for details. The location of the soma is indicated by the arrow.

**Figure S8. Correlation between centrality and dynamic range**. Scatter plot of relative compartmental centrality (see Methods for calculation) vs relative compartmental dynamic range for all compartments in all neurons A-Z (total number of compartments is *N* = 47411). The relative dynamic range is Δ_rel_ = 1 if a compartment has the largest dynamic range in the neuron, and Δ_rel_ = 0 if it has the lowest. The soma of each neuron is marked in red. For high values of *P*, the relative centrality and relative dynamic range correlate.

## References

1. Masland, R.H., Neuronal cell types. Current Biology, 2004. 14(13): p. R497–R500.

2. Ascoli, G.A., Sharing neuron data: carrots, sticks, and digital records. PLoS biology, 2015. 13(10): p. e1002275.

3. Ascoli, G.A., D.E. Donohue, and M. Halavi, NeuroMorpho. Org: a central resource for neuronal morphologies. Journal of Neuroscience, 2007. 27(35): p. 9247–9251.

4. Halavi, M., et al., Digital reconstructions of neuronal morphology: three decades of research trends. Frontiers in neuroscience, 2012. 6: p. 49.

5. Hines, M.L., T.M. Morse, and N.T. Carnevale, Model structure analysis in NEURON: toward interoperability among neural simulators, in Neuroinformatics. 2007, Humana Press. p. 91–102.

6. Van Ooyen, A., Using theoretical models to analyse neural development. Nature Reviews Neuroscience, 2011. 12(6): p. 311.

7. van Pelt, J., A. van Ooyen, and H.B. Uylings, The need for integrating neuronal morphology databases and computational environments in exploring neuronal structure and function. Anatomy and embryology, 2001. 204(4): p. 255–265.

8. Wearne, S., et al., New techniques for imaging, digitization and analysis of three-dimensional neural morphology on multiple scales. Neuroscience, 2005. 136(3): p. 661–680.

9. Sharpee, T.O., Toward functional classification of neuronal types. Neuron, 2014. 83(6): p. 1329–1334.

10. Kanari, L., et al., Objective Morphological Classification of Neocortical Pyramidal Cells. Cerebral Cortex, 2019. 29(4): p. 1719–1735.

11. Wen, Q. and D.B. Chklovskii, A cost–benefit analysis of neuronal morphology. Journal of neurophysiology, 2008. 99(5): p. 2320–2328.

12. Brunel, N. and M.C. Van Rossum, Lapicque’s 1907 paper: from frogs to integrate-and-fire. Biological cybernetics, 2007. 97(5-6): p. 337–339.

13. Häusser, M., N. Spruston, and G.J. Stuart, Diversity and dynamics of dendritic signaling. Science, 2000. 290(5492): p. 739–744.

14. Baer, S. and J. Rinzel, Propagation of dendritic spikes mediated by excitable spines: a continuum theory. Journal of neurophysiology, 1991. 65(4): p. 874–890.

15. Häusser, M. and B. Mel, Dendrites: bug or feature? Current opinion in neurobiology, 2003. 13(3): p. 372–383.

16. Gollo, L.L., O. Kinouchi, and M. Copelli, Active dendrites enhance neuronal dynamic range. PLoS computational biology, 2009. 5(6): p. e1000402.

17. Schmidt-Hieber, C., P. Jonas, and J. Bischofberger, Subthreshold dendritic signal processing and coincidence detection in dentate gyrus granule cells. Journal of Neuroscience, 2007. 27(31): p. 8430–8441.

18. De Sousa, G., et al., Dendritic morphology predicts pattern recognition performance in multi-compartmental model neurons with and without active conductances. Journal of computational neuroscience, 2015. 38(2): p. 221–234.

19. Koch, C. and I. Segev, The role of single neurons in information processing. Nature neuroscience, 2000. 3(11s): p. 1171.

20. Sardi, S., et al., New types of experiments reveal that a neuron functions as multiple independent threshold units. Scientific reports, 2017. 7(1): p. 18036.

21. Zang, Y., S. Dieudonné, and E. De Schutter, Voltage-and Branch-Specific Climbing Fiber Responses in Purkinje Cells. Cell reports, 2018. 24(6): p. 1536–1549.

22. Naud, R., A. Payeur, and A. Longtin, Noise gated by dendrosomatic interactions increases information transmission. Physical Review X, 2017. 7(3): p. 031045.

23. Gollo, L.L., O. Kinouchi, and M. Copelli, Statistical physics approach to dendritic computation: The excitable-wave mean-field approximation. Physical Review E, 2012. 85(1): p. 011911.

24. Avena-Koenigsberger, A., B. Misic, and O. Sporns, Communication dynamics in complex brain networks. Nature Reviews Neuroscience, 2018. 19(1): p. 17.

25. Segev, I. and M. London, Untangling dendrites with quantitative models. Science, 2000. 290(5492): p. 744–750.

26. Van Ooyen, A., et al., The effect of dendritic topology on firing patterns in model neurons. Network: Computation in neural systems, 2002. 13(3): p. 311–325.

27. Remme, M.W., J. Rinzel, and S. Schreiber, Function and energy consumption constrain neuronal biophysics in a canonical computation: Coincidence detection. PLoS computational biology, 2018. 14(12): p. e1006612.

28. Shepherd, G., et al., Signal enhancement in distal cortical dendrites by means of interactions between active dendritic spines. Proceedings of the National Academy of Sciences, 1985. 82(7): p. 2192–2195.

29. Keren, N., N. Peled, and A. Korngreen, Constraining compartmental models using multiple voltage-recordings and genetic algorithms. Journal of neurophysiology, 2005.

30. Mainen, Z.F. and T.J. Sejnowski, Influence of dendritic structure on firing pattern in model neocortical neurons. Nature, 1996. 382(6589): p. 363.

31. Eyal, G., et al., Human cortical pyramidal neurons: From spines to spikes via models. bioRxiv, 2018: p. 267898.

32. Zandt, B.-J., M.L. Veruki, and E. Hartveit, Electrotonic signal processing in AII amacrine cells: compartmental models and passive membrane properties for a gap junction-coupled retinal neuron. Brain Structure and Function, 2018. 223(7): p. 3383–3410.

33. Poirazi, P., T. Brannon, and B.W. Mel, Pyramidal neuron as two-layer neural network. Neuron, 2003. 37(6): p. 989–999.

34. Hay, E., et al., Models of neocortical layer 5b pyramidal cells capturing a wide range of dendritic and perisomatic active properties. PLoS computational biology, 2011. 7(7): p. e1002107.

35. Kinouchi, O. and M. Copelli, Optimal dynamical range of excitable networks at criticality. Nature physics, 2006. 2(5): p. 348.

36. Carnevale, N.T., et al., Comparative electrotonic analysis of three classes of rat hippocampal neurons. Journal of Neurophysiology, 1997. 78(2): p. 703–720.

37. Butler, A.B. and W. Hodos, Comparative vertebrate neuroanatomy: evolution and adaptation. 2005: John Wiley & Sons.

38. Gollo, L.L., O. Kinouchi, and M. Copelli, Single-neuron criticality optimizes analog dendritic computation. Scientific reports, 2013. 3: p. 3222.

39. D’Souza, R.D., et al., Recruitment of inhibition and excitation across mouse visual cortex depends on the hierarchy of interconnecting areas. Elife, 2016. 5: p. e19332.

40. Jacobs, B., et al., Comparative morphology of gigantopyramidal neurons in primary motor cortex across mammals. Journal of Comparative Neurology, 2018. 526(3): p. 496–536.

41. Watson, K.K., T.K. Jones, and J.M. Allman, Dendritic architecture of the von Economo neurons. Neuroscience, 2006. 141(3): p. 1107–1112.

42. Eyal, G., et al., Unique membrane properties and enhanced signal processing in human neocortical neurons. Elife, 2016. 5: p. e16553.

43. Mazzoni, F., E. Novelli, and E. Strettoi, Retinal ganglion cells survive and maintain normal dendritic morphology in a mouse model of inherited photoreceptor degeneration. Journal of Neuroscience, 2008. 28(52): p. 14282–14292.

44. Jacobs, B., et al., Regional dendritic and spine variation in human cerebral cortex: a quantitative golgi study. Cerebral cortex, 2001. 11(6): p. 558–571.

45. Coombs, J., et al., Morphological properties of mouse retinal ganglion cells. Neuroscience, 2006. 140(1): p. 123–136.

46. Seco, C.Z., et al., A homozygous FITM2 mutation causes a deafness-dystonia syndrome with motor regression and signs of ichthyosis and sensory neuropathy. Disease models & mechanisms, 2017. 10(2): p. 105–118.

47. Bastian, T.W., et al., Eltrombopag, a thrombopoietin mimetic, crosses the blood–brain barrier and impairs iron-dependent hippocampal neuron dendrite development. Journal of Thrombosis and Haemostasis, 2017. 15(3): p. 565–574.

48. Kougias, D.G., et al., Beta-hydroxy-beta-methylbutyrate ameliorates aging effects in the dendritic tree of pyramidal neurons in the medial prefrontal cortex of both male and female rats. Neurobiology of aging, 2016. 40: p. 78–85.

49. Routh, B.N., et al., Anatomical and electrophysiological comparison of CA1 pyramidal neurons of the rat and mouse. Journal of neurophysiology, 2009. 102(4): p. 2288–2302.

50. Boillot, M., et al., LGI1 acts presynaptically to regulate excitatory synaptic transmission during early postnatal development. Scientific reports, 2016. 6: p. 21769.

51. Briggs, F., et al., Morphological substrates for parallel streams of corticogeniculate feedback originating in both V1 and V2 of the macaque monkey. Neuron, 2016. 90(2): p. 388–399.

52. Rihn, L.L. and B.J. Claiborne, Dendritic growth and regression in rat dentate granule cells during late postnatal development. Developmental Brain Research, 1990. 54(1): p. 115–124.

53. Martone, M.E., et al., The cell-centered database. Neuroinformatics, 2003. 1(4): p. 379–395.

54. Chapleau, C.A., et al., Dendritic spine pathologies in hippocampal pyramidal neurons from Rett syndrome brain and after expression of Rett-associated MECP2 mutations. Neurobiology of disease, 2009. 35(2): p. 219–233.

55. Kuddannaya, S., et al., Geometrically Mediated Topographic Steering of Neurite Behaviors and Network Formation. Advanced Materials Interfaces, 2018. 5(7): p. 1700819.

56. Bu, Q., et al., CREB signaling is involved in Rett syndrome pathogenesis. Journal of Neuroscience, 2017. 37(13): p. 3671–3685.

57. Jayabal, S., L. Ljungberg, and A.J. Watt, Transient cerebellar alterations during development prior to obvious motor phenotype in a mouse model of spinocerebellar ataxia type 6. The Journal of physiology, 2017. 595(3): p. 949–966.

58. Cuntz, H., et al., Preserving neural function under extreme scaling. PloS one, 2013. 8(8): p. e71540.

59. Radley, J.J., et al., Chronic stress-induced alterations of dendritic spine subtypes predict functional decrements in an hypothalamo–pituitary–adrenal-inhibitory prefrontal circuit. Journal of Neuroscience, 2013. 33(36): p. 14379–14391.

60. Fukumitsu, K., et al., Mitochondrial fission protein Drp1 regulates mitochondrial transport and dendritic arborization in cerebellar Purkinje cells. Molecular and Cellular Neuroscience, 2016. 71: p. 56–65.

61. Nedelescu, H., M. Abdelhack, and A.T. Pritchard, Regional differences in Purkinje cell morphology in the cerebellar vermis of male mice. Journal of neuroscience research, 2018. 96(9): p. 1476–1489.

62. Royer, A.S. and R.F. Miller, Dendritic impulse collisions and shifting sites of action potential initiation contract and extend the receptive field of an amacrine cell. Visual neuroscience, 2007. 24(4): p. 619–634.

63. Waters, J., A. Schaefer, and B. Sakmann, Backpropagating action potentials in neurones: measurement, mechanisms and potential functions. Progress in biophysics and molecular biology, 2005. 87(1): p. 145–170.

64. Gollo, L.L., Coexistence of critical sensitivity and subcritical specificity can yield optimal population coding. Journal of The Royal Society Interface, 2017. 14(134): p. 20170207.

65. Gollo, L.L., M. Copelli, and J.A. Roberts, Diversity improves performance in excitable networks. PeerJ, 2016. 4: p. e1912.

66. Hasenstaub, A., et al., Metabolic cost as a unifying principle governing neuronal biophysics. Proceedings of the National Academy of Sciences, 2010. 107(27): p. 12329–12334.

67. Publio, R., C.C. Ceballos, and A.C. Roque, Dynamic range of vertebrate retina ganglion cells: Importance of active dendrites and coupling by electrical synapses. PloS one, 2012. 7(10): p. e48517.

68. London, M. and M. Häusser, Dendritic computation. Annu. Rev. Neurosci., 2005. 28: p. 503–532.

69. Halavi, M., et al., NeuroMorpho. Org implementation of digital neuroscience: dense coverage and integration with the NIF. Neuroinformatics, 2008. 6(3): p. 241.

70. Donohue, D.E. and G.A. Ascoli, Automated reconstruction of neuronal morphology: an overview. Brain research reviews, 2011. 67(1-2): p. 94–102.

71. Smith, K., L. Seligman, and V. Swarup, Everybody share: The challenge of data-sharing systems. Computer, 2008. 41(9): p. 54–61.

72. Parekh, R. and G.A. Ascoli, Neuronal morphology goes digital: a research hub for cellular and system neuroscience. Neuron, 2013. 77(6): p. 1017–1038.

73. Ascoli, G.A., et al., Win–win data sharing in neuroscience. Nature methods, 2017. 14(2): p. 112.

74. Markram, H., et al., Interneurons of the neocortical inhibitory system. Nature reviews neuroscience, 2004. 5(10): p. 793.

75. Mott, D.D. and R. Dingledine, Interneuron Diversity series: Interneuron research–challenges and strategies. Trends in neurosciences, 2003. 26(9): p. 484–488.

76. Sengupta, B., et al., Action potential energy efficiency varies among neuron types in vertebrates and invertebrates. PLoS computational biology, 2010. 6(7).

77. Girardi-Schappo, M., M. Tragtenberg, and O. Kinouchi, A brief history of excitable map-based neurons and neural networks. Journal of neuroscience methods, 2013. 220(2): p. 116–130.

78. Donohue, D.E. and G.A. Ascoli, A comparative computer simulation of dendritic morphology. PLoS computational biology, 2008. 4(6): p. e1000089.

79. Gollo, L.L., C. Mirasso, and V.M. Eguíluz, Signal integration enhances the dynamic range in neuronal systems. Physical Review E, 2012. 85(4): p. 040902.

80. Uylings, H.B. and J. De Brabander, Neuronal changes in normal human aging and Alzheimer’s disease. Brain and cognition, 2002. 49(3): p. 268–276.

81. Coleman, P.D. and D.G. Flood, Neuron numbers and dendritic extent in normal aging and Alzheimer’s disease. Neurobiology of aging, 1987. 8(6): p. 521–545.

82. Kulkarni, V.A. and B.L. Firestein, The dendritic tree and brain disorders. Molecular and Cellular Neuroscience, 2012. 50(1): p. 10–20.

83. Raymond, G.V., M.L. Bauman, and T.L. Kemper, Hippocampus in autism: a Golgi analysis. Acta neuropathologica, 1995. 91(1): p. 117–119.

84. Forrest, M.P., E. Parnell, and P. Penzes, Dendritic structural plasticity and neuropsychiatric disease. Nature reviews Neuroscience, 2018. 19(4): p. 215.

85. Kirch, C. and L.L. Gollo, Dynamical effects of dendritic pruning implicated in aging and neurodegeneration: Towards a measure of neuronal reserve. bioRxiv, 2020: p. 2020.04.09.035048.

