## SupplementaryFigures for "Spatially resolved dendritic integration: Towards a functional classification of neurons"

### Figure S1. Firing rate heat maps of all neurons varied over P.

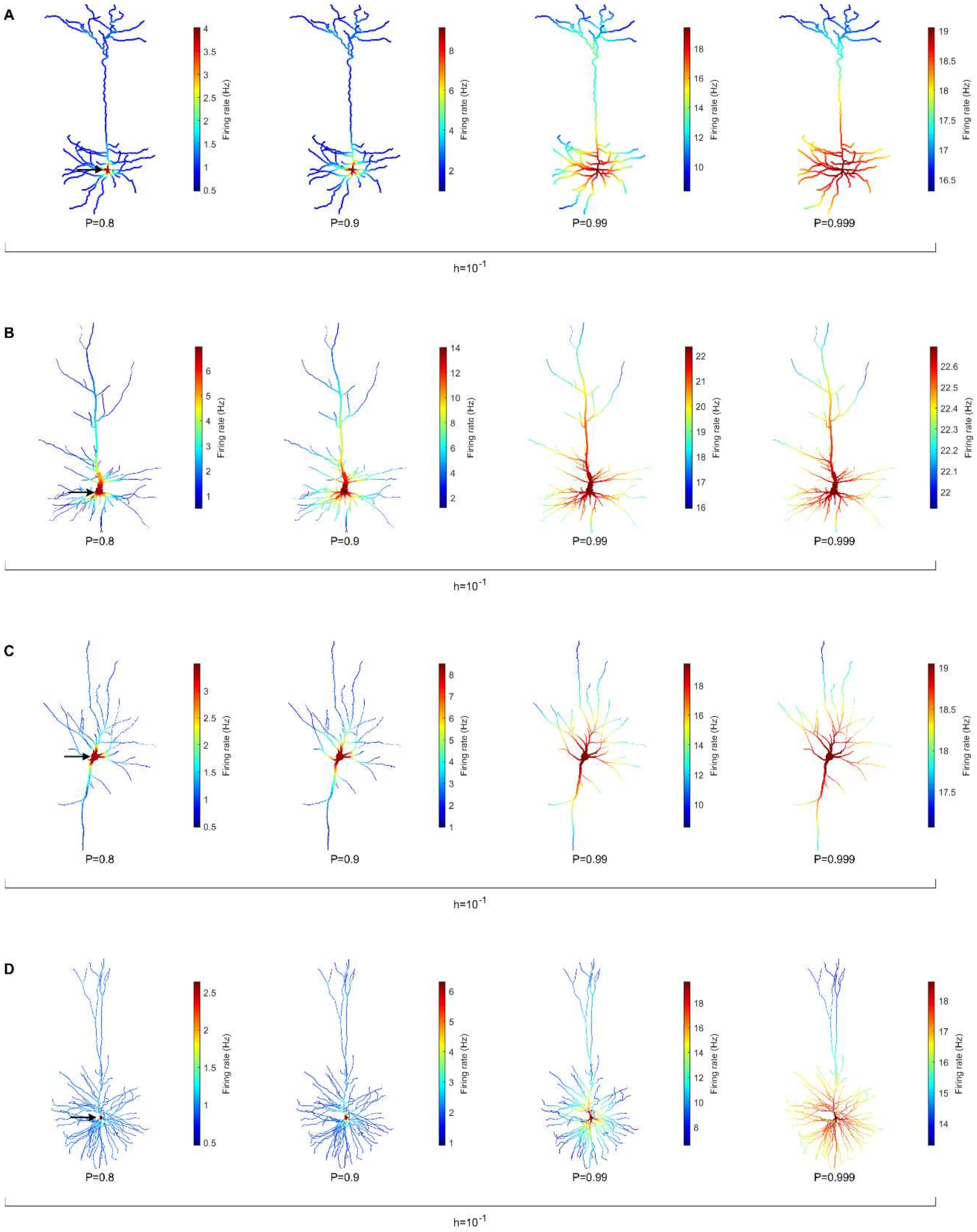

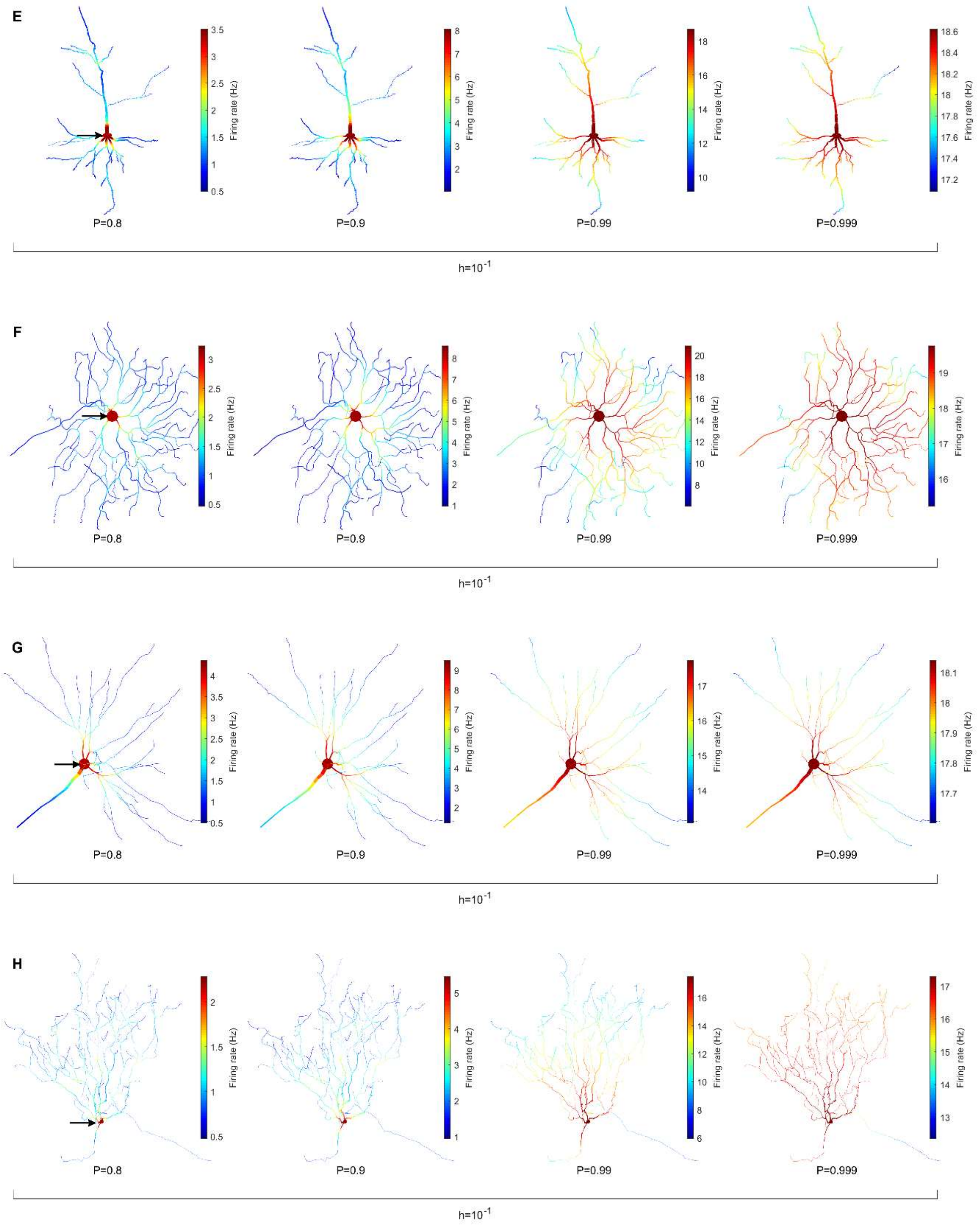
E
Firing rate (Hz)
P=0.8
P=0.9
P=0.99
P=0.999
h=10^-1
F
Firing rate (Hz)
P=0.8
P=0.9
P=0.99
P=0.999
h=10^-1
G
Firing rate (Hz)
P=0.8
P=0.9
P=0.99
P=0.999
h=10^-1
H
Firing rate (Hz)
P=0.8
P=0.9
P=0.99
P=0.999
h=10^-1

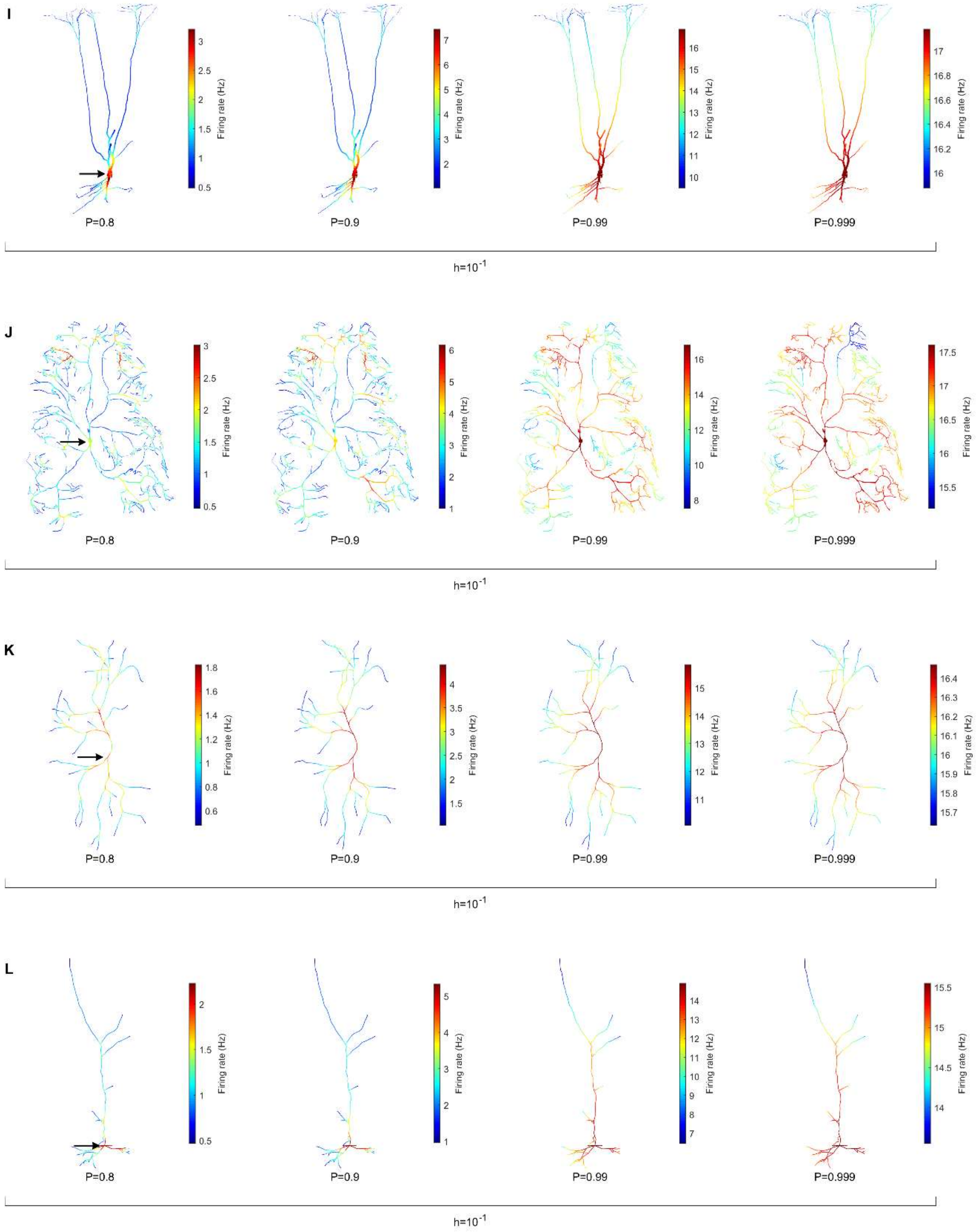
I
Firing rate (Hz)
P=0.8
P=0.9
P=0.99
P=0.999
h=$10^{-1}$
J
Firing rate (Hz)
P=0.8
P=0.9
P=0.99
P=0.999
h=$10^{-1}$
K
Firing rate (Hz)
P=0.8
P=0.9
P=0.99
P=0.999
h=$10^{-1}$
L
Firing rate (Hz)
P=0.8
P=0.9
P=0.99
P=0.999
h=$10^{-1}$

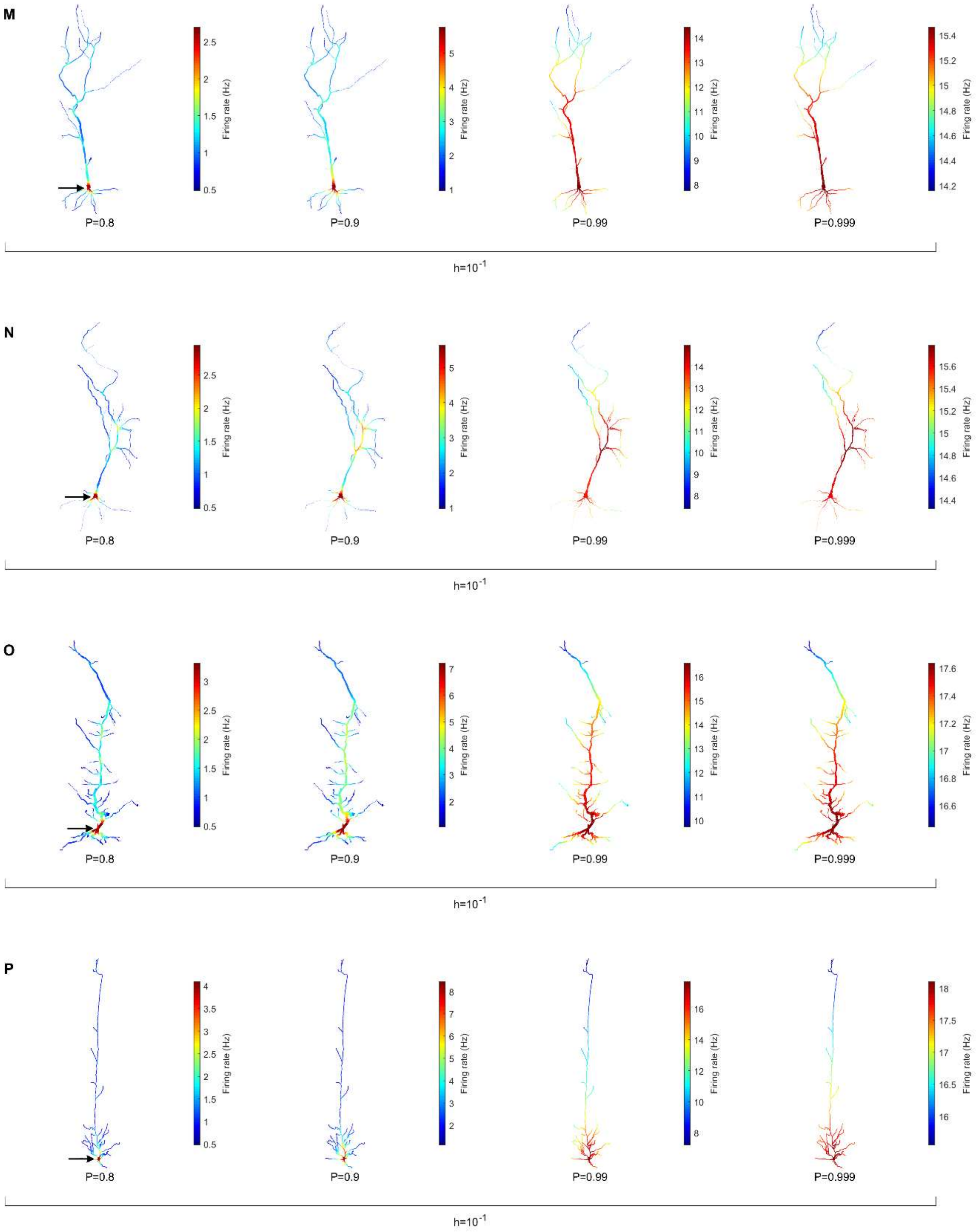

M
2.5
2
1.5
1
0.5
Firing rate (Hz)
5
4
3
2
1
Firing rate (Hz)
14
13
12
11
10
9
8
Firing rate (Hz)
15.4
15.2
15
14.8
14.6
14.4
14.2
Firing rate (Hz)
P=0.8
P=0.9
P=0.99
P=0.999
h=10$^{-1}$
N
2.5
2
1.5
1
0.5
Firing rate (Hz)
5
4
3
2
1
Firing rate (Hz)
14
13
12
11
10
9
8
Firing rate (Hz)
15.6
15.4
15.2
15
14.8
14.6
14.4
Firing rate (Hz)
P=0.8
P=0.9
P=0.99
P=0.999
h=10$^{-1}$
O
3
2.5
2
1.5
1
0.5
Firing rate (Hz)
7
6
5
4
3
2
Firing rate (Hz)
16
15
14
13
12
11
10
Firing rate (Hz)
17.6
17.4
17.2
17
16.8
16.6
Firing rate (Hz)
P=0.8
P=0.9
P=0.99
P=0.999
h=10$^{-1}$
P
4
3.5
3
2.5
2
1.5
1
0.5
Firing rate (Hz)
8
7
6
5
4
3
2
Firing rate (Hz)
16
14
12
10
8
Firing rate (Hz)
18
17.5
17
16.5
16
Firing rate (Hz)
P=0.8
P=0.9
P=0.99
P=0.999
h=10$^{-1}$

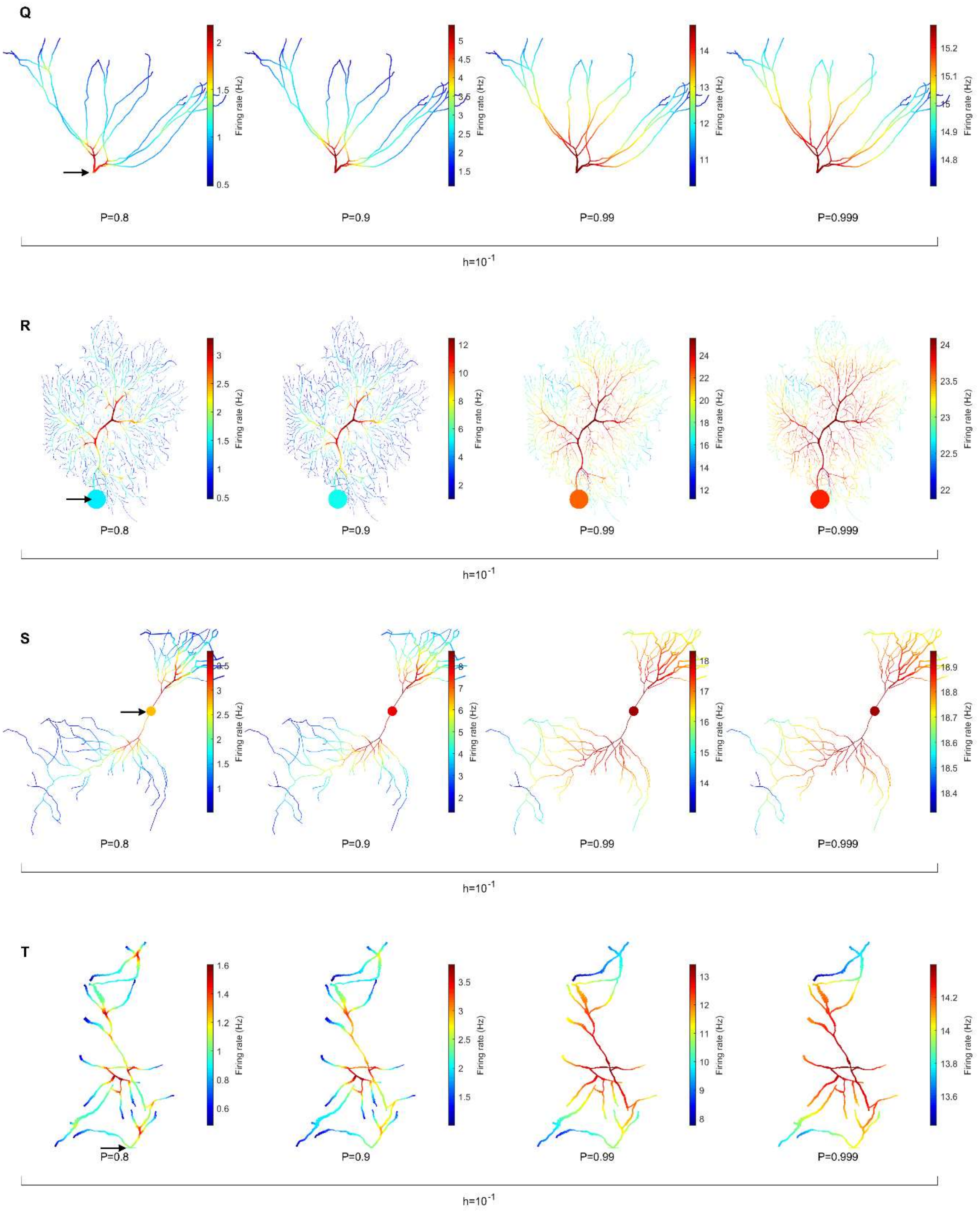
Q
Firing rate (Hz)
P=0.8
P=0.9
P=0.99
P=0.999
h=10^-1
R
Firing rate (Hz)
P=0.8
P=0.9
P=0.99
P=0.999
h=10^-1
S
Firing rate (Hz)
P=0.8
P=0.9
P=0.99
P=0.999
h=10^-1
T
Firing rate (Hz)
P=0.8
P=0.9
P=0.99
P=0.999
h=10^-1

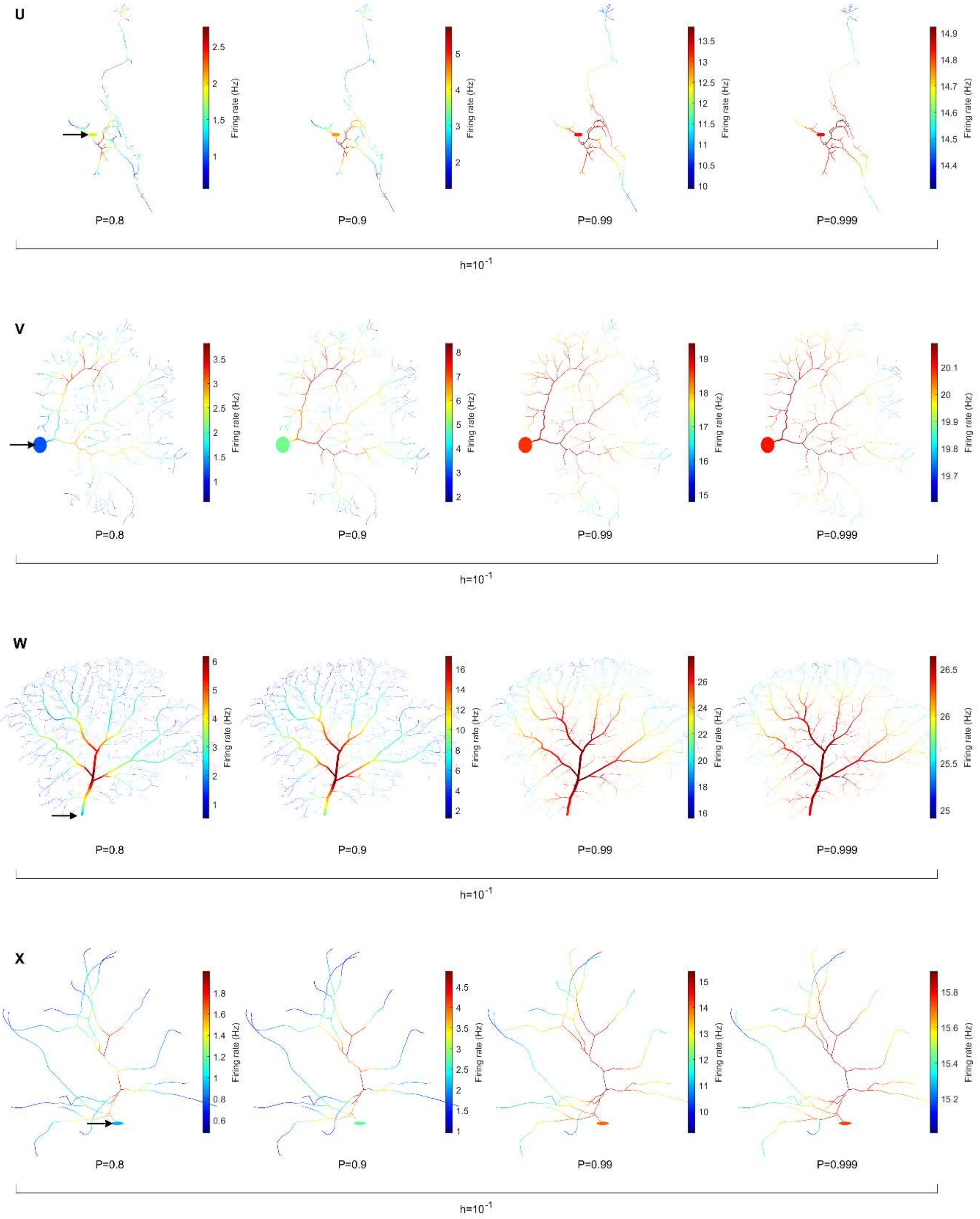
U
Firing rate (Hz)
2.5
2
1.5
1
5
4
3
2
13.5
13
12.5
12
11.5
11
10.5
10
14.9
14.8
14.7
14.6
14.5
14.4
P=0.8
P=0.9
P=0.99
P=0.999
h=10^-1
V
Firing rate (Hz)
3.5
3
2.5
2
1.5
1
8
7
6
5
4
3
2
19
18
17
16
15
20.1
20
19.9
19.8
19.7
P=0.8
P=0.9
P=0.99
P=0.999
h=10^-1
W
Firing rate (Hz)
6
5
4
3
2
1
16
14
12
10
8
6
4
2
26
24
22
20
18
16
26.5
26
25.5
25
P=0.8
P=0.9
P=0.99
P=0.999
h=10^-1
X
Firing rate (Hz)
1.8
1.6
1.4
1.2
1
0.8
0.6
4.5
4
3.5
3
2.5
2
1.5
1
15
14
13
12
11
10
15.8
15.6
15.4
15.2
P=0.8
P=0.9
P=0.99
P=0.999
h=10^-1

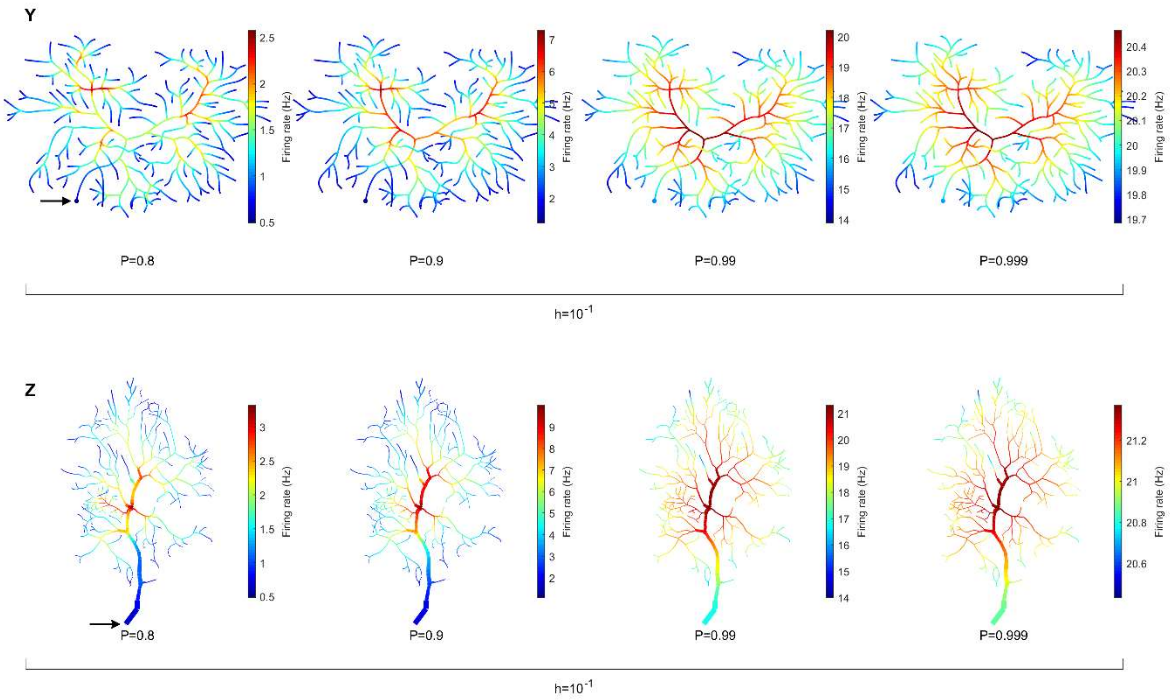


### Figure S2. Firing rate heat maps of all neurons varied over h.

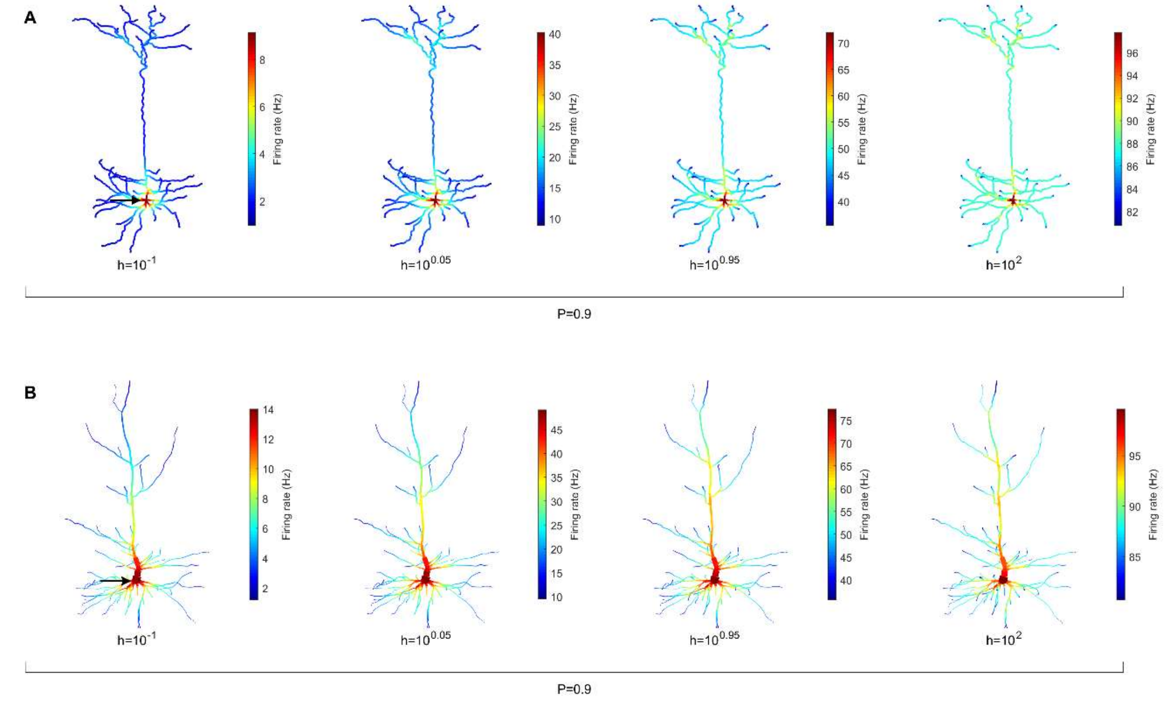

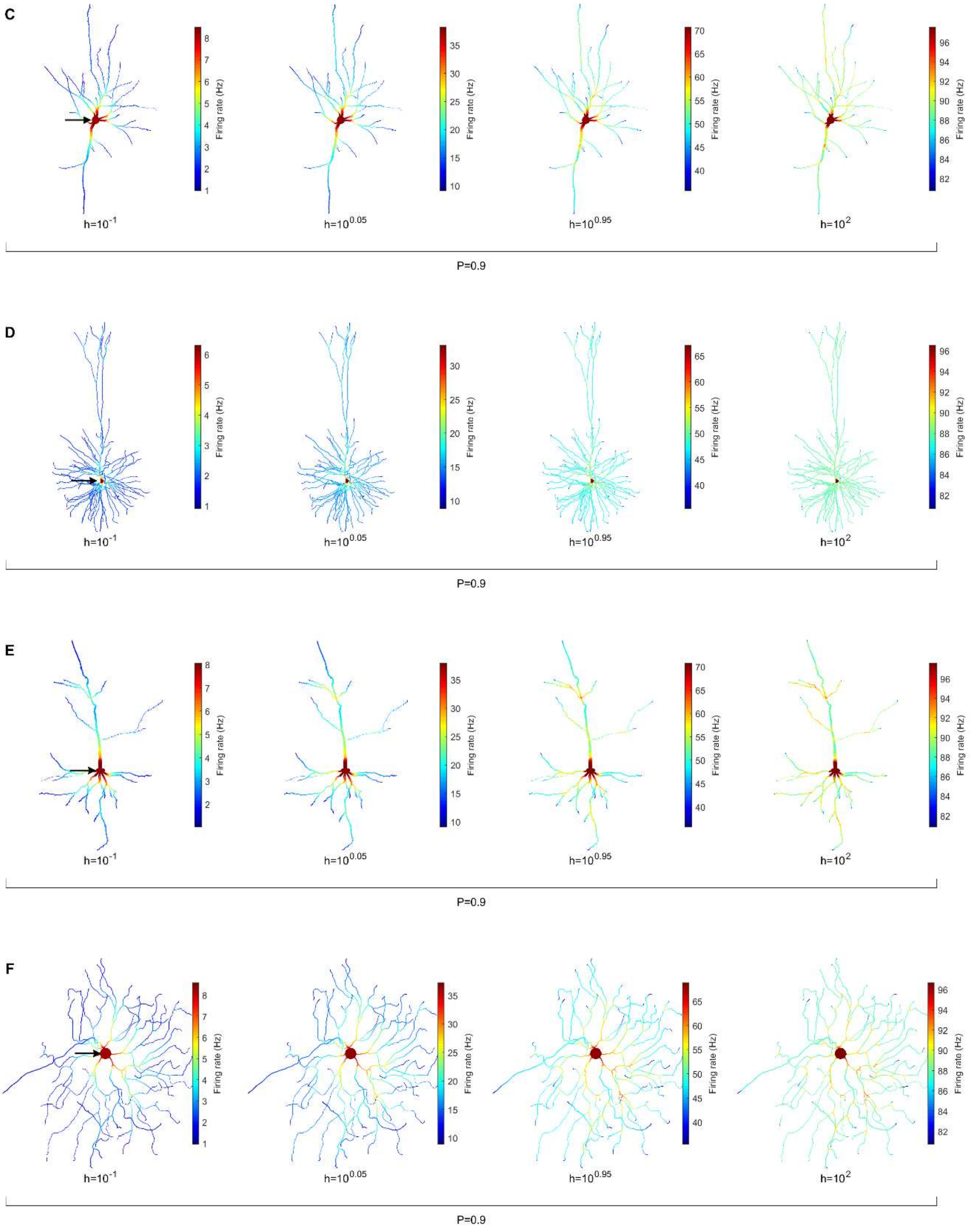

C
Firing rate (Hz)
h=10$^{-1}$
h=10$^{0.05}$
h=10$^{0.95}$
h=10$^{2}$
P=0.9
D
Firing rate (Hz)
h=10$^{-1}$
h=10$^{0.05}$
h=10$^{0.95}$
h=10$^{2}$
P=0.9
E
Firing rate (Hz)
h=10$^{-1}$
h=10$^{0.05}$
h=10$^{0.95}$
h=10$^{2}$
P=0.9
F
Firing rate (Hz)
h=10$^{-1}$
h=10$^{0.05}$
h=10$^{0.95}$
h=10$^{2}$
P=0.9

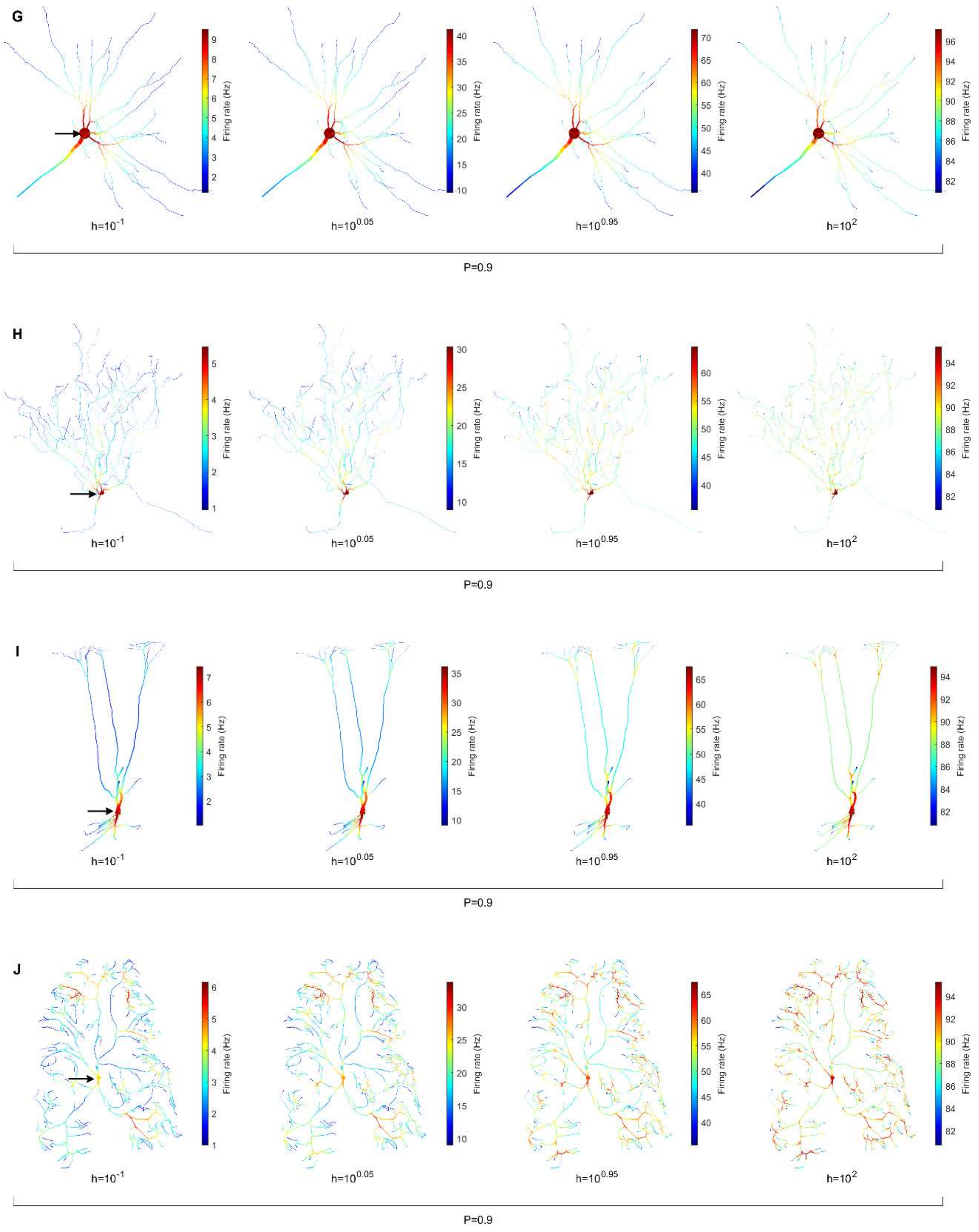
G
Firing rate (Hz)
h=10^-1
h=10^0.05
h=10^0.95
h=10^2
P=0.9
H
Firing rate (Hz)
h=10^-1
h=10^0.05
h=10^0.95
h=10^2
P=0.9
I
Firing rate (Hz)
h=10^-1
h=10^0.05
h=10^0.95
h=10^2
P=0.9
J
Firing rate (Hz)
h=10^-1
h=10^0.05
h=10^0.95
h=10^2
P=0.9

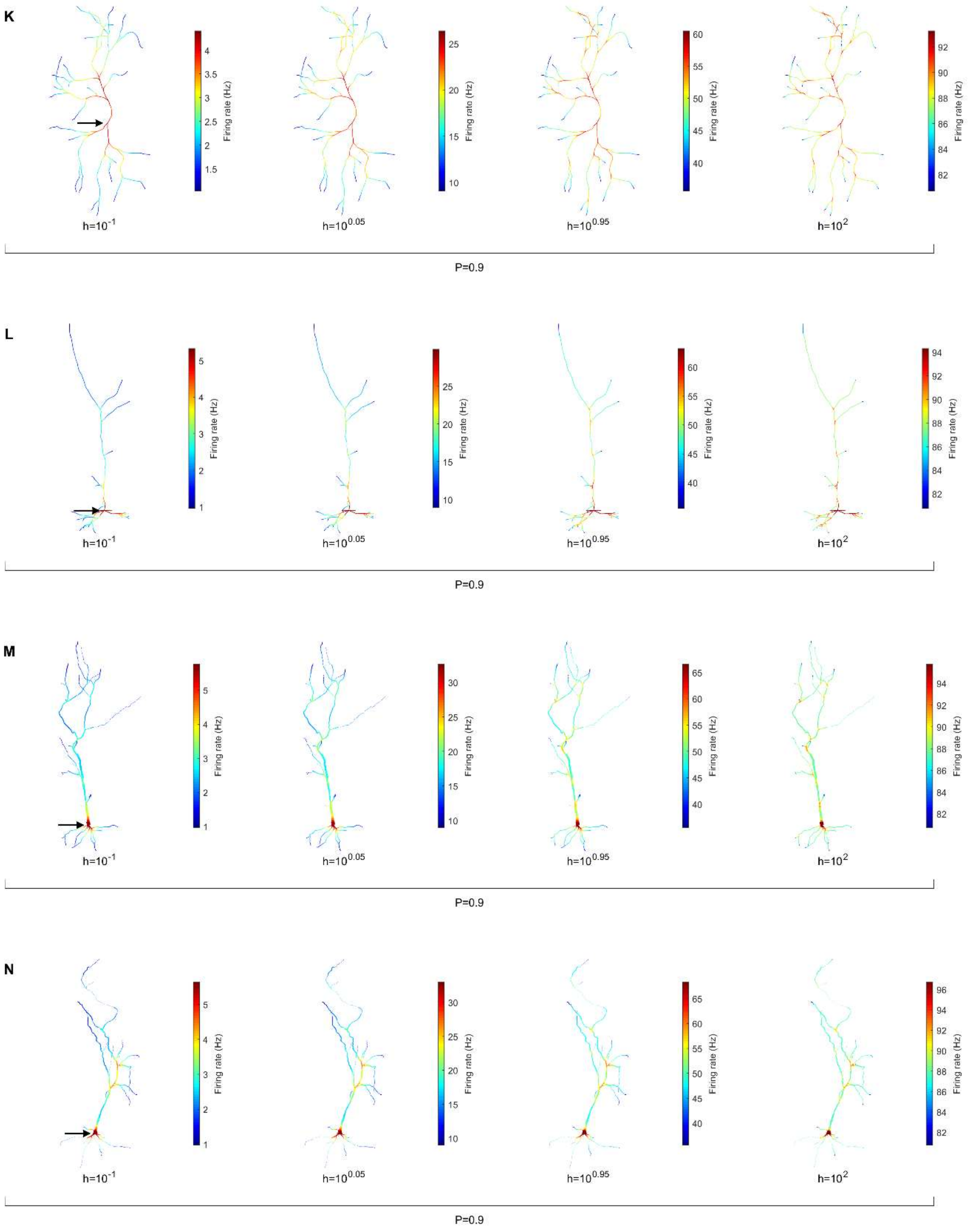
K
Firing rate (Hz)
h=10^-1
h=10^0.05
h=10^0.95
h=10^2
P=0.9
L
Firing rate (Hz)
h=10^-1
h=10^0.05
h=10^0.95
h=10^2
P=0.9
M
Firing rate (Hz)
h=10^-1
h=10^0.05
h=10^0.95
h=10^2
P=0.9
N
Firing rate (Hz)
h=10^-1
h=10^0.05
h=10^0.95
h=10^2
P=0.9

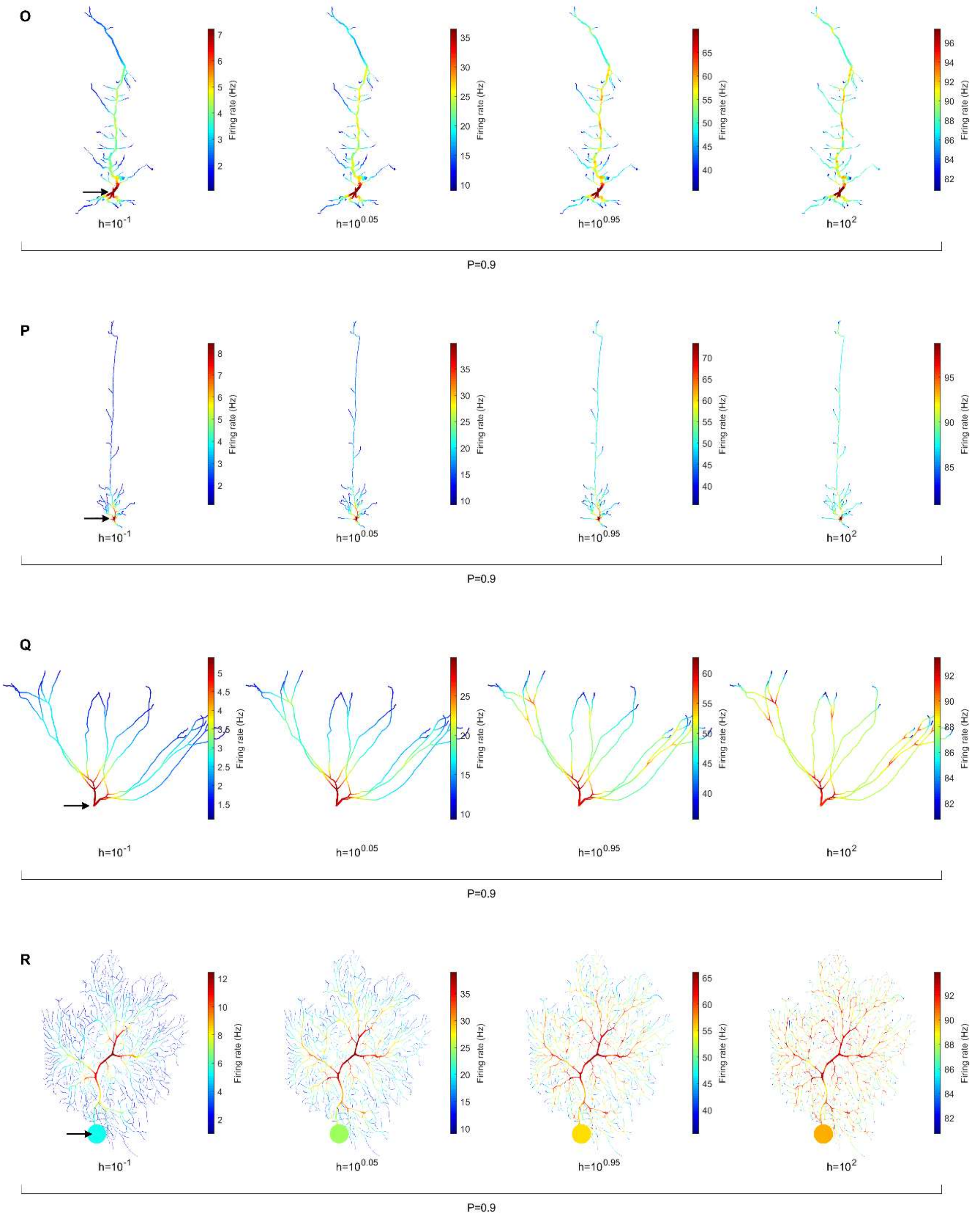
O
Firing rate (Hz)
h=10$^{-1}$
h=10$^{0.05}$
h=10$^{0.95}$
h=10$^{2}$
P=0.9
P
Firing rate (Hz)
h=10$^{-1}$
h=10$^{0.05}$
h=10$^{0.95}$
h=10$^{2}$
P=0.9
Q
Firing rate (Hz)
h=10$^{-1}$
h=10$^{0.05}$
h=10$^{0.95}$
h=10$^{2}$
P=0.9
R
Firing rate (Hz)
h=10$^{-1}$
h=10$^{0.05}$
h=10$^{0.95}$
h=10$^{2}$
P=0.9

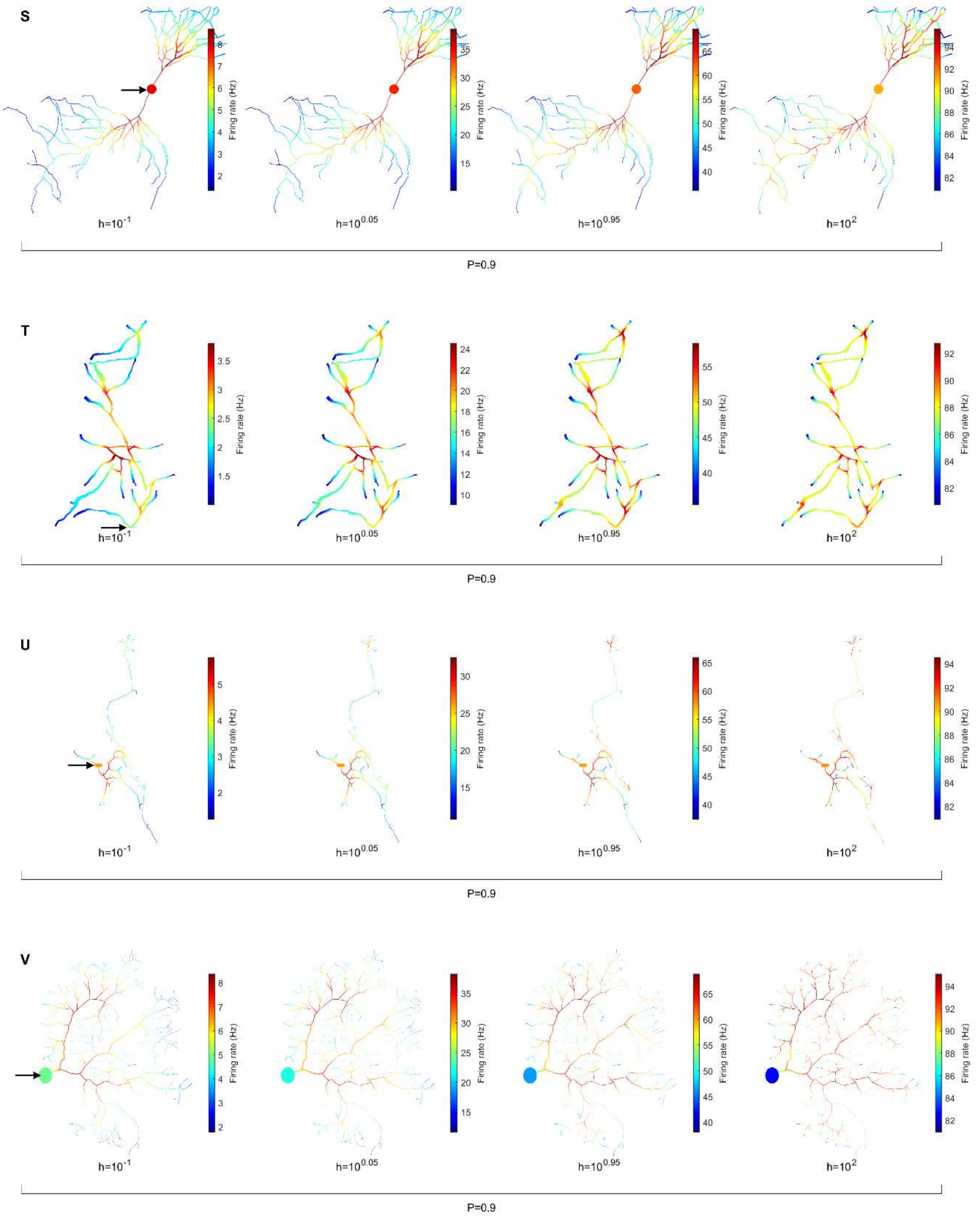

S
h=$10^{-1}$
h=$10^{0.05}$
h=$10^{0.95}$
h=$10^{2}$
P=0.9
T
h=$10^{-1}$
h=$10^{0.05}$
h=$10^{0.95}$
h=$10^{2}$
P=0.9
U
h=$10^{-1}$
h=$10^{0.05}$
h=$10^{0.95}$
h=$10^{2}$
P=0.9
V
h=$10^{-1}$
h=$10^{0.05}$
h=$10^{0.95}$
h=$10^{2}$
P=0.9
Firing rate (Hz)

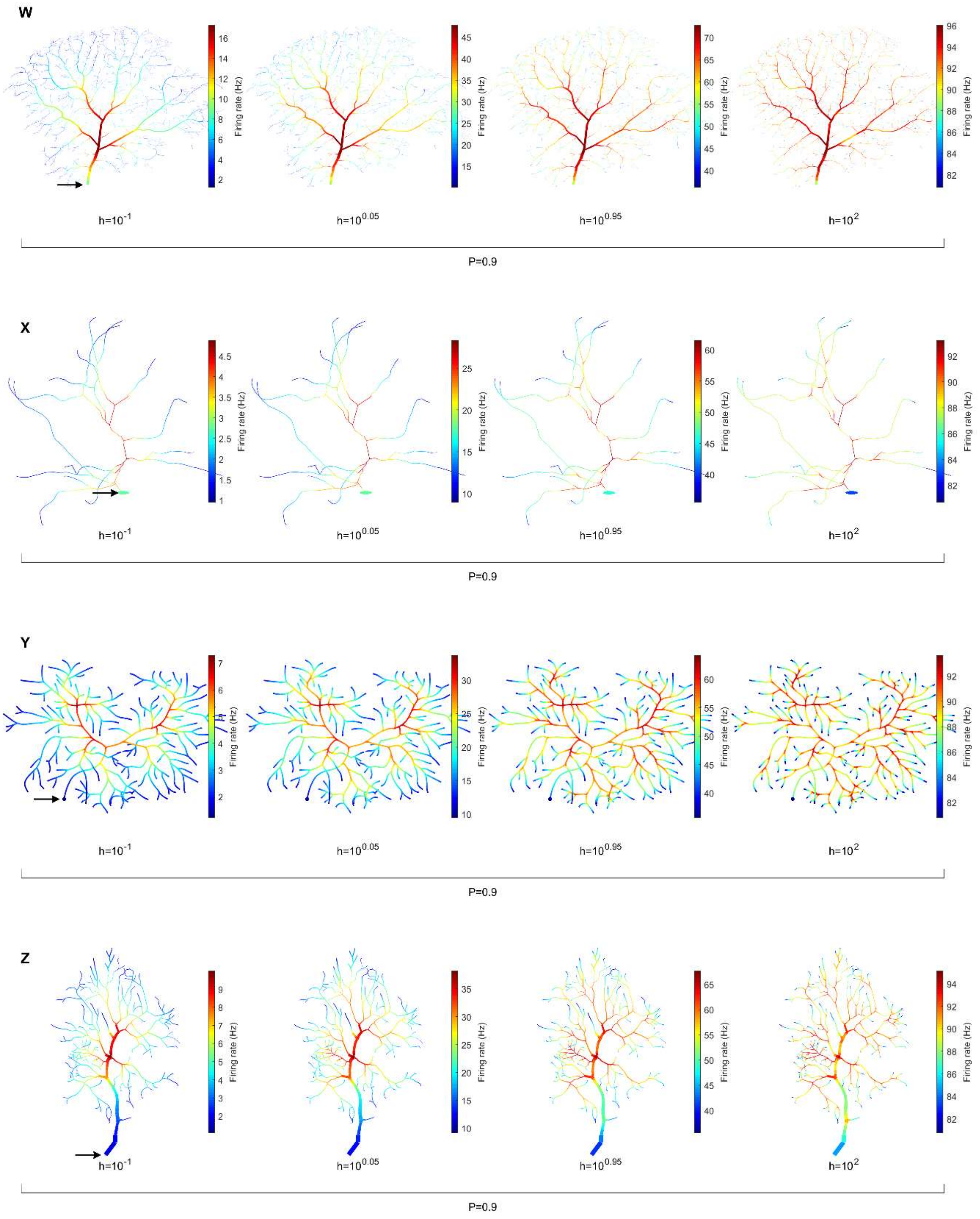

W
Firing rate (Hz)
2 4 6 8 10 12 14 16
15 20 25 30 35 40 45
40 45 50 55 60 65 70
82 84 86 88 90 92 94 96
h=10^-1
h=10^0.05
h=10^0.95
h=10^2
P=0.9
X
Firing rate (Hz)
1 1.5 2 2.5 3 3.5 4 4.5
10 15 20 25
40 45 50 55 60
82 84 86 88 90 92
h=10^-1
h=10^0.05
h=10^0.95
h=10^2
P=0.9
Y
Firing rate (Hz)
2 3 4 5 6 7
10 15 20 25 30
40 45 50 55 60
82 84 86 88 90 92
h=10^-1
h=10^0.05
h=10^0.95
h=10^2
P=0.9
Z
Firing rate (Hz)
2 3 4 5 6 7 8 9
10 15 20 25 30 35
40 45 50 55 60 65
82 84 86 88 90 92 94
h=10^-1
h=10^0.05
h=10^0.95
h=10^2
P=0.9

### Figure S3. Dynamic range heat maps of all neurons varied over P.

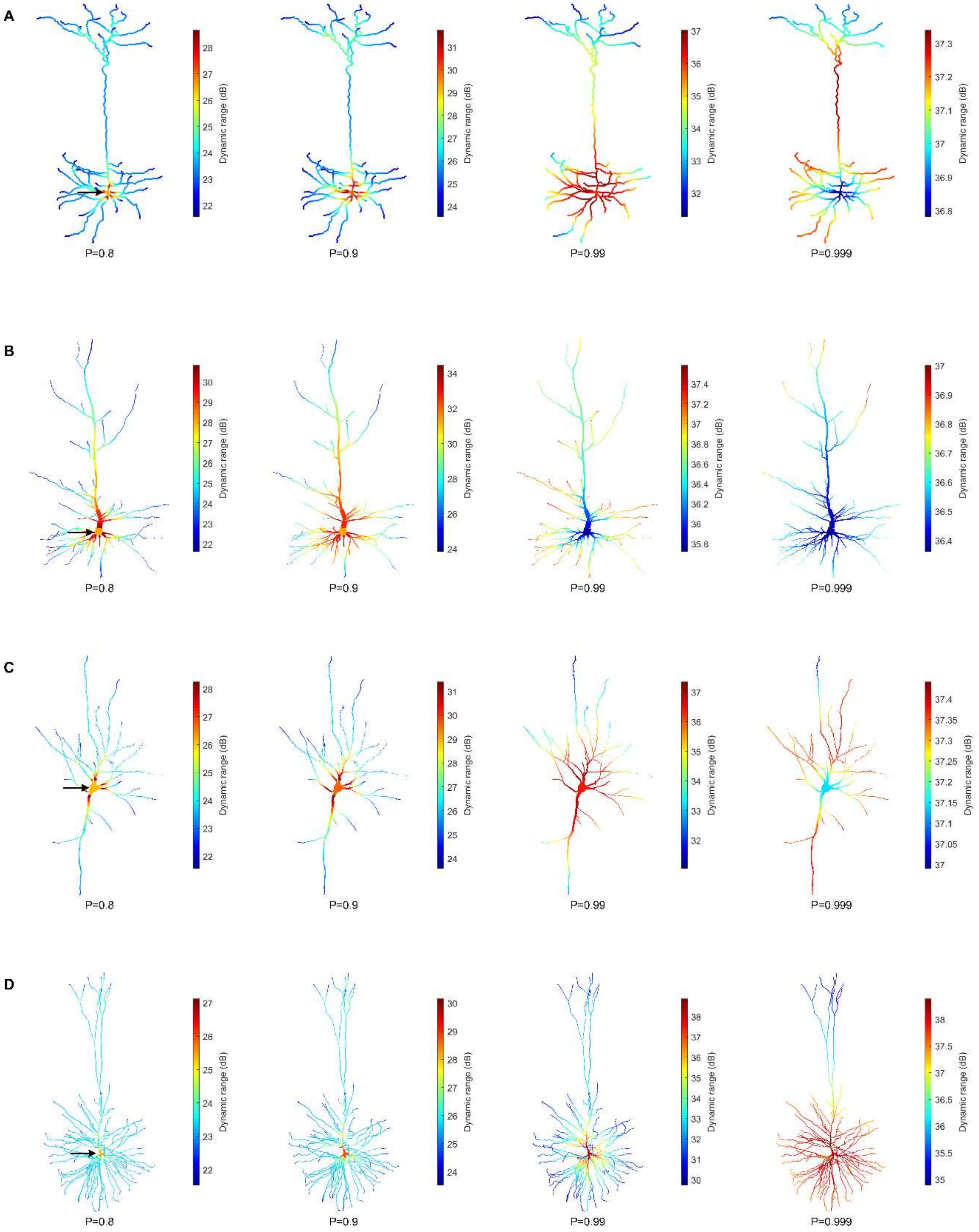

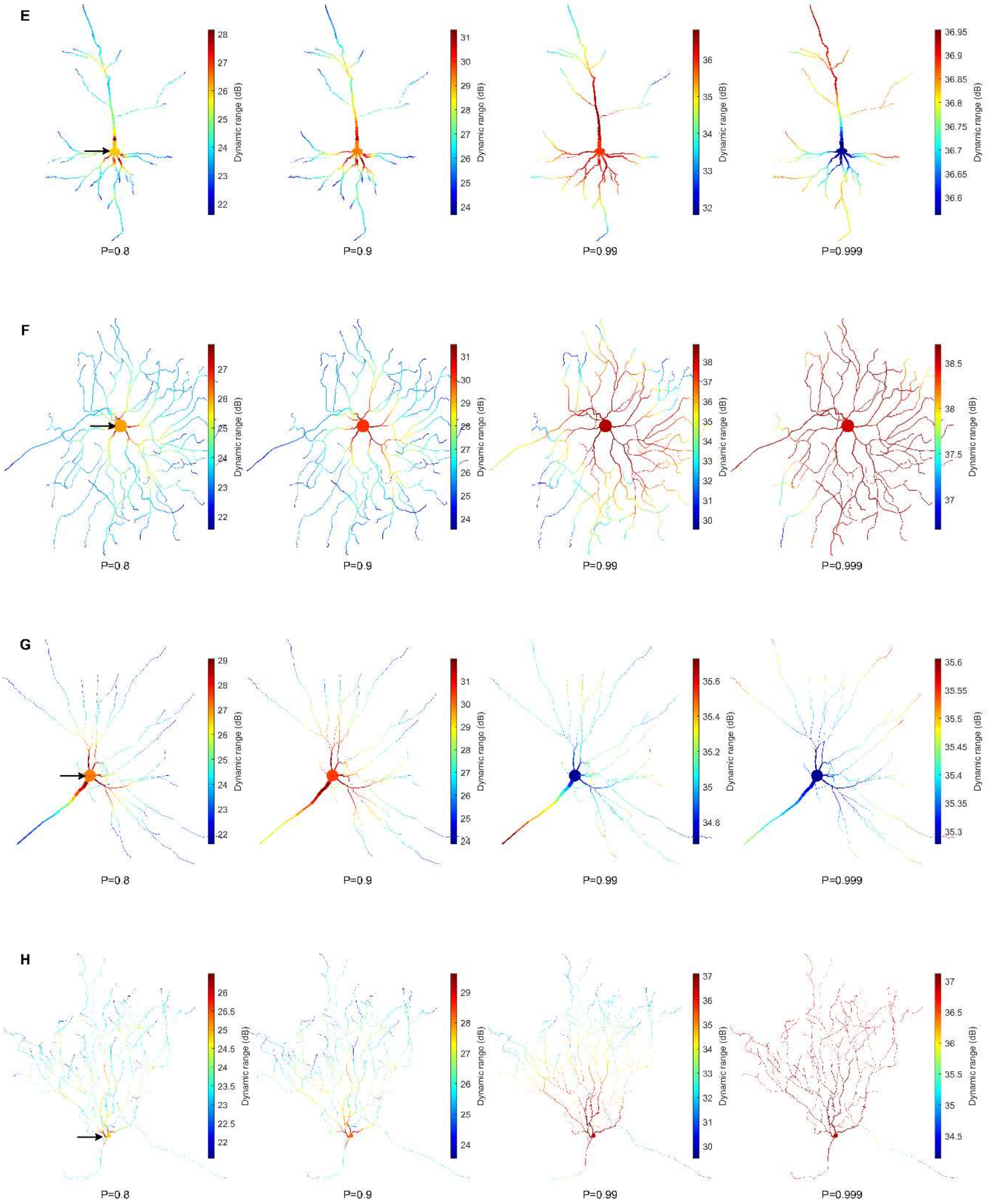

E
Dynamic range (dB)
28 27 26 25 24 23 22
P=0.8
31 30 29 28 27 26 25 24
P=0.9
36 35 34 33 32
P=0.99
36.95 36.9 36.85 36.8 36.75 36.7 36.65 36.6
P=0.999
F
27 26 25 24 23 22
P=0.8
31 30 29 28 27 26 25 24
P=0.9
38 37 36 35 34 33 32 31 30
P=0.99
38.5 38 37.5 37
P=0.999
G
29 28 27 26 25 24 23 22
P=0.8
31 30 29 28 27 26 25 24
P=0.9
35.6 35.4 35.2 35 34.8
P=0.99
35.6 35.55 35.5 35.45 35.4 35.35 35.3
P=0.999
H
26 25.5 25 24.5 24 23.5 23 22.5 22
P=0.8
29 28 27 26 25 24
P=0.9
37 36 35 34 33 32 31 30
P=0.99
37 36.5 36 35.5 35 34.5
P=0.999

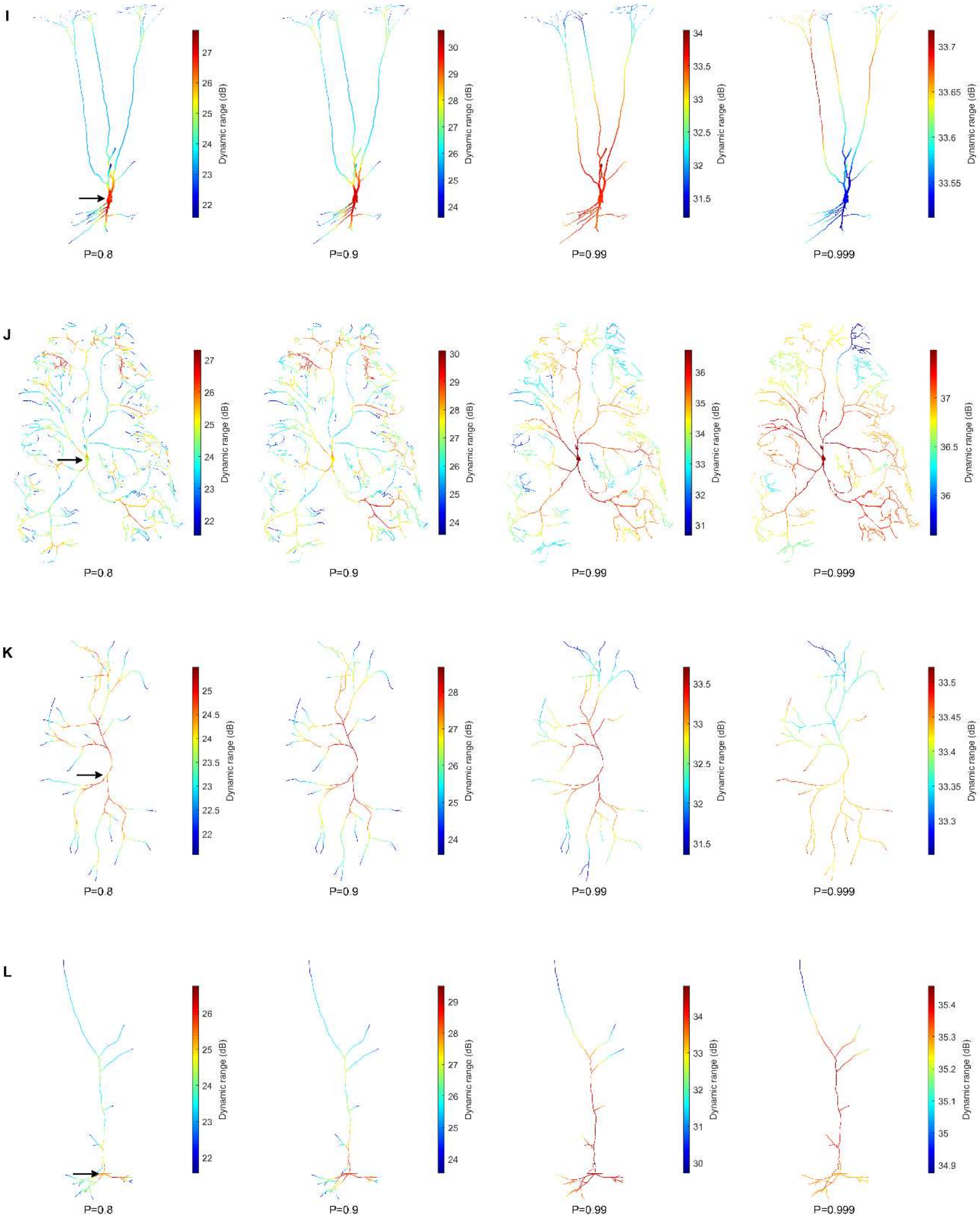
I
27
26
25
24
23
22
Dynamic range (dB)
P=0.8
30
29
28
27
26
25
24
Dynamic range (dB)
P=0.9
34
33.5
33
32.5
32
31.5
Dynamic range (dB)
P=0.99
33.7
33.65
33.6
33.55
Dynamic range (dB)
P=0.999
J
27
26
25
24
23
22
Dynamic range (dB)
P=0.8
30
29
28
27
26
25
24
Dynamic range (dB)
P=0.9
36
35
34
33
32
31
Dynamic range (dB)
P=0.99
37
36.5
36
Dynamic range (dB)
P=0.999
K
25
24.5
24
23.5
23
22.5
22
Dynamic range (dB)
P=0.8
28
27
26
25
24
Dynamic range (dB)
P=0.9
33.5
33
32.5
32
31.5
Dynamic range (dB)
P=0.99
33.5
33.45
33.4
33.35
33.3
Dynamic range (dB)
P=0.999
L
26
25
24
23
22
Dynamic range (dB)
P=0.8
29
28
27
26
25
24
Dynamic range (dB)
P=0.9
34
33
32
31
30
Dynamic range (dB)
P=0.99
35.4
35.3
35.2
35.1
35
34.9
Dynamic range (dB)
P=0.999

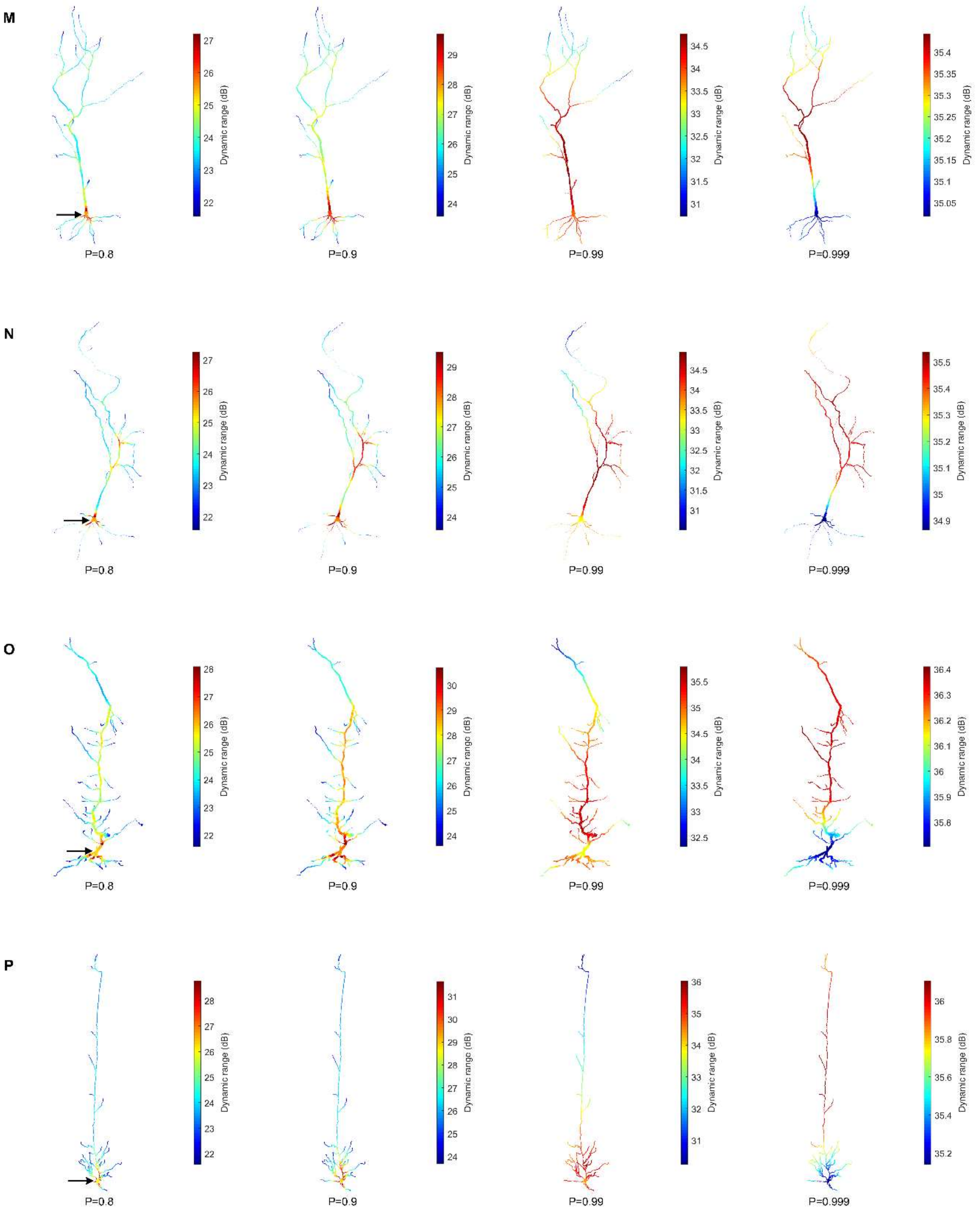
M
N
O
P
P=0.8
P=0.9
P=0.99
P=0.999
Dynamic range (dB)
27 26 25 24 23 22
29 28 27 26 25 24
34.5 34 33.5 33 32.5 32 31.5 31
35.4 35.35 35.3 35.25 35.2 35.15 35.1 35.05
35.5 35.4 35.3 35.2 35.1 35 34.9
28 27 26 25 24 23 22
30 29 28 27 26 25 24
35.5 35 34.5 34 33.5 33 32.5
36.4 36.3 36.2 36.1 36 35.9 35.8
31 30 29 28 27 26 25 24
36 35 34 33 32 31
36 35.8 35.6 35.4 35.2

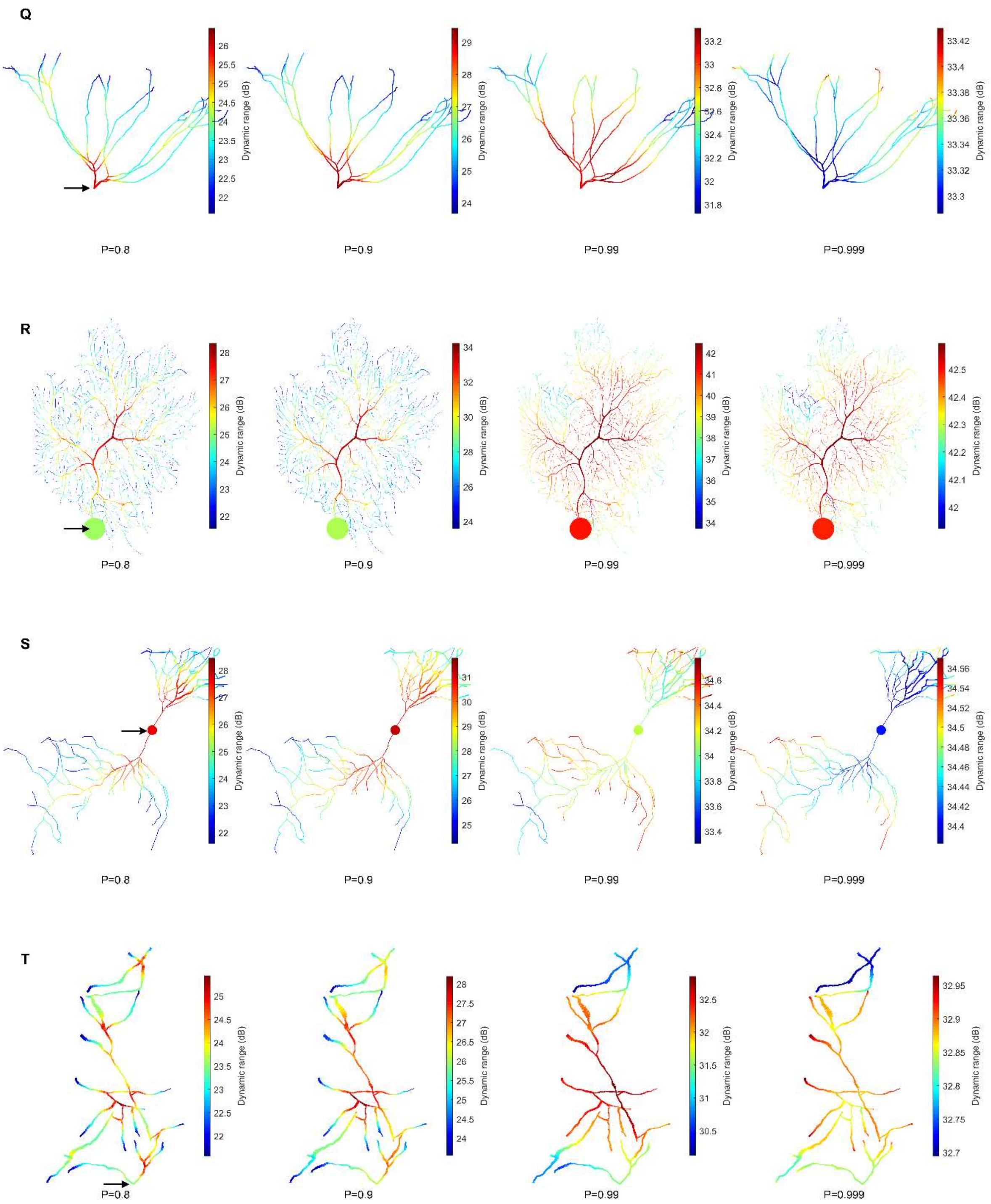
Q
Dynamic range (dB)
P=0.8
P=0.9
P=0.99
P=0.999
R
Dynamic range (dB)
P=0.8
P=0.9
P=0.99
P=0.999
S
Dynamic range (dB)
P=0.8
P=0.9
P=0.99
P=0.999
T
Dynamic range (dB)
P=0.8
P=0.9
P=0.99
P=0.999

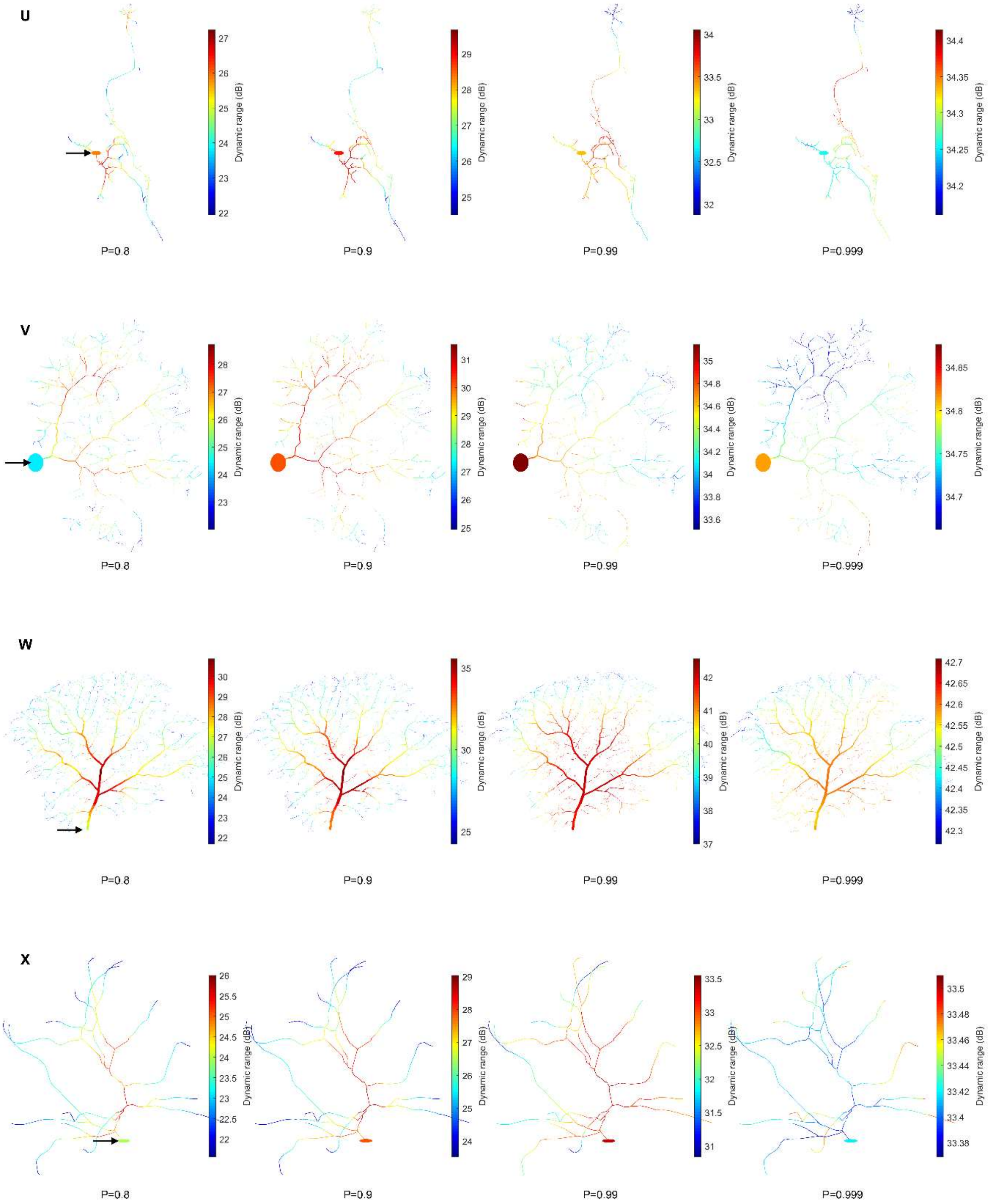
U
V
W
X
P=0.8
P=0.9
P=0.99
P=0.999
Dynamic range (dB)

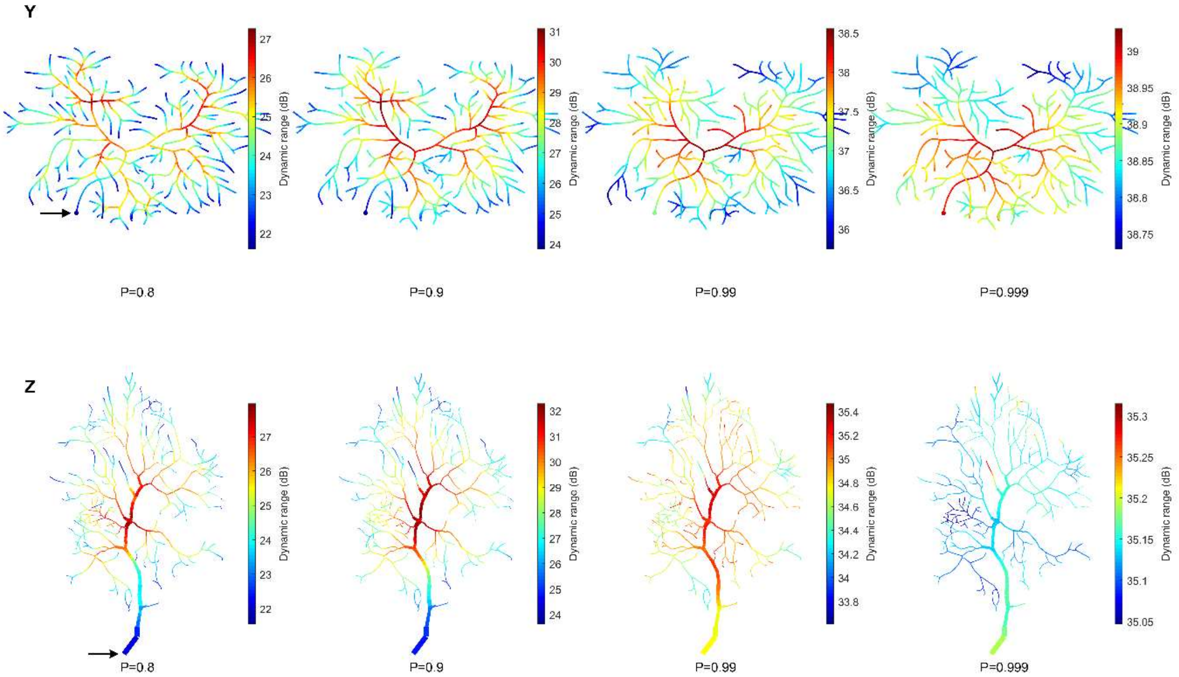


### Figure S4. Compartmental dynamic range as a function of distance from soma for all neurons.

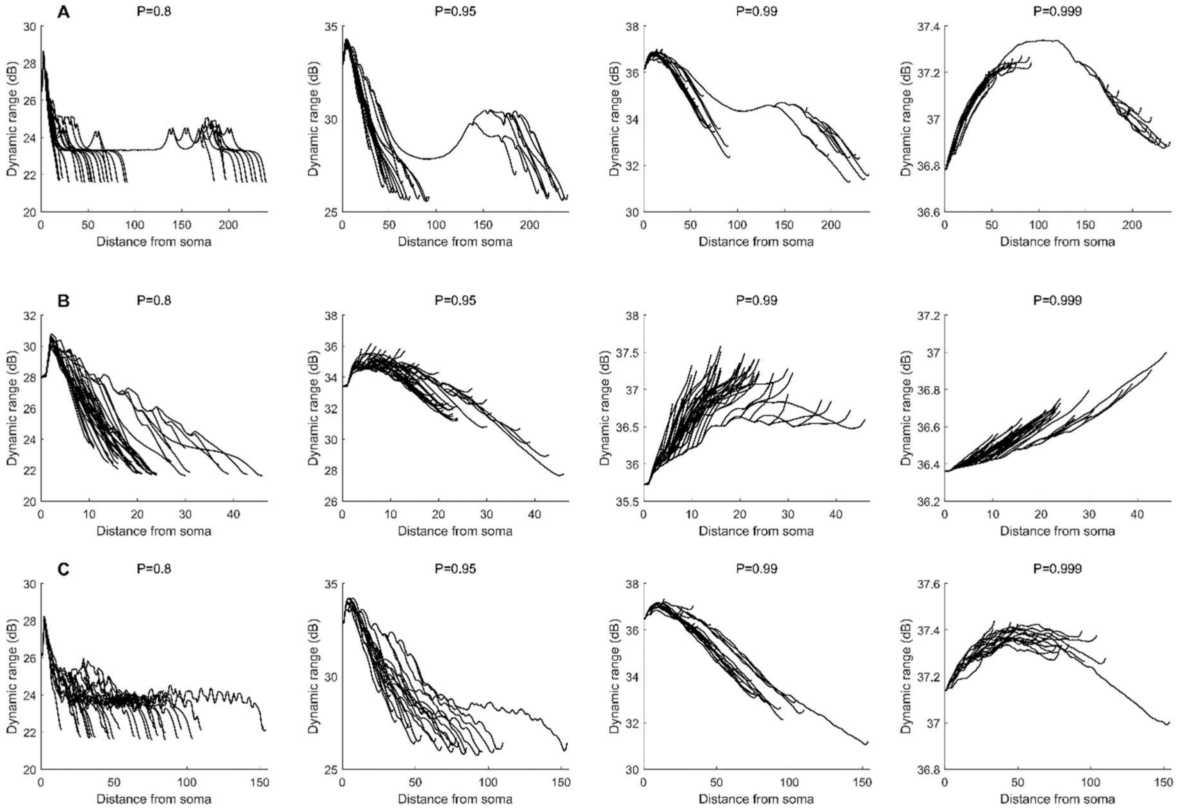

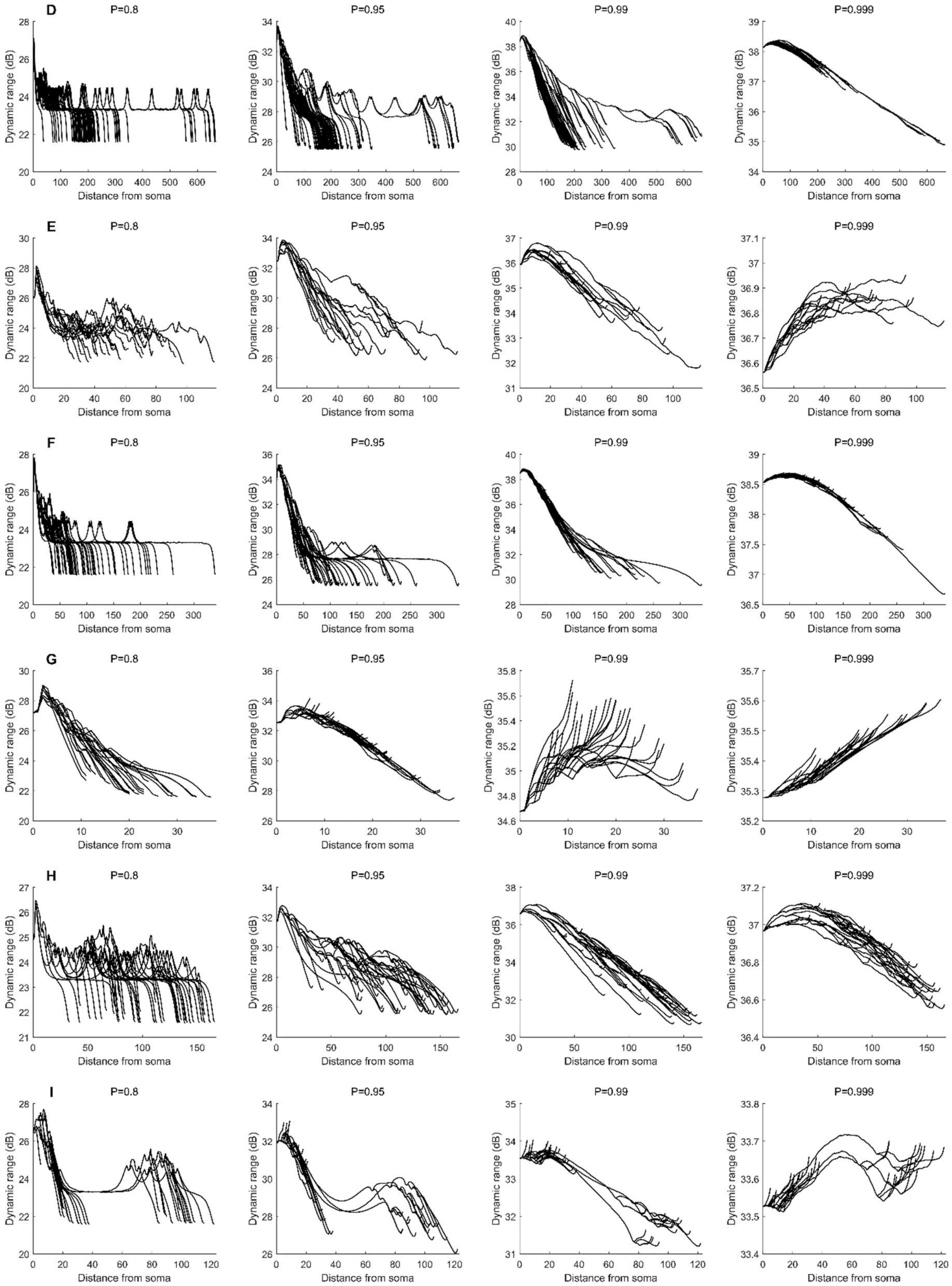
D
P=0.8
P=0.95
P=0.99
P=0.999
E
P=0.8
P=0.95
P=0.99
P=0.999
F
P=0.8
P=0.95
P=0.99
P=0.999
G
P=0.8
P=0.95
P=0.99
P=0.999
H
P=0.8
P=0.95
P=0.99
P=0.999
I
P=0.8
P=0.95
P=0.99
P=0.999
Dynamic range (dB)
Distance from soma

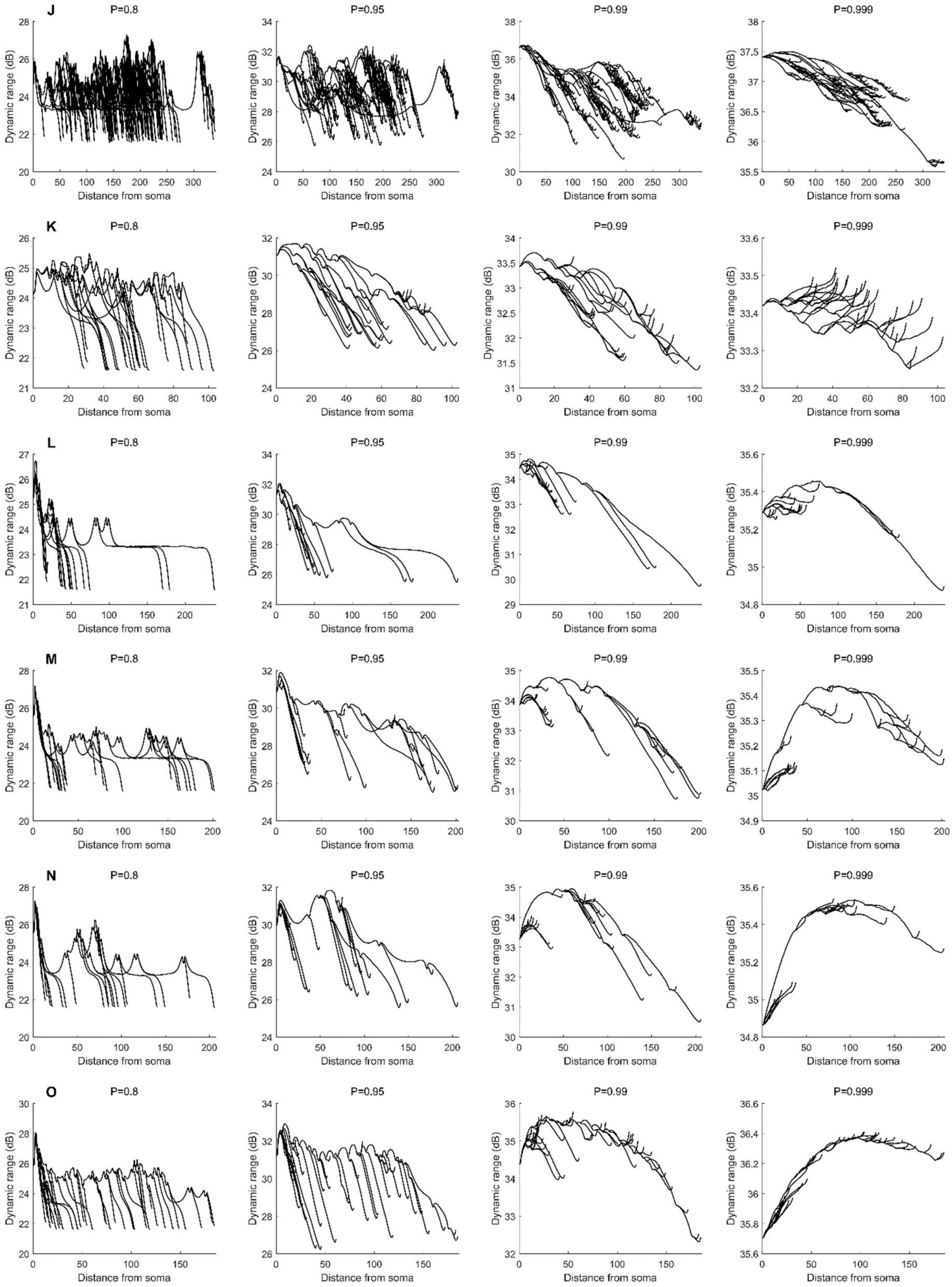

J
K
L
M
N
O
P=0.8
P=0.95
P=0.99
P=0.999
Dynamic range (dB)
Distance from soma

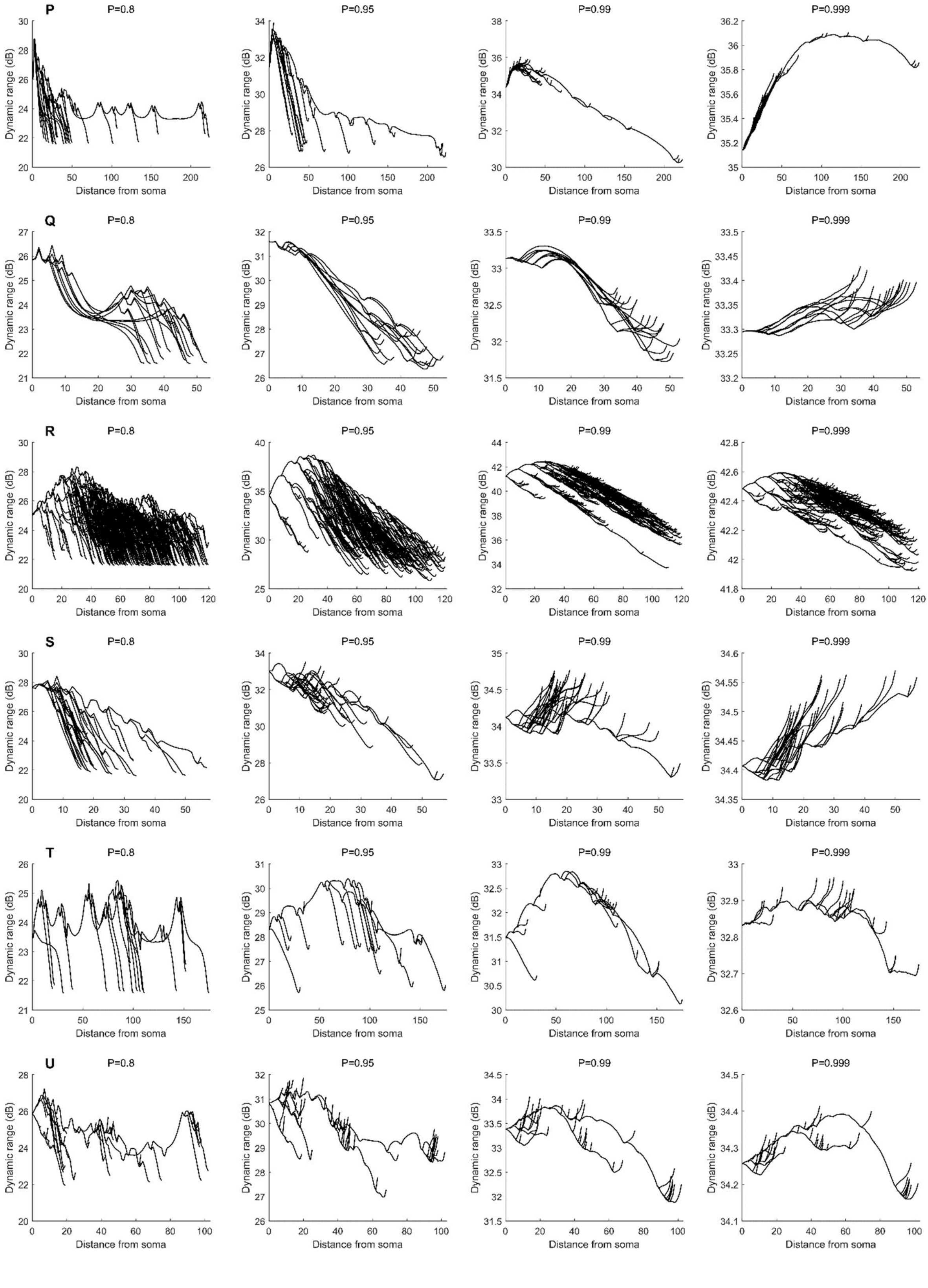
P
P=0.8
P=0.95
P=0.99
P=0.999
Q
P=0.8
P=0.95
P=0.99
P=0.999
R
P=0.8
P=0.95
P=0.99
P=0.999
S
P=0.8
P=0.95
P=0.99
P=0.999
T
P=0.8
P=0.95
P=0.99
P=0.999
U
P=0.8
P=0.95
P=0.99
P=0.999
Dynamic range (dB)
Distance from soma

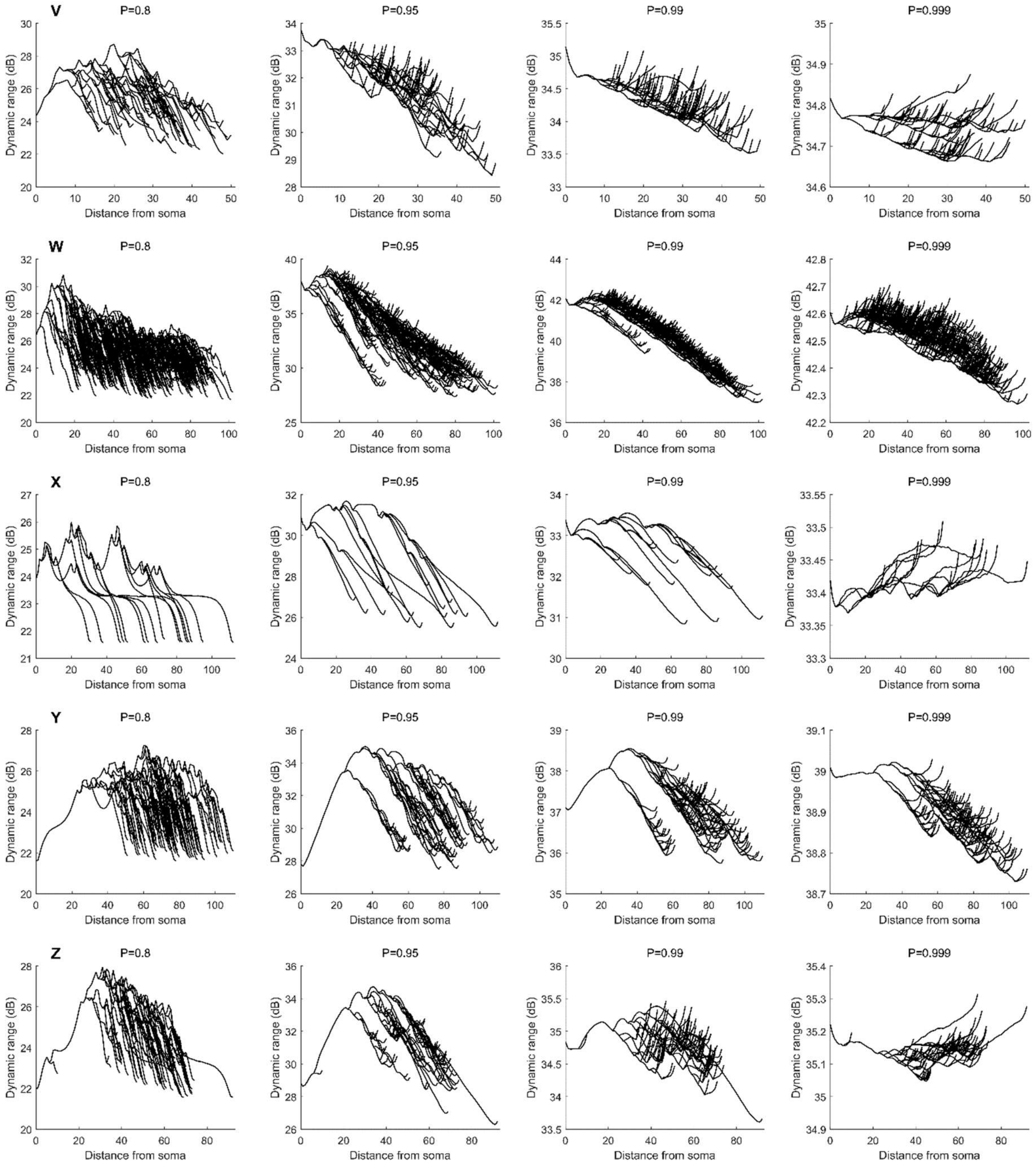

V
W
X
Y
Z
P=0.8
P=0.95
P=0.99
P=0.999
Dynamic range (dB)
Distance from soma

### Figure S5. Additional dynamics of benchmark neurites.

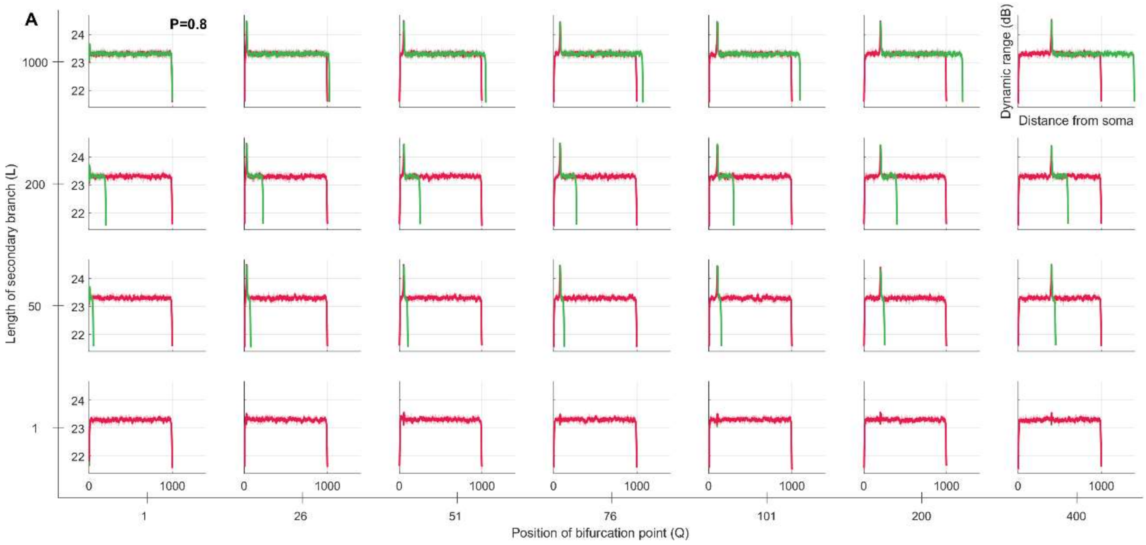

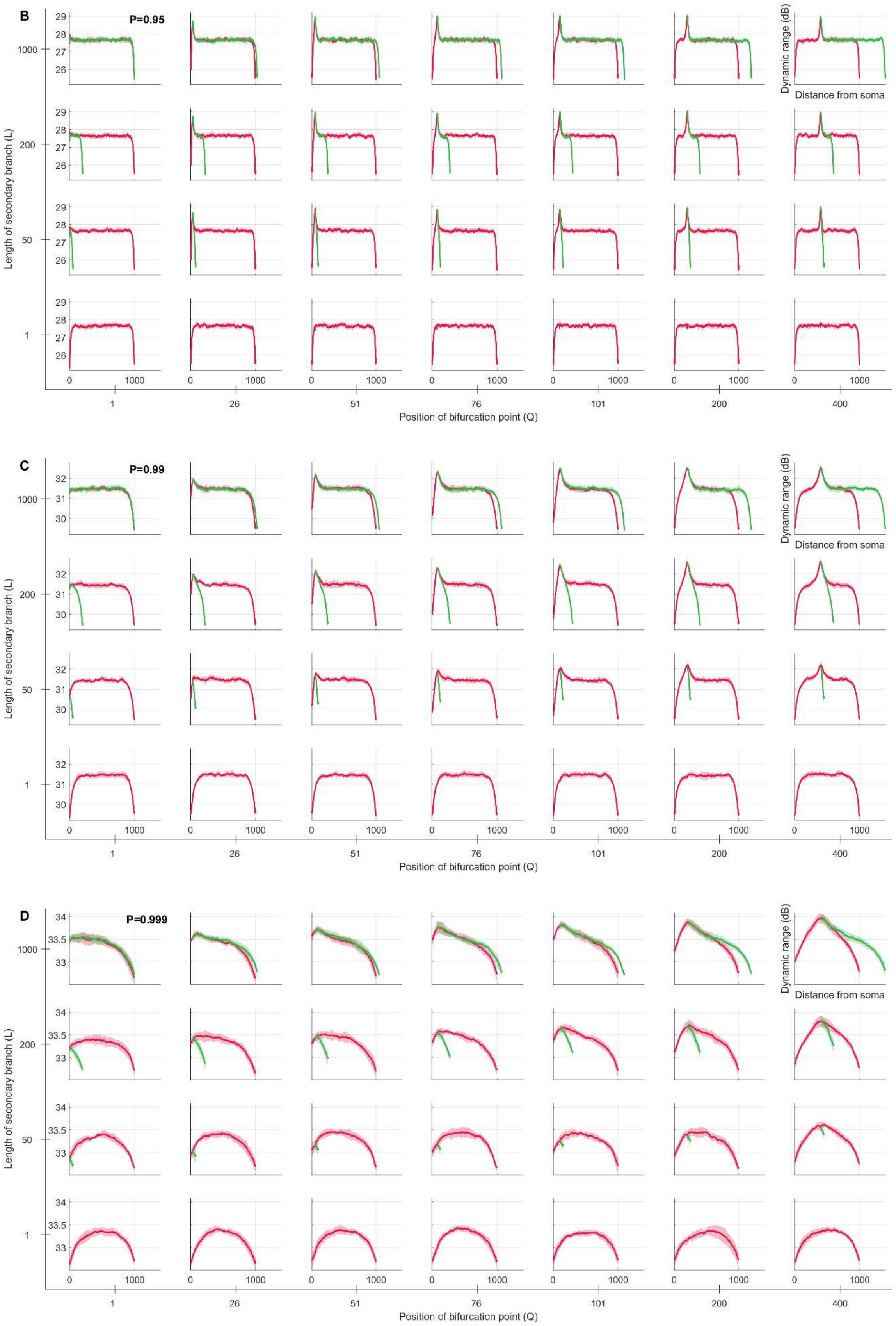
B
P=0.95
Dynamic range (dB)
Distance from soma
Length of secondary branch (L)
1000
200
50
1
29
28
27
26
0
1000
1
26
51
76
101
200
400
Position of bifurcation point (Q)
C
P=0.99
32
31
30
D
P=0.999
34
33.5
33

### Figure S6. Relative energy consumption plots of all neurons.

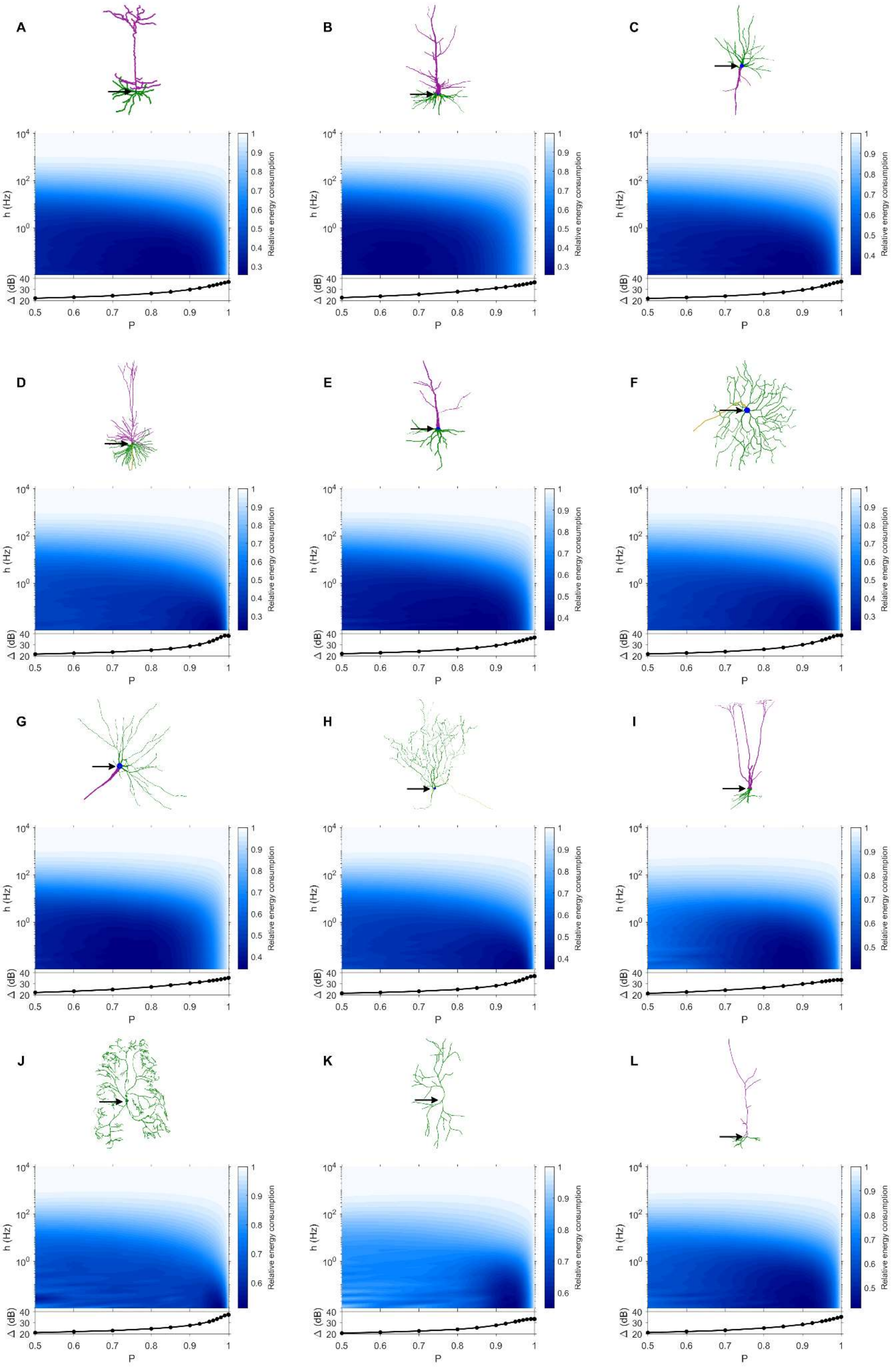

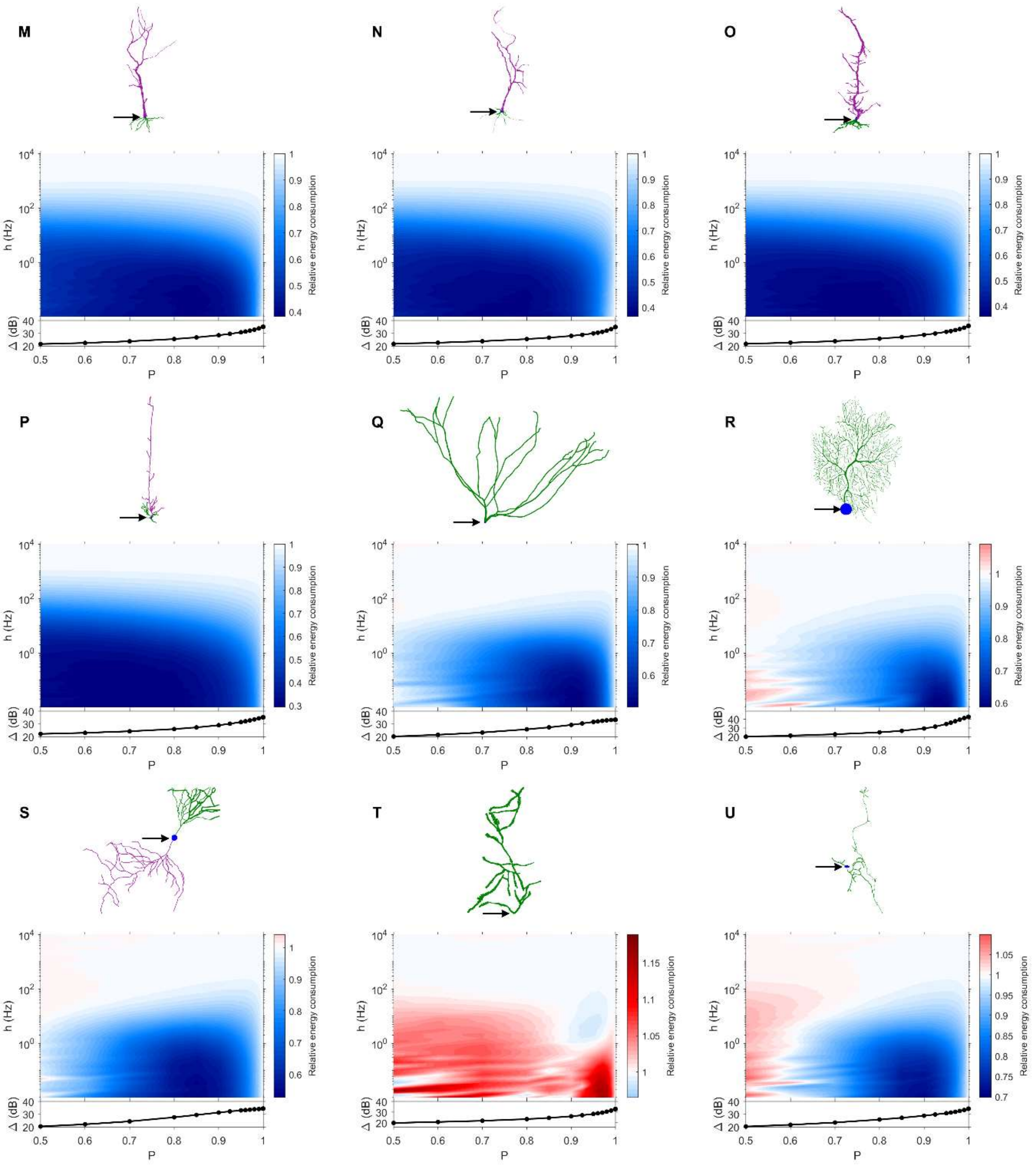

M
N
O
P
Q
R
S
T
U
h (Hz)
Δ (dB)
P
Relative energy consumption

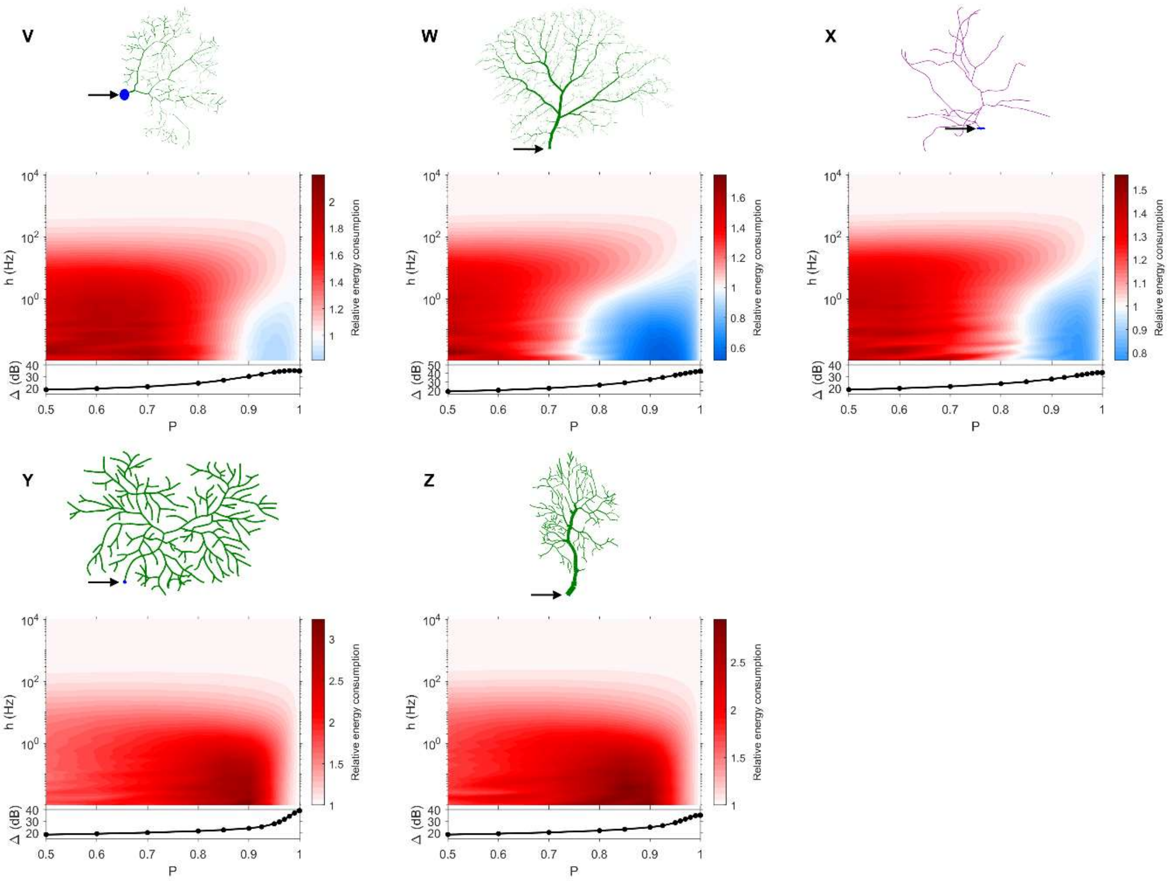


### Figure S7. Relative centrality heat maps for all neurons.

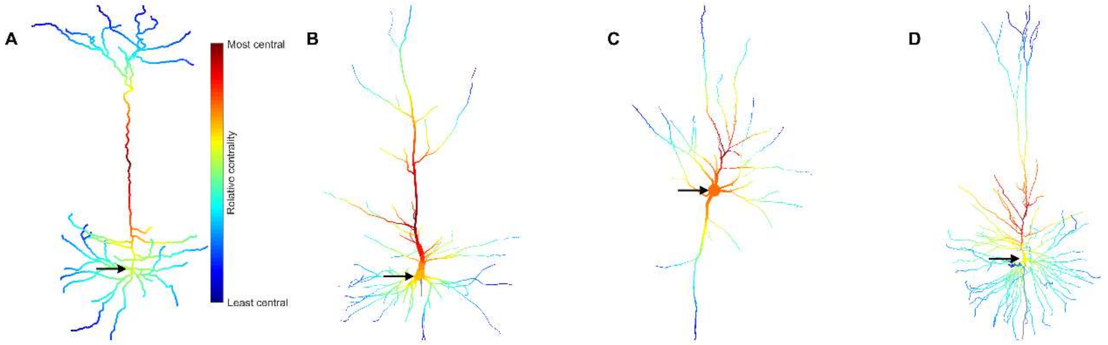

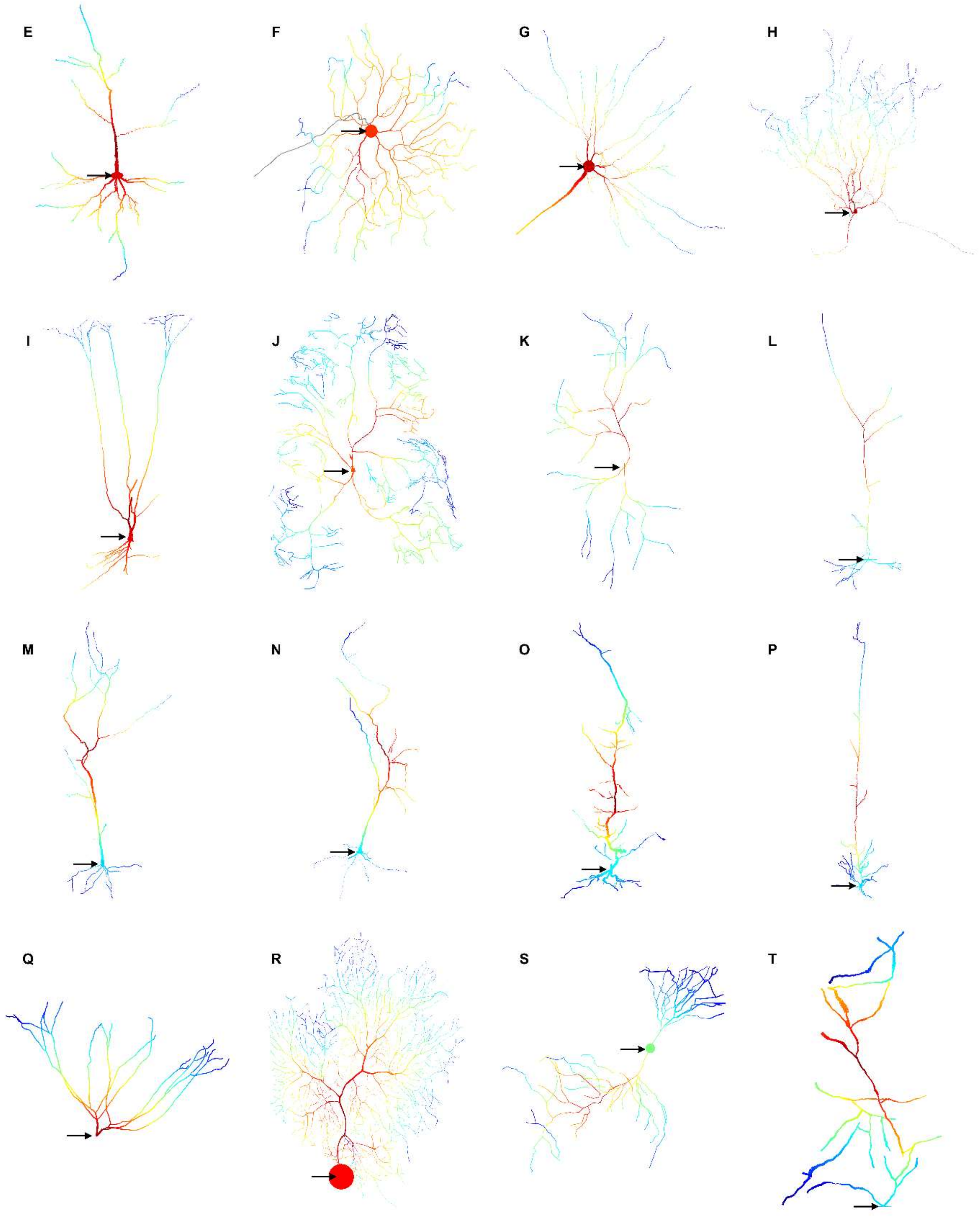
E
F
G
H
I
J
K
L
M
N
O
P
Q
R
S
T

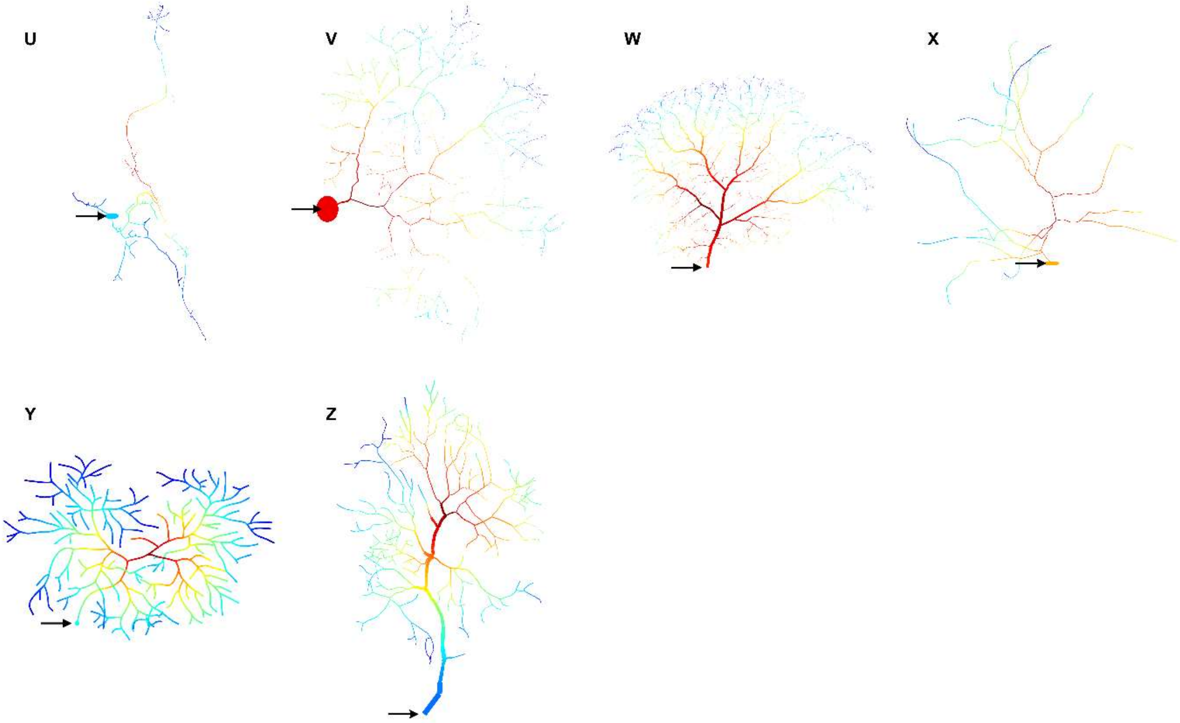
U
V
W
X
Y
Z

Figure S8. Correlation between centrality and dynamic range.

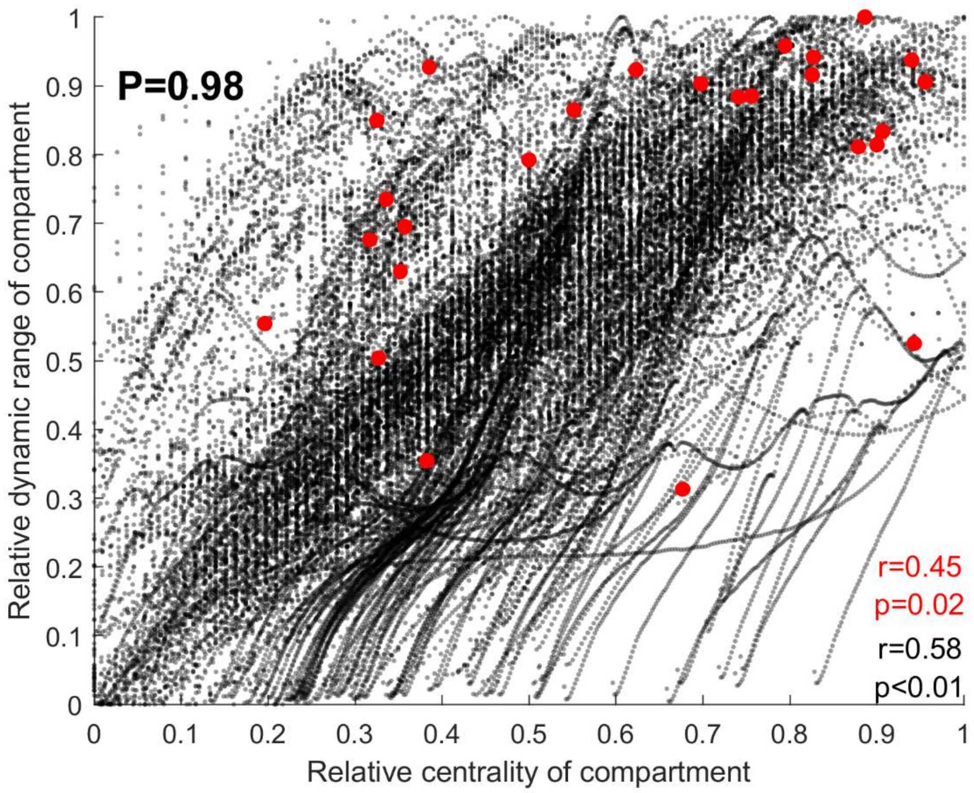


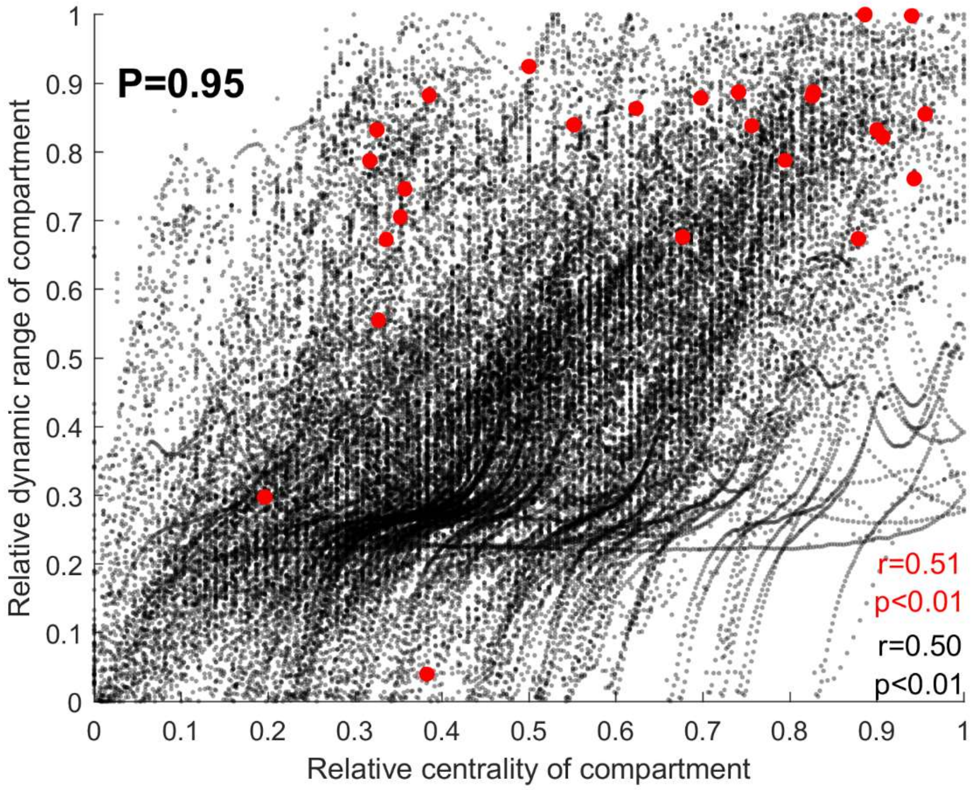
